# Phylogeny-agnostic strain-level prediction of phage-host interactions from genomes

**DOI:** 10.1101/2025.11.15.688630

**Authors:** Avery J. C. Noonan, Lucas Moriniere, Edwin O. Rivera-López, Krish Patel, Melina Pena, Madeline Svab, Alexey Kazakov, Adam Deutschbauer, Edward G. Dudley, Vivek K. Mutalik, Adam P. Arkin

## Abstract

Bacteriophages offer promising alternatives to antibiotics for treating drug-resistant infections and engineering microbiomes, but applications are limited by inability to select phages infecting specific bacterial strains. Selecting suitable phages requires either one-to-one experimental assays or strain-level predictions of phage-host interactions. Existing computational approaches either predict host taxonomy at broad ranks unsuitable for strain-level targeting or require species-specific mechanistic knowledge limiting generalizability. Here, we present a phylogenyagnostic machine learning framework predicting strain-level phage-host interactions across diverse bacterial genera from genome sequences alone. Systematically optimizing the workflow over 13.2 million training runs across six datasets (115,037 interactions, 949 bacterial strains, 518 phages), we achieved performance matching species-specific methods (AUROC 0.67-0.94) while eliminating phylogenetic constraints. Comprehensive feature engineering identifies biologically interpretable genetic determinants while minimizing overfitting in sparse, imbalanced datasets. Experimental validation through 1,240 novel interactions confirmed generalizability (AUROC 0.84), while genome-wide RB-TnSeq screens verified that 68.6% of experimentally identified infection mediators were captured computationally, including receptors and cell wall biosynthesis pathways. Model-guided cocktail design achieved up to 97.5% bacterial coverage with five phages, and up to a 3.1-fold improvement in single-phage selection over promiscuity-based selection. This platform enables rational phage therapy design and precision microbiome engineering with applications in combating antimicrobial resistance across clinical, agricultural, and industrial contexts.

## Introduction

Bacteriophage host specificity is the byproduct of an evolutionary arms race, where phages continuously respond to evolving host receptors, cell wall structures and defense mechanisms^1–7^. In the context of phage therapies, this specific, multivalent ‘recognition’ system enables the targeting of individual bacterial strains while minimizing collateral impact on surrounding microbiota. This offers significant advantages over traditional broad-spectrum antibiotics in the treatment of bacterial infections and contamination^8,9^, representing a promising frontier in microbiome engineering, with applications spanning therapeutics, agriculture, and industrial microbiology. Phages could provide an alternative treatment option for antibiotic-resistant infections, which the World Health Organization has identified as one of the most pressing global health threats^10^. They may also be effective tools for basic science, providing precision targeting for microbiome editing to dissect mechanisms of microbial ecological interaction and function *in situ*^11^.

The clinical and biotechnological application of phages depends on our ability to accurately select phages capable of infecting specific bacterial strains^8,9,12,13^. Although their narrow host-specificity relative to antibiotics can reduce the impact of treatment on beneficial commensal bacteria, this also presents a challenge in identifying suitable therapeutic phages for specific targets^14^. As the size of phage banks increases to capture the breadth of bacterial hosts, it becomes less feasible to test phage susceptibility experimentally for a new target pathogen or contaminant^14,15^. In situations where pathogens are not susceptible to broad host-range cocktails, predictive models offer strategies to quickly identify candidate phages across available phage banks from host genetic information^12,13^, enabling clinicians to focus efforts on high-probability phages. However, because of the complex, multifactorial mediators of phage-host interactions, robust predictive modeling of these interactions at strain-level requires high-quality interaction datasets capturing a range of representative host and phage phylogenies and their associated phage infection mechanisms^13,16,17^.

Unlike metagenomic host-classification tools, which operate at family or genus level and cannot resolve individual strain-phage interactions^16–37^, predicting phage-host interactions at single-strain granularity requires datasets of confirmed one-to-one interactions, documenting both successful and unsuccessful infections, of sufficient size and with appropriate variation of genetic components^16,17^. These datasets are time-consuming to generate, often relying on plaquing assays or microplate-based liquid assays, as well as isolation and sequencing of individual bacterial strain and phage genomes^9,16,17,38^. These experiments have revealed that interaction datasets can be highly imbalanced, with only a small fraction of interactions resulting in infection, and significant variability in susceptibility even within a single host species^38,39^. With as little as 2-5% of interactions resulting in infection in published *Klebsiella* and *Vibrionaceae* interaction matrices^38,40,41^, this further increases the scale of experimental efforts required to generate sufficient representations of infectious interactions for model generalizability.

Individual labs often generate only a limited amount of data about a target host and phage, while focusing on specific host phylogenies^6,16,38,40–43^. Consequently, there are few published datasets of sufficient size for predictive modeling, and relatively few tools have been developed to specifically predict interactions between unrelated strain-phage pairs^13,16,17^, with none of these tools supporting phylogeny-independent prediction of interactions between novel bacterial strains and phages. These data limitations complicate predictive modeling through experimental heterogeneity across laboratories, lineage-specific variation that limits cross-phylogeny transfer learning, and uneven phylogenetic sampling within species. It would therefore be advantageous to establish workflows that leverage and combine community-generated datasets across diverse host and phage phylogenies, increasing statistical power while capturing shared mechanisms of phage-host interaction.

Machine-learning (ML)-based predictive modeling of phage-host interactions requires the representation of training data as numerical feature tables associated with a phenotype. Converting complex genomic data into numerical values results in high-dimensionality feature tables, presenting a risk of overfitting when training on small and imbalanced phenotypic datasets^13,44^. Many existing tools for strain-level phage-host interaction prediction address these challenges through reduction of feature-set dimensionality by selecting genetic features known to mediate these interactions^16,17,45^. Although effective, this strategy limits the applicability of these workflows across diverse phage and host phylogenies and hinders the identification of novel features mediating their interactions^16,17^. Some approaches also employ dimension-reduction steps that impede biological interpretability^16^. In the context of phage-host interaction prediction, many existing workflows rely on phage-specific models, which restricts predictions to whether unseen bacterial strains are infected by an existing set of phages^13,16^. Collectively, these constraints limit the applicability of existing models in combating diverse emerging pathogens and contaminants.

To address these limitations, we leveraged 6 published phage-host interaction datasets totaling 115,037 interactions, across 949 bacterial strains and 518 phages^16,38,40–42^ to develop a predictive modeling framework that incorporates versatile feature assignment, comprehensive feature selection, and ensemble-learning-based modeling, parameterized for the characteristics of phage-host interaction data and other sparse microbial phenotypic data. Bacterial and phage genomes are represented as pangenome-like presence-absence matrices of protein family or amino acid *k*-mer content, enabling phylogeny-agnostic predictions for novel strain-phage pairs without requiring prior mechanistic knowledge. Filtering of phylogenetically-linked features minimizes the identification of clade signatures rather than functionally relevant mechanisms, enabling biological interpretation of predictions and identification of both known and potentially novel determinants of phage-host interactions.

Here, we show that our approach can match or improve upon the performance of previous models trained on published datasets, without prior assumptions of important genetic features; demonstrate predictive capacity on new hosts; demonstrate how models can guide phage cocktail design for previously unseen bacterial strains; and reveal predictive features that highlight proteins responsible for key specificity mechanisms between phage and host, which could serve as targets for future engineering. Finally, we validate model performance experimentally with a 1,240-interaction phage-host interaction matrix and demonstrate the biological relevance of predictive features by performing RB-TnSeq genome-wide fitness assays in the presence of a panel of phages^7,46^. Combined, this integrated approach provides a versatile framework to predict phage-host interactions at the strain level, with potential applications in precision medicine, microbiome manipulation, and biotechnology.

## Results

### Dataset overview

We used six previously published phage-host interaction datasets, totaling 115,037 interactions across 949 bacterial strains and 518 phages to develop and validate a phylogeny-agnostic phage-host interaction prediction workflow (*Fig. 1* / *Supp. Fig. 1-8* / *Table 1*). A detailed description of these datasets and their use in modeling workflow development and validation can be found in the Supplementary Materials (*Supp. Materials 1.1.1*).

**Figure 1.**
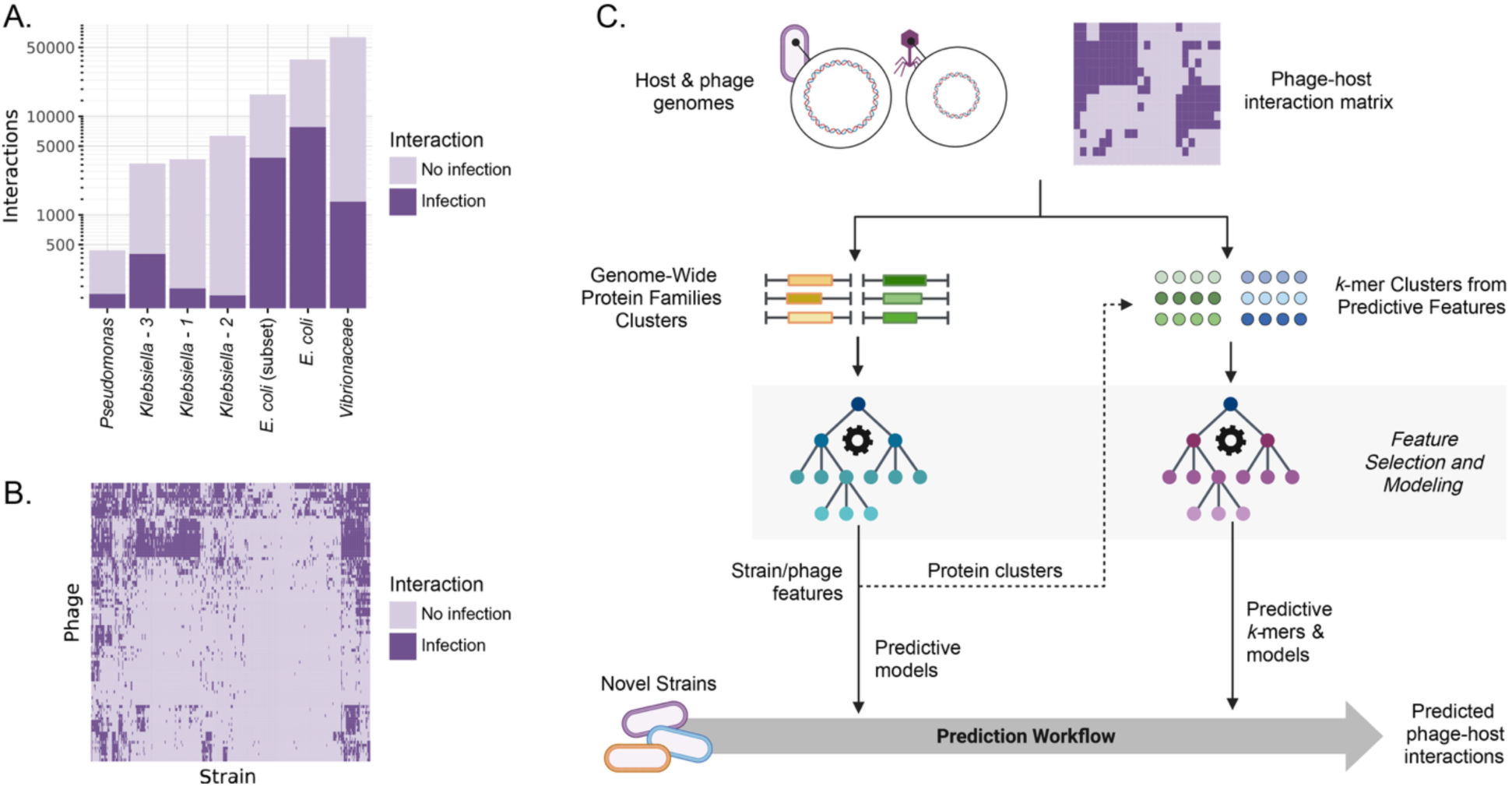
Phage-host interaction dataset and modeling workflow overview. A) Interaction datasets range from 437 to 63,488 total interactions, with proportions of infectious interactions ranging from 2.1% to 36.2%. B) The interaction matrix from the *E. coli* dataset includes 402 *E. coli* strains and 94 phages, with 20.7% of interactions resulting in infection. Clusters of bacterial strains and phages indicate conserved infection patterns within these groups. Dark purple represents interactions resulting in infection and light purple indicates interactions that did not result in infection. C) The predictive modeling workflow uses phage and bacterial genomes and phage-host interaction matrices to generate protein-family-based or *k*-mer-based feature tables for feature selection and predictive modeling. Amino acid sequences from predictive proteins can also be used to generate *k*-mer features for downstream modeling. Protein-family-based and *k-*mer-based features can be assigned to new bacterial strains and phages to predict novel interactions or can be used to explore biological mechanisms mediating phage-host interactions.

**Table 1.** Phage-host interaction dataset overview.

| Dataset ID | Host Taxonomy | Host Strains | Phages | Total Interactions | Positive Interactions | Positive Interactions (%) | Ref. |
| --- | --- | --- | --- | --- | --- | --- | --- |
| *E. coli* (subset) | *E. coli* | 402 (177) | 94 | 37,788 (16,638) | 7,813 (3,813) | 20.7 (22.9) | <sup>16</sup> |
| *Klebsiella*-1 | *Klebsiella pneumoniae* | 62 | 59 | 3,658 | 180 | 4.9 | <sup>40</sup> |
| *Klebsiella*-2 | *Klebsiella* | 138 | 46 | 6,348 | 153 | 2.4 | <sup>41</sup> |
| *Klebsiella*-3 | *Klebsiella* | 68 | 52 | 3,318 | 403 | 12.1 | <sup>47</sup> |
| *Pseudomonas* | *Pseudomonas* | 23 | 19 | 437 | 158 | 36.2 | <sup>42</sup> |
| *Vibrionaceae* | *Vibrio*<br>*Enterovibrio*<br>*Shewanella* | 256 | 248 | 63,488 | 1,364 | 2.1 | <sup>38</sup> |

### Workflow development and optimization

Our objective was to develop a modeling workflow for strain-level prediction of phage-host interactions meeting the following three key criteria: 1) predict interactions between individual bacterial strain-phage pairs; 2) be independent of host and phage phylogeny without requiring prior knowledge of genetic determinants of phage-host interaction; 3) enable predictions for novel bacterial strains or phages based on genomic information alone. Additionally, we considered secondary criteria, including biological interpretability of predictive features and the applicability of resulting predictions to phage selection and phage cocktail design.

To meet these criteria, we performed an extensive modeling parameter optimization experiment, evaluating over 200 parameter settings and training more than 13.2 million individual predictive models. This enabled optimization of multiple modeling processes, including feature engineering and selection, predictive model training, and feature assignment to novel bacterial strains and phages. This produced a robust modeling pipeline to handle the high degree of sparsity, imbalance, and feature table dimensionality characteristic of phage-host interaction and genomic datasets (*Fig. 1A-B* / *Supp. Fig. 9-20 / Supp.Table 1-2*). A detailed description of the optimization workflow, optimized parameters, and resulting architecture can be found in the Supplementary Materials (*Supp. Materials 1.1.2*).

### Model performance

Model performance was evaluated through a series of nested cross-validation experiments, across three model configurations: 1) predicting the interaction between a novel bacterial strain and a known phage, representing the most likely application, where a user selects a phage from an existing phage set (“phage bank”) that interacts with a target bacterial strain; 2) predicting the interaction between a known bacterial strain and a novel phage; 3) predicting the interaction between a novel bacterial strain and a novel phage. We tested the impact of genomic representation, dataset size and structure, and dataset combination on inter- and intra-dataset prediction (*Supp. Fig. 23-30* / *Supp. Table 3-4*).

Focusing on the first configuration, we observed no significant differences in performance based on genomic representation strategy, including protein-family and *k-*mer-based models, suggesting that all representations captured sufficient biological information to predict phage-host interactions (*Fig. 2A*). We selected protein family-based models for downstream applications, which offered the best tradeoff between model performance, computational demand, and feature interpretability (*Supp. Fig. 21-22* / *Supp. Table 5-6*). Across datasets, model performance varied from mediocre to excellent, with area under the receiver operating characteristic curve (AUROC) ranging from 0.67 to 0.94, Matthews Correlation Coefficient (MCC) from 0.13 to 0.54, and normalized area under the precision-recall curve (AUPR) from 0.07 to 0.60 (*Fig. 2B / Supp. Fig. 23 / Table 2*). For *E. coli*, our models achieved an AUROC of 0.87, closely matching the performance reported by the original study (AUROC = 0.86) despite fundamental differences in approach. Notably, the published method required phage-specific models and host features based on known *E. coli* mediators of phage infection^16^, while our approach achieved comparable performance without being constrained to well-studied host phylogenies. Performance increased significantly with dataset size, with Pearson correlation coefficients of r = 0.60 for AUROC and r = 0.47 for MCC (both p < 1×10⁻⁶) and decreasing variance, showing improved stability and reliability in larger datasets (*Supp. Fig. 24-26*). Notably, the lowest-performing dataset (*Klebsiella*-1, AUROC = 0.67) comprised only 3,658 interactions with 4.9% positive interactions, highlighting that performance at the lower end of this range reflects dataset size and sparsity constraints rather than fundamental limitations of the modeling approach.

**Figure 2.**
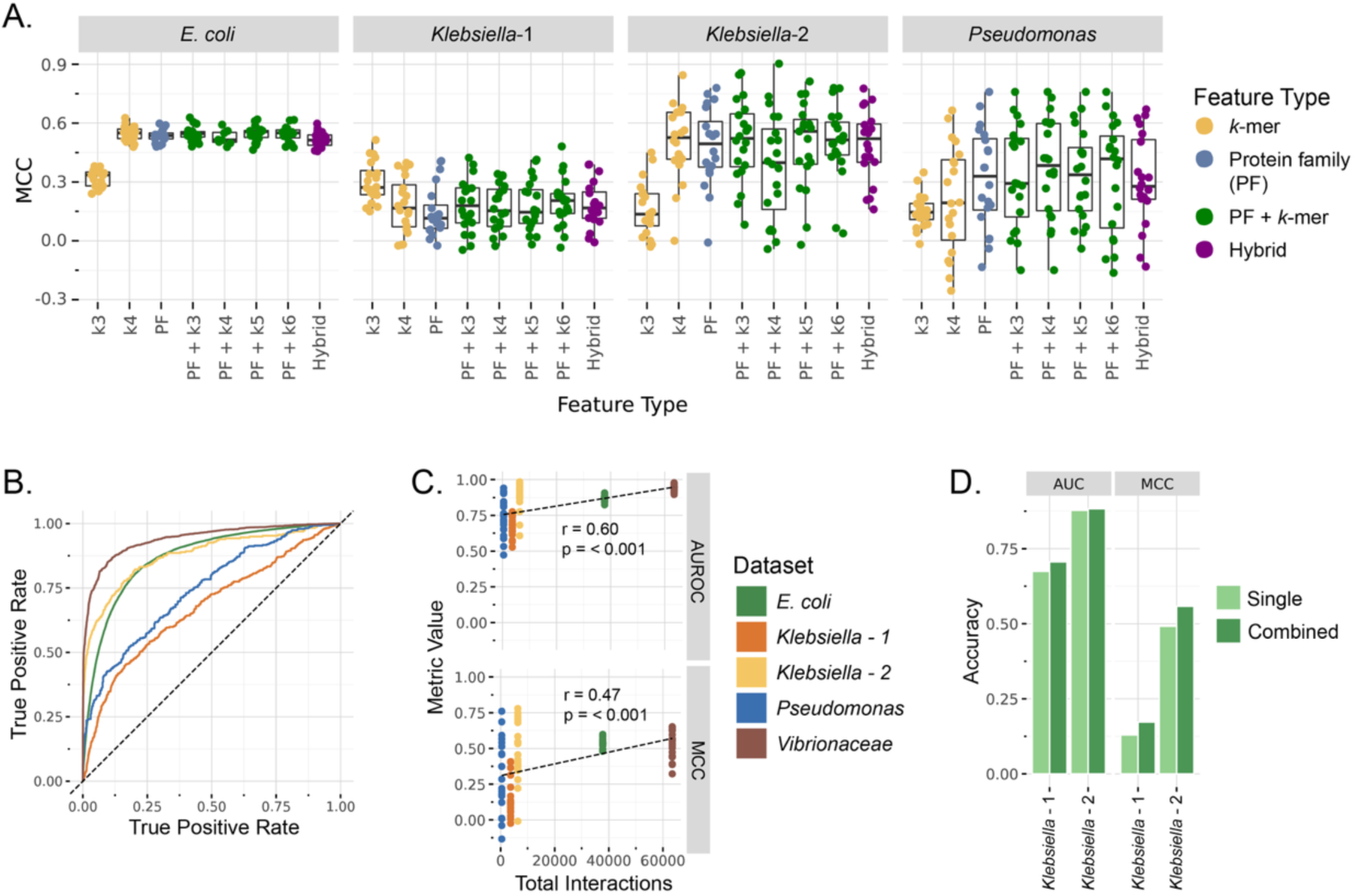
Phage-host interaction prediction performance in 20-fold cross-validation. A) Performance of models (MCC) across genome representations (*k-*mer: yellow / protein family (PF): blue / protein families with *k*-mers from predictive proteins: green / hybrid with protein families for bacterial strain and *k*-mers for phage: purple), showing no significant differences with *k* > 3. B) Receiver operating characteristic (ROC) curves show prediction performance variability across datasets, when predicting phages infecting previously unseen strains. Colors represent datasets. The dashed line indicates performance of random predictions (AUROC=0.5). C) Dataset size correlates with prediction performance. D) Combining datasets (dark green) leads to improved performance across *Klebsiella* datasets, in comparison to single-dataset models (light green).

**Table 2.** Model performance across datasets when predicting infection of unseen bacterial strains.

| Dataset | AUROC | MCC | Normalized AUPR | Brier score |
| --- | --- | --- | --- | --- |
| *Vibrionaceae* | 0.941 | 0.533 | 0.602 | 0.017 |
| *Klebsiella*-2 | 0.878 | 0.492 | 0.464 | 0.017 |
| *E. coli* | 0.869 | 0.536 | 0.562 | 0.125 |
| *Pseudomonas* | 0.745 | 0.326 | 0.504 | 0.221 |
| *Klebsiella*-1 | 0.674 | 0.130 | 0.069 | 0.061 |

Combining *Klebsiella* datasets led to modest but statistically significant performance improvements in the *Klebsiella*-2 dataset (p = 0.024) and a non-significant improvement in the *Klebsiella*-1 dataset (p = 0.398) compared to single-dataset models, with MCC increases of 0.04 and 0.07 (*Fig. 2D* / *Supp. Fig. 27*). The AUROC of 0.812 of this combined model also nearly matches the performance of models reported by a previous study (AUROC = 0.818), but whose method is also dependent on *Klebsiella*-specific knowledge, using only *K*-locus genes and phage tail fibers^17^. Unlike within-genus combinations, merging cross-genus datasets did not significantly improve performance of any dataset (*Supp. Fig. 28-29*). Leave-one-group-out (LOGO) cross-validation at the genus-level showed poor performance, with AUROC only slightly above random for all datasets, at 0.60, 0.55, and 0.57 in *E. coli*, *Klebsiella*, and *Pseudomonas* datasets, respectively (*Supp Fig. 30*). This indicates that cross-genus prediction remains challenging, likely reflecting a combination of the genus-specificity of phage-host interaction determinants and the limitations of protein families in categorizing divergent genetic features. Despite this, these models significantly enriched for phages with activity against out-of-genus strains when ranking candidates by predicted interaction probability, showing a 1.45-fold and 1.74-fold increase over a random phage set in *E. coli* and *Klebsiella*, respectively (*Supp Fig. 30*). This could reduce the experimental overhead for researchers or clinicians attempting to select phages targeting under-studied host phylogenies.

A detailed overview of performance across model configurations and datasets, potential biological interpretations, and types, sources, and implications of model errors can be found in the Supplementary Materials (*Supp. Materials - 1.1.4* / *Supp. Fig. 31-35*).

### Phage Selection and Phage Cocktail Design

Having established that models generate reliable probabilistic predictions of phage-host interactions, we next evaluated how these predictions could be translated into practical phage selection and cocktail design strategies. Applying predictive models to microbiome engineering or phage therapy involves selecting phages targeting a specific host strain based on model predictions. As models generate continuous probability scores (0-1) representing the likelihood of phage-host interaction, these probabilities must be translated into actionable phage selection strategies. In the context of phage therapies, combining multiple phages into “phage cocktails” simultaneously increases infection probability through multiple independent mechanisms while reducing the likelihood that target bacteria possess or can evolve resistance to all cocktail components^13,15,37^. We evaluated model-based phage selection strategies and phage cocktail design by conducting *in silico* experiments where models selected phage sets to target unseen bacterial strains from our cross-validation holdouts (*Supp. Fig. 36-38* / *Supp. Materials - 1.1.5*). The final cocktail design workflow balances predicted interaction probability and phage mechanistic diversity through unsupervised phage clustering into “activity groups” based on interaction profiles and the selection of the top predicted phage across clusters (*Fig. 3A-B*)^48^.

**Figure 3.**
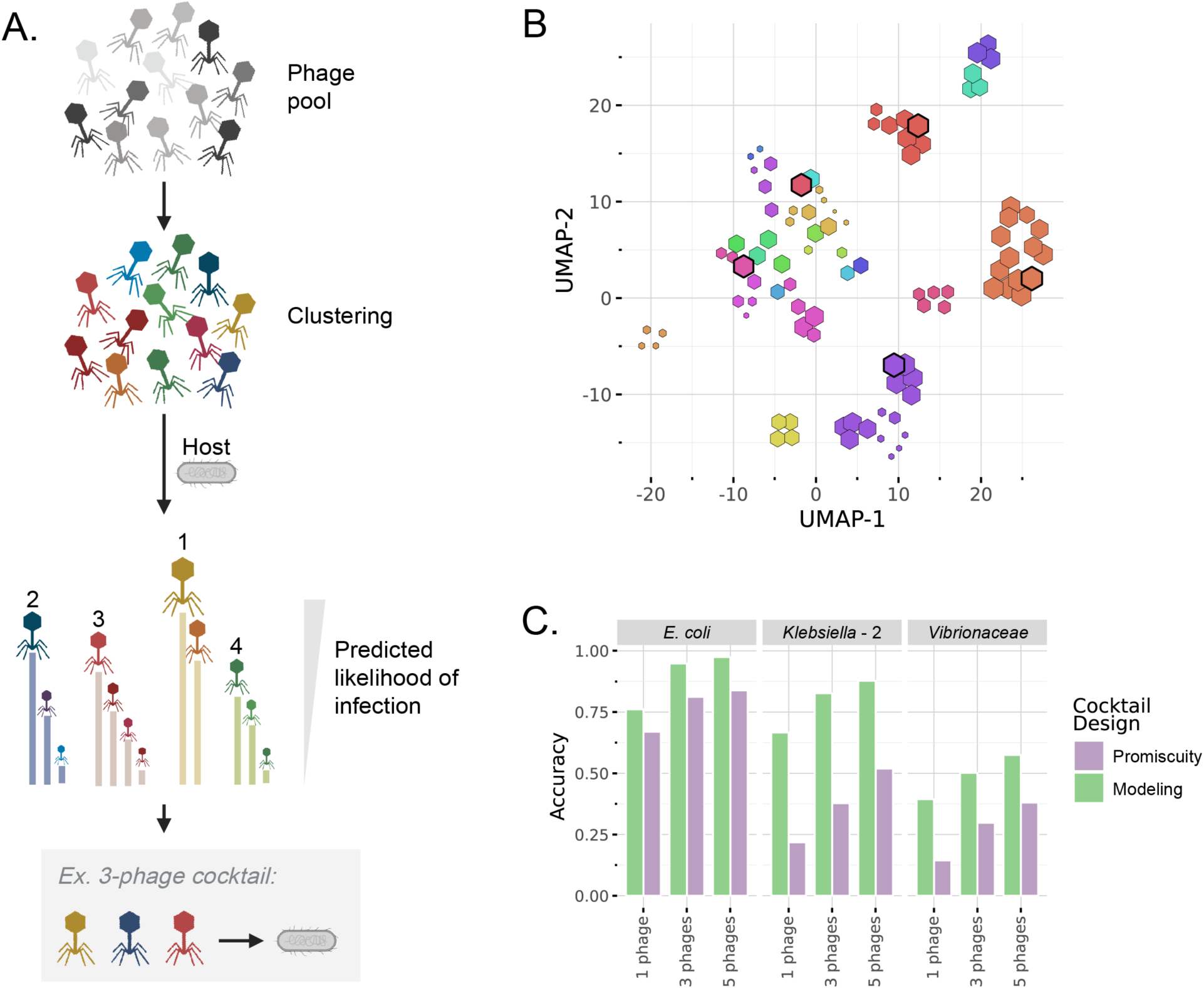
Phage cocktail design workflow and in-silico validation. A) The phage-cocktail design workflow involves HDBSCAN clustering of phages based on predictive feature content, ranking of phages within each cluster based on model predictions and selecting the top-performing phages from *n* clusters. B) UMAP projection shows the distribution of phages sized by their predicted likelihood of activity on *E. coli* ECOR13. Colors represent HDBSCAN phage clusters and selected phages are highlighted in black. C) Phage cocktails based on model predictions (green) significantly outperformed cocktails based on phage promiscuity (purple) across the three largest and highest-performing datasets.

We tested single phage selections alongside 3-phage and 5-phage cocktails, comparing model-guided selection against baseline cocktails composed of the most promiscuous phages across clusters^13^, using 20-fold cross-validation. Model-guided cocktail design outperformed promiscuity-based selection in the three largest datasets (*E. coli*, *Klebsiella*-2, and *Vibrionaceae* datasets) across 1-, 3-, and 5-phage cocktails (*Fig. 3C*). For single-phage selection, models identified active phages for 66.9% of *E. coli* strains, 66.7% of *Klebsiella* strains, and 39.4% of *Vibrionaceae* strains, equating to 1.1-fold, 3.1-fold, and 2.7-fold increases over promiscuity-based cocktails, respectively. For 5-phage cocktails, models selected active phages for 97.5%, 87.8%, and 57.5% of *E. coli*, *Klebsiella*, and *Vibrionaceae* strains, respectively, representing 1.2-fold, 1.7-fold, and 1.5-fold improvements over promiscuity-based cocktails. Overall, these results demonstrate that predictive models can be effectively integrated into a phage selection and cocktail design workflow, and that models can identify predictive features that meaningfully improve cocktail design beyond simple promiscuity-based heuristics. Additional information related to cocktail design workflow optimization and validation can be found in the Supplementary Materials (*Supp. Materials - 1.1.5*).

### Experimental Validation

Although cross-validation experiments showed strong performances across multiple datasets, for this modeling workflow to be broadly applicable by researchers, clinicians, and engineers we need to confirm that it is identifying generalizable genetic features that can predict phage-host interaction in multiple experimental contexts. To validate model generalizability and the biological relevance of predictive genetic features, we generated two complementary high-throughput experimental datasets. The first is a 1,300-interaction matrix testing 52 novel phages (*Supp. Table 3*) against 25 *E. coli* strains present in the modeling dataset. The second is a series of genome-wide RB-TnSeq screens testing the importance of 3,804 gene knockouts in *E. coli* ECOR27 to interaction with a panel of 21 phages (*Supp. Table 7*).

The interaction matrix allowed us to directly test predictive performance of the *E. coli* dataset models on experimental data generated in a distinct laboratory environment. As we did not have access to any of the phages used in the published dataset, we could not test the primary model configuration predicting phages active against unseen bacterial strains. We instead selected 25 *E. coli* strains belonging to the ECOR collection that were included in the published dataset (*E. coli* ECOR13 - ECOR38) and predicted interaction of these strains with a set of 52 previously unseen phages belonging to the BASEL phage collection^5,16,49–51^. Spotting assays were then performed with these phages on all 25 *E. coli* strains, resulting in 1,300 tested interactions of which 60 (4.6%) did not show a clear phenotype (*Fig. 4A*). Efficiency of plating (EOP) assays across a representative subset of strains and phages (18.5% of total interactions) confirmed the reliability of spotting assay phenotypes (*Supp. Fig. 39-41*). We then compared experimental plaquing phenotypes of the remaining 1,240 interactions to predicted activity, resulting in an AUROC of 0.84 (*Fig. 4B* / *Supp. Fig. 42-43*). This is slightly lower than the cross-validation performance (AUROC = 0.87) but shows the capacity of the models to generalize to unseen phages and phage-host interaction data generated in a distinct laboratory setting with different experimental protocols. To evaluate model generalizability outside of *E. coli*, we leveraged a published 3,318-interaction *Klebsiella* dataset to test the predictive performance of our *Klebsiella*-specific models (*Supp. Materials - 1.1.6*).

**Figure 4.**
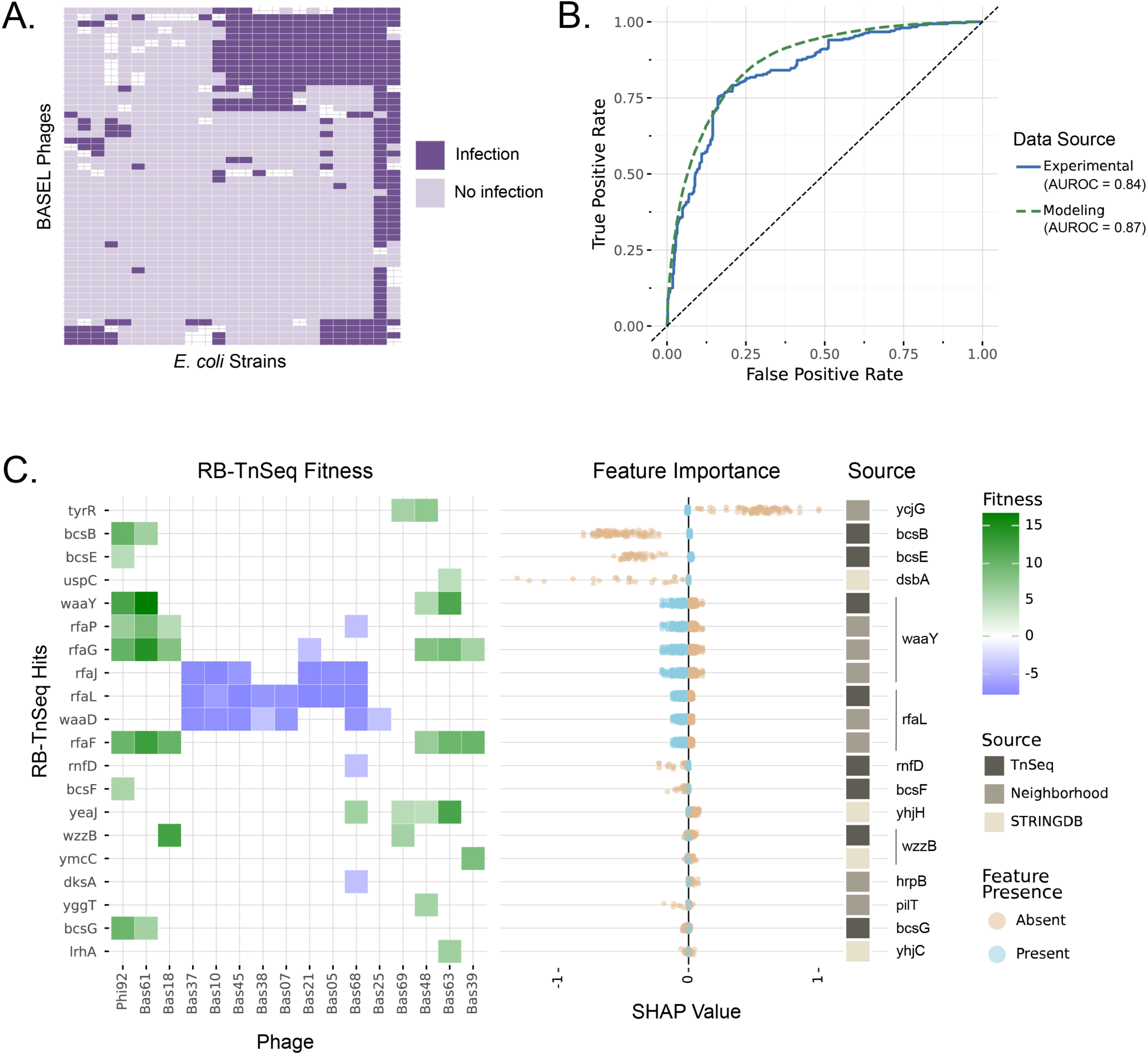
Experimental validation of model performance and predictive features. A) A 1,300-interaction one-to-one phage-host interaction matrix showing phage-host interaction outcomes from spotting assays. Dark purple shows interactions resulting in infection and light purple shows interactions not resulting in infection. Missing values indicate interactions where phenotypes were unclear. B) ROC curve showing performance of host predictions for previously unseen BASEL phages (blue) in comparison to the 20-fold cross-validation performance (dashed green). C) The heatmap shows fitness values associated with 20 RB-TnSeq hits linked to top predictive features in *E. coli* ECOR27. The X-axis shows tested phages and the Y-axis gene names of RB-TnSeq hits ordered by Shapley Additive exPlanations (SHAP) feature importances. SHAP values indicate whether presence of a given feature (blue) or absence (orange) is associated with increased likelihood of infection (SHAP value > 0) or decreased likelihood of infection (SHAP value < 0). The colored bar shows the relationship between RB-TnSeq hits and predictive features, including whether the predictive feature is directly linked to the gene in question (dark brown), linked through a neighborhood analysis (medium brown), or linked through STRING-DB (light brown). Gene names on the far right show the annotation associated with the predictive feature.

To validate the biological relevance of the predictive features, we performed an RB-TnSeq genetic screening experiment in *E. coli* ECOR27, which was included in both the modeling dataset and our interaction matrix described above. In these experiments, pooled genome-wide mutant libraries are subjected to phages, allowing us to identify host factors associated with phage infection through the increased fitness of their mutants^7^. Although these high-throughput genetic screening experiments are known to enrich for phage receptors, whose regulation has the most significant impact on host fitness^7,52^, this comparison revealed that our workflow successfully identified multiple screening hits, related regulators, or genes within the same operon or gene cluster as predictive features. The *E. coli* ECOR27 library was screened against 21 phages, identifying 51 genes as high-scoring hits for phage infection in this host (*Supp. Fig. 44*). We then compared these RB-TnSeq hits to predictive features, ranked based on SHapley Additive exPlanations (SHAP) values as a metric of model feature importance. Of the 51 genes identified through RB-TnSeq, 12 mapped directly to predictive features, 10 fell within a 3-gene window of a predictive feature (4 were directly adjacent), and 13 were functionally related to predictive features through STRING-DB^53^, totaling 35 predictive genes with links to RB-TnSeq hits (68.6%) (*Fig. 4C* / *Supp. Fig. 45*). With 155 features assigned to *E. coli* ECOR27, corresponding to 217 genes (a total of 1060 genes once expanded to include those identified through neighborhood and STRING-DB analyses), this represents significant enrichment in mechanistically important genes, with direct matches 5.0-fold above random expectation (p = 2.8 × 10^−6^) and a 3.0-fold enrichment for the neighborhood and STRING-DB extended set (p = 4.3 × 10^−12^) (*Supp. Fig. 46*). Although multiple genes in single-gene functional clusters often all showed fitness phenotypes in RB-TnSeq data, we observed that the predictive models would frequently only identify a single member of these pathways, including regulators for porins *tsx*, *fadL*, and *nfrA*, as well as single genes in LPS, O-antigen, and capsule biosynthesis clusters (*Fig. 4C*). This pattern demonstrates that the models may capture pathway- or operon-level regulatory mechanisms rather than just direct gene effects. We also compared annotations of predictive features to published RB-TnSeq, DubSeq, and CRISPRi experiments, which identify host genes affecting phage susceptibility through transposon mutagenesis, overexpression, and inactivation screens, respectively^7^. This revealed multiple additional screening hits or related genes that our workflow successfully identified as predictive features. These included multiple porins known to be primary phage receptors (*ZuA*, *nfrA*, *ompC*, *ompF*, *ompT*), contributors to outer-membrane structure (*bcsA*, *waaY*), and regulators of key metabolic and surface-associated pathways (*Supp. Materials 1.1.6*).

These complementary experimental validations demonstrate that our approach generalizes effectively to novel datasets while capturing biologically meaningful determinants of phage infection, providing confidence that computationally identified features represent genuine mediators of phage-host interactions and that resulting models could be valuable tools in phage cocktail design and targeted microbiome engineering.

### Biological Interpretation

A key objective of this modeling workflow was to enable biological interpretation of predictions by identification of specific genetic determinants of phage-host interactions. Protein family features allow direct mapping of predictive features to genes and gene variants that may mediate infection. Having demonstrated through RB-TnSeq experiments that computationally identified features are enriched in functionally important genes, we sought to understand the mechanistic basis of these predictions and the relationship between these features and known mediators of phage-host interaction across datasets.

We leveraged SHAP values to quantify feature importance across datasets and classify model features based on their predicted impact on infection^54,55^. Features were categorized as having a positive impact when their presence increased infection probability or a negative impact when their presence decreased infection probability. To systematically interpret these findings, we used established bioinformatics workflows to classify host proteins associated with predictive features based on genomic origin (viral, plasmid, or chromosomal)^24^, cellular localization (cytoplasm, membrane, or extracellular)^56^, relationship to known phage-host mechanisms, and classification as defense or anti-defense systems^57^ (*Supp. Fig. 47*). Across the four development datasets, we did not observe an enrichment of features with either “positive” or “negative” impacts across any feature functional classifications, apart from defense and anti-defense systems, where defense systems typically had negative impacts on infection while anti-defense systems had positive impacts.

Analysis of the top 25 most predictive features in the *E. coli* dataset revealed various functional categories associated with known phage-host interaction mechanisms (*Fig. 5A*). These included features linked to cell wall biosynthesis, mobile genetic elements, RM systems, and genes of viral origin, with varying directional impacts on interaction probability, alongside eight features with no clear mechanistic link, potentially representing novel mediators. STRING-DB and gene neighborhood analyses of the top predictive features suggested potentially novel mechanisms, including a role for putrescine catabolism^58^ and regulation of the known phage receptor *fepA*^59^, warranting further experimental investigation (*Supp. Materials 1.1.7*).

**Figure 5.**
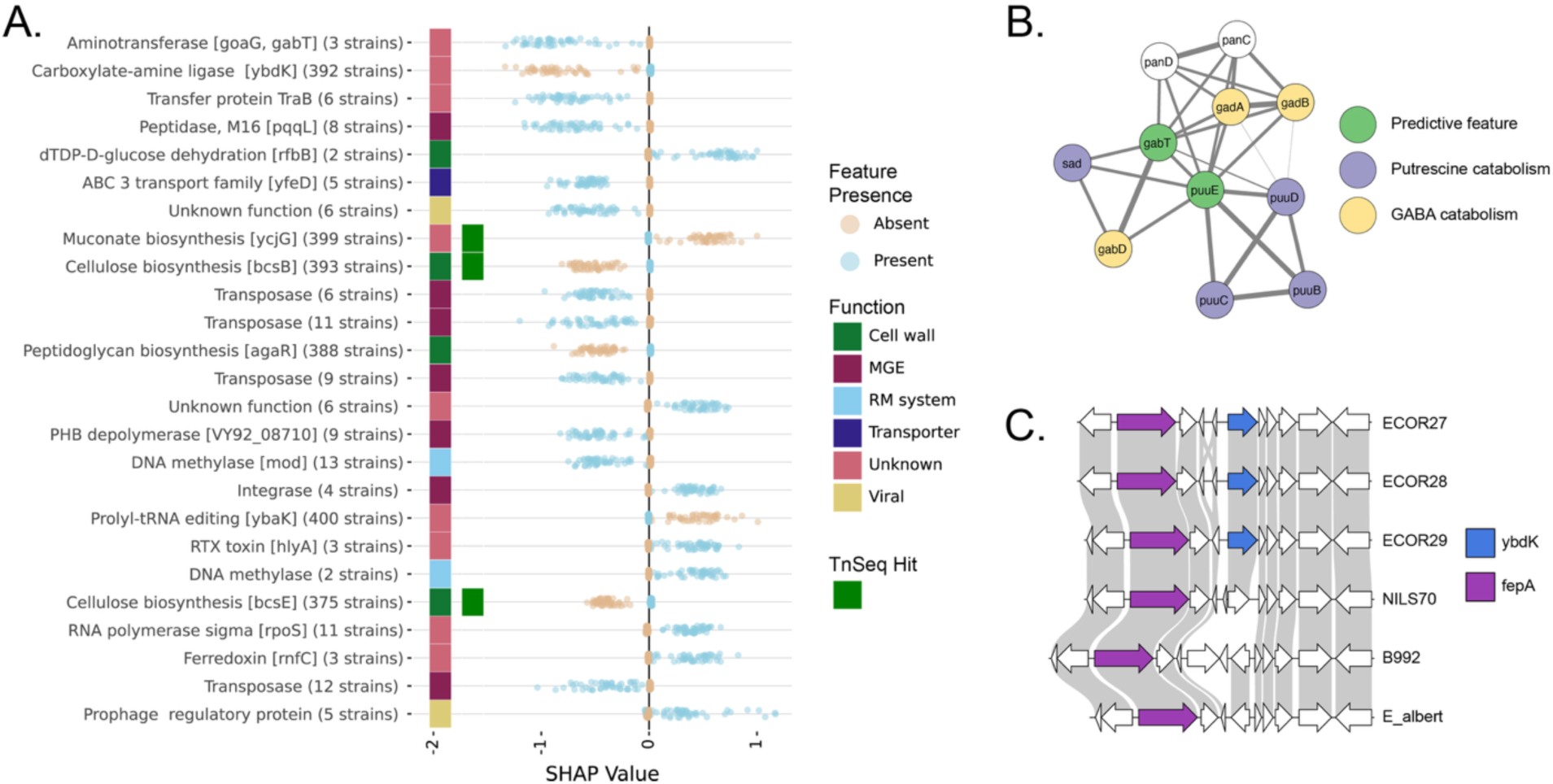
Biological interpretation from predictive features. A) Feature annotations ordered by Shapley Additive exPlanations (SHAP) feature importances. SHAP values indicate whether presence of a given feature (blue) or absence (orange) is associated with increased likelihood of infection (SHAP value > 0) or decreased likelihood of infection (SHAP value < 0). The colored bar shows functional categories associated with predictive features or whether the feature is associated with an RB-TnSeq hit in *E. coli* ECOR27. B) The STRING-DB network associated with genes *goaG* (*puuE*) and *gabT*. C) Synteny plot showing *fepA* adjacent to predictive feature *ybdK* in three strains with and three strains without *ybdK*.

## Discussion

Recent studies have advanced machine learning approaches for predicting bacteriophage host range, exploring various genomic representation strategies and modeling algorithms^13,16,17,45,60^. However, most have remained restricted to single genera, fixed receptor mechanisms, or taxonomic classification tasks, and lack experimental or genetic validation of predicted determinants.

In contrast, our framework predicts binary infection outcomes between arbitrary phage-host pairs across multiple genera without requiring prior knowledge of receptor biology or species-specific mechanisms. Through recursive feature elimination, cluster-aware train-test splitting, and per-phage class weighting, we optimized model configurations to produce a phylogeny-agnostic workflow that generalizes across five independent datasets. Direct performance comparison demonstrated our approach matched existing methods (AUROC 0.87 vs. 0.86 for *E. coli* and AUROC 0.812 vs. 0.818 for *Klebsiella*) while removing major constraints: the published methods required features based on known infection mediators, whereas our framework achieved comparable performance across diverse bacterial genera without these limitations. Crucially, we demonstrate external generalization to 52 unseen phages from the independent BASEL collection (AUROC 0.84) and mechanistic concordance with genome-wide RB-TnSeq assays, in which 68.6% of genetically validated host-determinant genes align with computationally identified predictive features.

The workflow’s key innovation lies in converting complex whole-genome data into numerical feature representations that capture sufficient biological information for accurate prediction without mechanistic assumptions, while minimizing overfitting through dynamic feature filtering and selection. Extensive workflow optimization revealed that algorithmic choices in feature selection and machine learning implementation had greater impact on performance than genomic representation strategies, indicating that extracting relevant signals from high-dimensionality data represents the primary computational challenge. Integration with cocktail design strategies translated these probabilistic predictions into actionable therapeutic selection, achieving 57.5-97.5% bacterial strain coverage across datasets using 5-phage cocktails and up to 3.1-fold improvements in single phage selection over promiscuity-based methods. Despite this strong performance, training and validation were limited to laboratory conditions and modeling was based entirely on bacterial and phage genomes, assuming static conditions. Future models and datasets should incorporate environmental conditions influencing bacterial gene regulation and phage infection dynamics.

Biological interpretation of predictive features revealed both established and potentially novel genetic determinants of phage susceptibility while providing mechanistic insights into infection processes. RB-TnSeq experiments directly validated the functional relevance of computationally identified features, including established receptors (*tsx*, *nfrA*) and cell wall biosynthesis pathways. Importantly, the modeling approach frequently identified single regulatory genes within broader pathways rather than all pathway components, suggesting the framework could capture pathway-level regulatory mechanisms rather than just direct gene effects. However, feature validation through RB-TnSeq is limited to non-essential genes showing strong fitness effects under phage selection, preventing its use in validation of modeling features associated with essential genes. Future studies should consider alternative validation methods, such as CRISPRi, which allow perturbation of essential genes and could provide a more comprehensive view of genetic factors influencing phage-host interactions.

Performance correlations with dataset characteristics highlight strategies for collaborative model development while revealing current limitations. Larger datasets consistently improved performance, with the requirement for substantial positive interaction data currently constraining application to a limited set of host systems. However, successful combination of *Klebsiella* datasets from different research groups demonstrated viable strategies for distributed dataset generation, suggesting that standardized experimental protocols could enable community-driven model improvement. The absence of correlation between bacterial phylogenetic relatedness and prediction accuracy within single phylogenetic groups likely reflects, at least in part, the removal of phylogenetically linked features during filtering. However, the observation that adding closely related strains to training data does not consistently improve prediction accuracy suggests a more complex relationship between genotype and phage susceptibility than phylogenetic proximity alone would predict, consistent with a continuous arms race and an important role of horizontally transferred elements as determinants of phage susceptibility. The poor performance of cross-genus prediction and the lack of performance improvement when combining phylogenetically-diverse datasets suggest either genus-specific determinants of phage-host interaction or the limitations of protein-family-based features for capturing mediators of phage infection conserved across genera. Differences in assay conditions between datasets, including solid versus liquid medium and varying susceptibility thresholds, may also contribute to reduced performance when combining cross-genus or cross-laboratory datasets, as phenotypic definitions of a positive interaction are not fully standardized. In the future, models that incorporate environmental conditions and use alternative genomic representations that capture structural information, such as protein language model embeddings, may improve cross-genus and cross-dataset transfer learning.

An additional potential application of this framework is the prediction of interactions between phages and bacterial hosts co-identified within the same microbial community. For instance pairing phage and bacterial metagenome-assembled genomes (MAGs) within a single metagenome. Unlike existing taxonomic host-assignment tools, our approach could predict one-to-one interaction probabilities between phage and host genome pairs within a single community. However, as cross-genus prediction performance remains poor, this application would require interaction datasets for the target host phylogeny, or validation with microbial community datasets containing known phage-host interactions. Feature assignment to MAGs may also be complicated by genome incompleteness. Exploring this application represents an interesting direction for future work.

This framework establishes computational infrastructure for rational phage therapy design and precision microbiome engineering applications. The demonstrated ability to achieve broad strain coverage across phylogenetic clades addresses key requirements for therapeutic efficacy and resistance management. Future developments should prioritize active learning approaches that leverage model confidence metrics to strategically guide experimental design. Our analysis revealed a strong correlation between model confidence and prediction reliability, suggesting that uncertain predictions (those near the 0.5 classification threshold) could identify high-value experiments for model improvement. Such adaptive experimental design could substantially reduce the scale of interaction screening required while maximizing information gain. As antimicrobial resistance continues to escalate globally, this framework provides a scalable platform for evidence-based therapeutic design.

## Methods

### Dataset Collection and Preprocessing

Six phage-host interaction datasets with binary phenotype data and corresponding genomes were obtained from published sources^16,17,38,40–42^. Four datasets were used for model development: *E. coli* subset (16,638 interactions, 177 strains × 94 phages)^16^, mixed *Klebsiella* spp. (3,658 interactions, 62 strains × 59 phages)^40^, *Klebsiella pneumoniae* (6,348 interactions, 138 strains × 46 phages)^41^, and *Pseudomonas* (437 interactions, 23 strains × 19 phages) ^42^. The mixed *Vibrionaceae* dataset (63,488 interactions, 256 strains x 248 phages)^38^, a third *Klebsiella* dataset (3,318 interactions, 68 strains × 25 phages), and the full *E. coli* dataset (37,788 interactions, 402 strains x 94 phages)^16^ were reserved for validation. Development datasets totaled 27,081 interactions including 4,304 positive interactions (2.4%-36.2% positive rate across datasets). The *E. coli* interaction matrix was generated by assaying on solid medium, with susceptibility qualitatively assessed based on the clearance achieved by three phage dilutions and positive interactions defined as any non-zero interaction. All other datasets were generated by assaying in liquid medium and relied on the area under the optical density curve to quantify susceptibility, with positive interactions defined based on thresholds set by publishing authors for other datasets.

### Phylogenetic Analysis

Bacterial strain phylogenies were inferred from core genome alignments using PPanGGOLiN (v2.0.3)^61^. Pangenome analysis identified persistent (core) genes across all bacterial strains in each dataset, and multiple sequence alignments of concatenated amino acid sequences from these genes were generated. Maximum-likelihood phylogenetic trees were constructed using FastTree (v2.1.11) with default parameters^62^. Pairwise patristic distances (sum of branch lengths between terminal nodes) were calculated for all strain pairs using BioPython’s Phylo module, generating complete distance matrices for each dataset.

Phage evolutionary distances were calculated using PhamClust protein family similarity scores (PEQ values), which quantify shared gene content between phage pairs^63^. PEQ similarity scores were converted to distance metrics as d(i,j) = 1 - PEQ(i,j), where d(i,j) represents the distance between phages i and j. This approach captures both sequence similarity and gene content divergence. Phage gene-sharing networks were additionally constructed using vConTACT2 (v0.11.3) to visualize evolutionary relationships and identify phage clusters based on shared protein content across datasets^18^.

Phylogenetic isolation was quantified for each bacterial strain or phage as the mean patristic distance to its 5 nearest phylogenetic neighbors, excluding self-distance. This metric captures local phylogenetic context rather than global tree position and provides a measure of how evolutionarily distinct each entity is from its closest relatives. Correlations between phylogenetic isolation and prediction performance (MCC) were assessed using Pearson correlation coefficients with two-tailed significance tests (α = 0.05).

### Protein Family Construction

Protein sequences from all genomes were clustered using MMSeqs2 (v. 15.6f452) with optimized parameters^64^. MMSeqs2 databases were created using the `createdb` command for all input FASTA files. Protein clustering was performed using the `cluster` command with sequence identity threshold 0.4, coverage requirement 0.8, and sensitivity 7.5. The complete clustering workflow used the following MMSeqs2 commands:

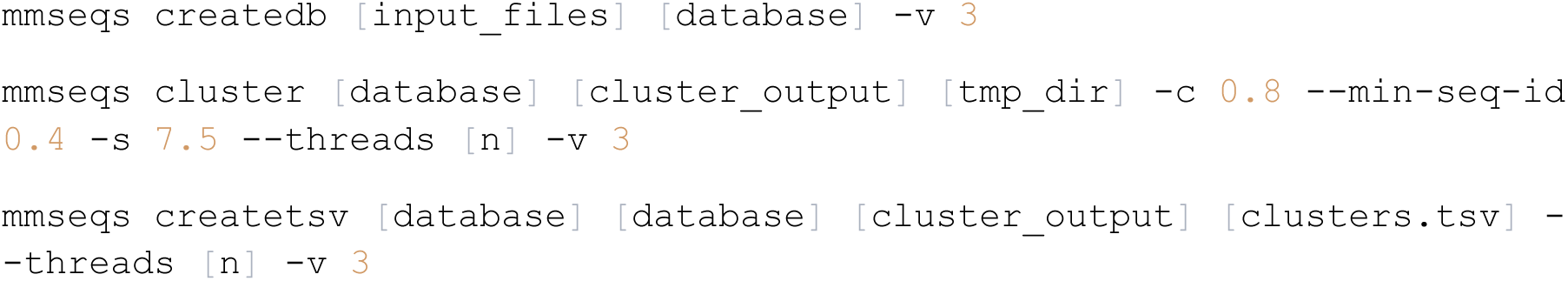

Clusters present in only a single genome were removed to focus on shared protein families. Binary presence-absence matrices were constructed with genomes as rows and protein clusters as columns. Clusters with identical presence-absence patterns were consolidated using hash-based grouping to reduce redundancy.

Parameter optimization tested pairwise combinations of MMSeqs2 identity thresholds (0.2, 0.4, 0.6, 0.8, 0.9) with fixed coverage of 0.8 across 25 parameter combinations using mixed-effects modeling. Parameter optimization was performed within ensemble-learning runs, with performance metrics calculated for each threshold combination across multiple modeling iterations. Model predictions were aggregated using median confidence scores across iterations before calculating performance metrics for each parameter combination. The model formula `performance_metric ∼ C(strain_identity) * C(phage_identity)` with dataset as random effect was used to validate parameter selection. Results showed bacterial strain and phage identity thresholds explained 0.0% to 3.35% of Matthews Correlation Coefficient (MCC) variance across datasets, confirming minimal impact of parameter choice. When standard fitting failed due to convergence issues, optimization used the Limited-memory Broyden-Fletcher-Goldfarb-Shanno (LBFGS) method, a quasi-Newton optimization algorithm that provides improved convergence for complex mixed-effects models^65^.

An alternative clustering approach using DIAMOND (version 2.0+) and Markov Clustering (MCL) was evaluated for comparison^66,67^. This workflow employed DIAMOND blastp for homology search with sensitive mode and e-value threshold of 0.001, followed by MCL clustering with pairwise testing of inflation parameter values of 1.4, 2.0, 4.0, and 6.0 for phage and bacterial genomes. DIAMOND results were converted to ABC format for MCL input, and resulting clusters were processed to generate equivalent presence-absence matrices. While this approach resulted in comparable model performance, MMSeqs2 was selected for final analyses due to superior computational efficiency.

### K-mer Feature Construction

Amino acid *k*-mers of *length k* were extracted from all proteins using sliding window approaches. *K*-mers present in only a single gene were filtered. Binary presence-absence matrices were generated for unique *k*-mers either across complete proteomes or within protein families. *K*-mer features were assigned to genomes based on exact subsequence matching or protein family membership and exact subsequence matching, respectively. Hash-based consolidation was applied to *k*-mers with identical occurrence patterns across genomes. Alternative k-values (3-15) were tested showing minimal performance impact, but *k*=6 resulted in the most consistently high performing models in training data. For complete proteome feature tables, the theoretical maximum number of amino acid *k*-mers is 20*^k^*. While the observed biological feature space is significantly sparser, the number of unique presence-absence patterns after hash-based consolidation still increases significantly with *k* (*Supp. Fig. 22* / *Supp. Table 3*). A value of *k*=4 was therefore used for complete proteome feature tables as increasing above this value led to untenably large feature tables that challenged computational tractability during iterative ensemble modeling.

For novel genome feature assignment, *k*-mer matching required presence of exact *k*-mer subsequences within assigned protein sequences. A permissive threshold of 0.001 was used for *k*-mer assignment, allowing features to be assigned when ≥0.1% of *k*-mers were present in the assigned proteins.

### Machine Learning Algorithm Selection and Optimization

Eight machine learning algorithms were systematically compared across all development datasets: CatBoost gradient-boosted decision trees (CatBoost)^68^, random forest, *k*-nearest neighbor (KNN), logistic regression, multilayer perceptron (MLP), naïve Bayes, support vector machines (SVM), and gradient boosting^69^. Algorithm comparison employed comprehensive grid search hyperparameter optimization with algorithm-specific parameter ranges tested across multiple iterations. Grid search parameters included:

- **CatBoost**: iterations (500, 1000), learning_rate (0.05, 0.1), depth (4, 6)
- **Random Forest**: n_estimators (100, 200), max_depth (None, 10, 20)
- **Logistic Regression**: C (0.1, 1, 10), solver (’liblinear’, ‘lbfgs’)
- **SVM**: C (0.1, 1, 10), kernel (’linear’, ‘rbf’)
- **Gradient Boosting**: n_estimators (100, 200), learning_rate (0.01, 0.1), max_depth (3, 5)
- **KNN**: n_neighbors (3, 5, 7), weights (’uniform’, ‘distance’)
- **MLP**: hidden_layer_sizes ((50,), (100,), (50, 50)), activation (’tanh’, ‘relu’)

Model evaluation employed bacterial strain-based train-test splitting (80% strains for training). Performance was evaluated using multiple metrics (MCC, AUROC, accuracy, precision, recall, F1-score) with best models selected based on Matthews Correlation Coefficient to account for dataset imbalance. Statistical significance was assessed using Mann-Whitney U tests for pairwise algorithm comparisons. Model predictions were aggregated using median confidence scores across 100 modeling iterations, with 0.5 threshold for binary classification.

CatBoost gradient-boosted decision trees demonstrated the most consistent high-performance across datasets, achieving equal or superior MCC scores compared to alternative algorithms while also minimizing computational resource requirements.

Phage-specific class weights were implemented to address dataset imbalance with two available methods: ‘log10’ and ‘inverse_frequency’. For each phage, positive and negative interaction counts were calculated separately, and weights were computed as follows:

- `Log10 method: weight_1 = max(1.0, log10(neg_count / (pos_count + smoothing)) + 1)`
- Inverse frequency method: `weight_1 = total_count / (pos_count + smoothing)`

where smoothing = 1.0 to prevent extreme weight ratios when positive counts are low. Negative class weights were set to 1.0 for all phages, while positive class weights were adjusted based on the phage-specific positive/negative ratio. This approach accounts for the varying interaction rates across different phages in the dataset.

Ensemble predictions were generated using median values across 50 modeling iterations to improve prediction stability and reduce overfitting.

### Feature Selection Methodology

Six feature selection methods were systematically compared: Recursive Feature Elimination (RFE), SHAP-based selection, SHAP-RFE hybrid, LASSO, chi-squared, and SelectKBest. RFE demonstrated optimal performance in 3 of 5 datasets with significantly lower computational requirements and was therefore selected for final analyses. Performance was evaluated across multiple datasets using Mann-Whitney U tests for pairwise method comparisons.

In the final modeling workflow RFE was implemented using CatBoost as the base estimator with the following configuration:

- n_features_to_select: automatically determined or specified
- step_size: `max(1, int((total_features - n_features_to_select) / 10))`

RFE was performed for 25 rounds with different train-test splits, with 10 feature elimination steps per round. Features were ranked by occurrence frequency across feature selection iterations, with final feature tables generated using occurrence cut-offs. Final models used features present across the maximum number of iterations, including no more than total_features/20 features in the final selection. This feature occurrence tracking across iterations enabled identification of stable, predictive features while reducing overfitting to specific data splits. Final models used features selected based on optimal occurrence thresholds determined by model performance.

### Dataset Balancing and Overfitting Prevention

Dataset imbalance (2.1%-36.2% positive interactions) was addressed through systematic comparison of 14 parameterization strategies across three methodological dimensions: clustering methods for train-test splitting (HDBSCAN, hierarchical, or random splitting), phage-specific class weighting approaches (log10 or inverse-frequency), and feature filtering strategies (cluster-based, train-test presence, or no filtering). Strategy performance was evaluated using twenty-fold cross-validation with 10% of bacterial strains systematically withheld from the entire feature selection and modeling workflow.

The optimized approach combined: (1) hierarchical clustering of bacterial strains into 20 clusters using Ward linkage and Euclidean distance (2) inverse-frequency class weights computed per phage, and (3) feature filtering requiring presence across ≥2 clusters to exclude features unique to single bacterial groupings. This strategy provided optimal balance between model generalizability and performance while addressing dataset-specific imbalance and strain heterogeneity.

### Cross-Validation and Model Evaluation

Model generalization was evaluated using three distinct model configurations designed to assess different aspects of biological transferability. Twenty-fold cross-validation was performed for each strategy, with 10% of samples systematically withheld from the entire feature selection and modeling workflow to ensure unbiased evaluation. Features for validation genomes were assigned using similarity search (MMSeqs2, sequence identity ≥0.4, coverage ≥0.8) against training protein clusters to simulate realistic inference conditions.

Host strain-based cross-validation evaluated generalization to novel bacterial hosts by training models on 90% of strains (using all available phages) and validating predictions on the remaining 10% of strains. This approach tests the model’s ability to predict phage susceptibility for previously unseen bacterial genotypes. The phage-based configuration assessed generalization to novel phages by training on all bacterial strains with 90% of phages and validating on interactions between all strains and the held-out 10% of phages. This strategy evaluates model performance when predicting host ranges for unseen phages. Bacterial strain and phage-based cross-validation examined the most challenging scenario of complete novelty by training exclusively on interactions between 90% of host strains and 90% of phages (81% of total interactions), then validating on interactions between the held-out 10% of strains and 10% of phages. This approach tests generalization to entirely novel bacteria-phage combinations representing real-world discovery scenarios.

Statistical comparison of modeling parameters using cross-validation employed rank-order analysis with median performance metrics across the 20 iterations per parameters set. Pairwise differences were assessed using Mann-Whitney U tests, and overall parameterization differences were evaluated using the Friedman test for non-parametric comparison of ranks across multiple datasets.

Model performance was quantified using multiple complementary metrics implemented in scikit-learn (v1.5.2) to account for dataset imbalance and provide comprehensive evaluation^69^. Predicted probabilities were converted to binary predictions using a classification threshold of 0.5 for calculation of Matthews Correlation Coefficient (MCC), accuracy, precision, recall, and F1-score. MCC was selected as the primary performance metric because it provides a balanced measure accounting for all four confusion matrix categories (true/false positives and negatives), ranging from -1 (complete disagreement) to +1 (perfect prediction), with particular suitability for imbalanced datasets.

Threshold-independent metrics included area under the receiver operating characteristic curve (AUROC), which measures the model’s ability to rank positive instances higher than negative instances with values from 0.5 (random) to 1.0 (perfect), and area under the precision-recall curve (AUPR), which quantifies precision-recall trade-offs and is particularly informative for imbalanced datasets. AUPR values were normalized relative to random baseline performance as AUPR_normalized = (AUPR - baseline) / (1.0 - baseline), where baseline equals the proportion of positive interactions. This normalization enables direct comparison across datasets with different imbalance ratios, with values ranging from 0 (random) to 1 (perfect).

Prediction calibration was assessed using Brier score (described in Model Calibration Analysis section) and calibration curves comparing predicted probabilities to observed infection frequencies.

### Feature Assignment Protocol for Novel Genomes

Novel genome proteins were queried against reference protein clusters using MMSeqs2 similarity search with optimized parameters: sensitivity 7.5, coverage ≥0.8, and sequence identity threshold ≥0.4. The search was performed using:

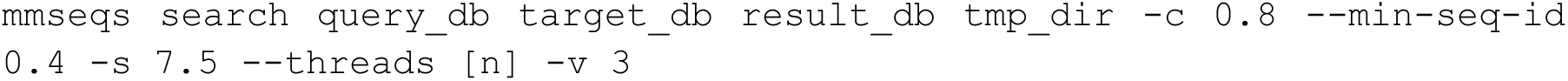

Each protein was assigned to the best-matching cluster above the identity threshold, with duplicate protein IDs resolved by prefixing with genome identifiers when detected. Cluster assignments were then mapped to specific features using the feature mapping established during model training, which defined the relationship between protein clusters and selected features from the reference dataset. Features were assigned to a host strain or phage if a single protein family from a given feature was present. Binary feature vectors were constructed indicating protein family presence/absence for each genome based on these cluster-to-feature mappings.

*K*-mer features were assigned based on amino acid sequence content, requiring exact *k*-mer subsequence matches within assigned protein sequences. A permissive threshold of 0.001 was used for *k*-mer assignment, allowing features to be assigned when ≥0.1% of *k*-mers within a protein family were present in the assigned proteins. This protocol ensured feature assignment reflected realistic conditions where novel genomes lack complete sequence identity to reference databases while maintaining biological relevance through sequence similarity requirements.

### Model Error Analysis

To identify systematic failures in phage-host interaction prediction and prioritize targets for biological investigation, we developed a comprehensive error analysis framework consisting of five complementary approaches.

## High-Confidence Misclassification Analysis

High-confidence misclassifications were identified as predictions where *C*_*abs*_ > 0.8 and the prediction disagreed with ground truth. These represent cases where the model is most confident but incorrect, indicating systematic feature misinterpretation rather than uncertain boundary cases. The threshold of 0.8 corresponds to model confidence scores <0.1 or >0.9, representing the top quintile of model certainty.

## Systematic Bias Detection

Entities exhibiting consistent over- or under-prediction were identified using prediction bias thresholds of ±0.2, corresponding to systematic errors of >20 percentage points between predicted and observed interaction rates. This threshold captures biologically meaningful differences while excluding noise from natural prediction variance. Entities were classified as over-predictors (*Bias*_*i*_ > 0.2) or under-predictors *Bias*_*i*_ < 0.2.

## Interaction Pattern Analysis Using Jaccard Similarity

To identify cases where the model fails to distinguish biologically distinct interaction patterns, we implemented pairwise comparisons using Jaccard similarities. For each bacterial strain or phage pair (i, j), we calculated model prediction similarity 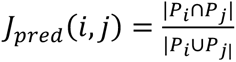 and true interaction similarity 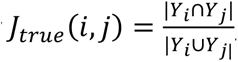, where *P_i_* and *Y_i_* represent binary prediction and interaction vectors for entity i across all tested partners.

This approach directly compares interaction overlap patterns rather than summary statistics, capturing the complete interaction landscape and enabling direct comparison of predicted outcome to experimentally determined interaction patterns. Analysis was conducted independently for both strain-level and phage-level perspectives.

Extreme discordance cases were identified as high-priority targets for biological investigation: Type I cases (true Jaccard similarity < 0.3 with model similarity > 0.9) represent cases where the model incorrectly predicts nearly identical interaction patterns for biologically distinct entities, while Type II cases (true Jaccard similarity > 0.7 with model similarity < 0.3) represent cases where the model fails to recognize highly similar interaction patterns.

## Individual Strain Performance Analysis

Individual strain performance was quantified using standard binary classification metrics (MCC, AUROC, accuracy, precision, recall) calculated from each strain’s complete set of phage interaction predictions. AUROC calculations required at least two distinct classes; strains with exclusively positive or negative interactions were assigned AUROC = NaN. The relationship between strain infectivity (total number of infecting phages) and prediction performance was assessed using Pearson correlation analysis to determine whether models perform better on promiscuous versus restrictive strains.

## Confidence Distribution Analysis

Model confidence scores were analyzed across different prediction outcomes to identify systematic uncertainty patterns. Predictions were categorized by confusion matrix outcomes (TP, FP, TN, FN) and confidence distributions were visualized using kernel density estimation. Confidence scores were binned into 0.1-width intervals to assess model calibration and identify confidence ranges associated with specific error types.

## Model Calibration Analysis

Model calibration was assessed to evaluate the reliability of predicted probabilities and quantify prediction accuracy. Calibration curves were generated by binning predictions into 10 decile intervals (0.0-0.1, 0.1-0.2, …, 0.9-1.0) and calculating the mean predicted probability and observed infection frequency within each bin. Perfect calibration, where predicted probabilities match observed frequencies, is represented by the diagonal line y = x. Deviations from this line indicate over- or under-confidence in predictions.

The Brier score was calculated to provide a single metric of prediction accuracy that combines both calibration and resolution:

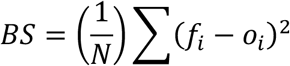

where *f_i_* is the predicted probability for interaction *i*, *o_i_* is the observed binary outcome (0 or 1), and *N* is the total number of predictions. Brier scores range from 0 (perfect accuracy) to 1 (complete inaccuracy), with scores <0.25 generally indicating useful predictions. Brier scores were calculated using scikit-learn’s brier_score_loss function across all predictions for each dataset and model configuration.

### Phage Cocktail Design Strategies

Phage cocktail design was evaluated using a factorial framework that varied phage ranking method, clustering strategy, and phage representation used for clustering. Phages were ranked using either (i) model-predicted interaction confidence or (ii) phage promiscuity estimated from the training data. Three cocktail design strategies were compared: direct rank-based selection without clustering, hierarchical clustering-constrained selection, and HDBSCAN-constrained selection. For clustering-based strategies, phages were represented either by host interaction profiles or by pangenome-derived feature vectors. In total, 10 cocktail design strategies were evaluated, with no-clustering strategies being independent of phage representation.

For prediction-based ranking, candidate phages were ordered by model-derived confidence scores for each strain in each held-out evaluation subset. For promiscuity-based ranking, each phage was assigned a global score equal to the fraction of training strains infected by that phage, calculated using only the non-held-out strains from the corresponding cross-validation iteration. This ensured that promiscuity estimates were derived exclusively from training data and prevented information leakage from the evaluation strains.

Cocktail selection was performed separately for each target strain. In the no-clustering condition, the top *n* phages were selected directly by ranking score, yielding a purely rank-based cocktail without explicit diversity constraints. In the hierarchical clustering condition, phages were clustered using Ward linkage with Euclidean distance, with the number of clusters set to the desired cocktail size *n*; the highest-ranked phage from each cluster was then selected. In the HDBSCAN condition, phages were clustered using density-based clustering with a minimum cluster size of 2. Phages labeled as noise were reassigned unique cluster identifiers so that they could still contribute as distinct candidates. The highest-ranked phage from each cluster was selected first; if fewer than *n* phages were obtained, additional phages were iteratively added in rank order from the remaining candidates until the desired cocktail size was reached. These clustering-based procedures were intended to reduce redundancy among selected phages while preserving mechanistic diversity.

When clustering was based on host interaction profiles, phages were represented by binary strain-by-phage interaction patterns derived from the training interaction matrix^13^. When clustering was based on pangenome features, phages were represented by their genomic feature vectors. This allowed comparison of diversity-aware cocktail design using either phenotypic similarity in host range or genomic similarity in gene content.

Performance was assessed as the fraction of target strains for which the selected cocktail achieved at least one experimentally confirmed interaction. Cocktail sizes of 1, 3, and 5 phages were evaluated. All analyses were conducted across 20 cross-validation iterations, and cocktail performance was summarized across held-out strains for each combination of ranking method, clustering procedure, representation type, and cocktail size.

### Bacterial Strains, Culture Media and Growth Conditions

*Escherichia coli* strains from the *E. coli* Reference (ECOR) Collection^50,51^ were acquired from the Thomas S. Whittam Shiga-toxin producing *E. coli* (STEC) Center (Michigan State University, USA). Host strains used for phage amplification were acquired with the corresponding phages (see below). Bacterial strains were kept in 15% glycerol for long-term storage, and routinely cultivated in LB broth and on LB agar plates at 37°C. Bacterial lawns for phage assays were prepared following a standardized double-agar overlay method: overnight bacterial cultures were grown from isolated colonies, and 100 µL were added to 10 mL of molten Tryptone Growth Top Agar (T-top agar) supplemented with CaCl_2_ at 10 mM. Suspensions were then poured onto rectangular Nunc™ Omnitray™ single-well plates (Thermo Scientific) filled with 30 mL of LB agar. Plates were allowed to solidify and dry for 15 min, and phages were spotted within the next hour.

### Phage Acquisition, Amplification and Titration

Phage aliquots from the BActeriophage SElection for your Laboratory (BASEL) collection^5,49^ were kindly provided by Prof. A. Harms (Biozentrum, University of Basel, Switzerland), and phage phi92^70^ was provided by Prof. Petr Leiman (Department of Biochemistry & Molecular Biology, University of Texas Medical Branch, USA). The remaining phages were from our lab stocks^7^. All phages were amplified using *E. coli* MG1655ΔRM as a host, except phage phi92 which was amplified on *E. coli* Bos12. For phage amplification, overnight cultures of the respective host strains were adjusted to an OD_600_ of 0.1 in 35 mL of LB broth supplemented with CaCl_2_ at 10 mM in 125-mL culture flasks. Isolated phage plaques obtained after two rounds of streak-purification were inoculated, and incubation was carried out at 37°C with shaking at 200 rpm for up to 4h or until complete clearance was observed. The resulting lysates were centrifuged for 10 minutes at 20,913 x g to pellet cell debris, and the supernatants were filtered on 0.22-µm Millex® polyethersulfone syringe filters (Millipore-Sigma). Phage titers were determined by serially-diluting lysates in SM buffer (Teknova) + 10 mM CaCl_2_ and spotting 2 µL droplets in triplicate on lawns of the appropriate bacterial hosts. High-titer phage lysates were kept at 4°C and considered stable over the duration of the experiments.

### Experimental Validation of Model Performance

52 BASEL collection phages were used for experimental validation on 25 *E. coli* strains from the ECOR collection present in the training dataset (ECOR13 - ECOR38)^50,51^. Phage suspensions were diluted with SM buffer + 10 mM CaCl_2_ in a Nunc™ 96-Well Polypropylene DeepWell™ plate (Thermo Scientific) to achieve target titers of 2.5 × 10^9^ PFU/mL based on phage titers determined on relevant amplification hosts. 100 µL of *E. coli* overnight cultures was added to the top agar of double overlay agar plates (approximately 10^8^ CFU). Using a Rainin MicroPro300 semi-automated pipettor (Mettler-Toledo Rainin), 2 µL droplets of all 52 phages were spotted in triplicate on bacterial lawns of the tested strains. This equates to an approximate ratio of phage virions to bacterial cells of 10. Phage clearance was manually recorded after 16-20h of overnight incubation at 37°C, and was scored as a binary phenotype (clearance/no-clearance). This clearance phenotype can result from productive infection (full viral replication), or non-productive lysis (mechanical lysis, endolysin activity, or abortive infection), representing a limitation of plaquing-based assays. Unclear phenotypes, where phage clearance was barely perceptible and/or not consistent over triplicates were excluded from the dataset. The curated phage-host interaction phenotypes were then compared to model predictions to evaluate model performance and generalizability.

A complementary experiment was conducted with subsets of bacterial strains and phages to verify whether the binary clearance phenotypes corresponded to productive phage infection or non-productive lysis^71,72^. A subset of 20 phages representative of the genomic diversity existing within the 52 tested phages was selected to determine their efficiencies of plating (EOP, ratio of the phage titer calculated on a test strain versus its titer on the amplification host strain) on a similarly representative subset of 12 *E. coli* ECOR strains. The selected phages were again normalized to 2.5 × 10^9^ PFU/mL and then serially diluted up to 10^−7^ in a 96-well PCR plate. Two µL droplets were spotted in triplicates on the amplification host strain MG1655ΔRM and the tested ECOR strains following the same protocol described above. Phage plaques were enumerated after an overnight incubation at 37°C, and titers were calculated by averaging triplicates. EOP values were then calculated by dividing the titer obtained on tested strains by the titer obtained on MG1655ΔRM.

### RB-TnSeq Validation of Predictive Features

The RB-TnSeq library in *E. coli* ECOR27 was built following published protocols by conjugating the pHLL250 mariner transposon vector library (AMD290) from donor strain *E. coli WM3064* ^46,73,74^. An aliquot of the barcoded donor library was thawed and allowed to recover at 37°C for 4h in 60 mL of LB supplemented with diaminopimelic acid (DAP) at 300 µM and carbenicillin at 50 µg/mL. The pellet was subsequently washed by centrifuging 5 min at 7,155 x g and resuspending in an equal volume of LB + DAP. Recipient strain ECOR27 was grown overnight at 37°C in 10 mL of LB. Mating was achieved by mixing the donor and recipient cultures at a 1:1 ratio, centrifuging 5 min at 7,155 x g, resuspending and spreading the pellet on a LB + DAP plate, and incubating for 5h at room temperature. The mixture was scraped and resuspended in LB before being plated on LB plates supplemented with kanamycin at 25 µg/mL to select for transconjugants. After an overnight incubation at 37°C, the resulting colonies were scraped and resuspended in 25 mL of LB. The mutant suspension was diluted to an OD_600_ of 0.5 in 50 mL of LB + Kan^25^ and allowed to grow for 4h at 37°C. Glycerol was then added at 15% final concentration and 1 mL aliquots were prepared in cryovials and frozen at -80°C for later use. Genomic DNA was extracted from one aliquot with the DNeasy Blood and Tissue Kit (QIAgen). Barcode mapping to transposon insertion sites was carried out using a two-step PCR method^75^, revealing that the library contained 157,637 uniquely barcoded mutants covering 4,032 protein-coding genes out of 4,677 total, with a mean coverage of 25 unique mutants per gene.

Phage competitive fitness assays were conducted in a high-throughput manner in a 48-well culture plate. An aliquot of the library was inoculated into 25 mL of LB + Kan^25^ and grown under agitation for 4 h at 37°C to reach an OD_600_ of 0.4. Three samples of 1 mL each were immediately retrieved, centrifuged for 5 min at 15,900 x g and temporarily frozen at -20°C to serve as time-zero reference for BarSeq. The mutant library was diluted to an OD_600_ of 0.04 in 2X LB + Kan^25^ (∼ 4 × 10^7^ CFU/mL) and 350 µL was transferred to each well of a sterile 48-well culture microplate. The same volume of phage suspensions normalized at 10^9^ PFU/mL was added to the wells. The plate was then sealed with a Breathe-Easy® membrane and incubated at 37°C with orbital shaking for 16h in a BioTek 800 TS plate reader (Agilent Technologies). Surviving mutants were collected by retrieving the content of the wells and centrifuging for 5 min at 15,900 x g. Genomic DNAs were extracted using the QIAamp 96 DNA QIAcube HT Kit (QIAgen) following manufacturer’s instructions. BarSeq PCRs and Illumina sequencing were conducted as described previously^7^. Genes displaying a log2 fold-change above or equal to 4 (16-fold increase in mutant abundance compared to time zero) and a t-statistic^46^ absolute value above or equal to 5 were considered to be significantly enriched in the final pool and thus critical for phage infection in this host strain. Similarly, genes showing a log2 fold-change below or equal to -4 (16-fold decrease in mutant abundance) and a t-test absolute value above or equal to 5 were classified as low-fitness genes for which disruption increases the host phage susceptibility.

### Biological Interpretation

Shapley Additive exPlanations (SHAP) values were calculated for all features across all samples using TreeExplainer with approximate=True to handle large feature spaces efficiently^55^. Features were classified as having a positive impact (presence increased infection probability) or negative impact (presence decreased infection probability) based on mean SHAP values across all predictions.

Functional annotation was performed using multiple complementary computational approaches. Defense system classification employed Defense-Finder to identify bacterial defense mechanisms and anti-defense systems relevant to phage-bacteria interactions^57^. Functional annotation utilized eggNOG-mapper for comprehensive protein family assignment and Gene Ontology (GO) classification, creating high-confidence and low-confidence categories for phage-host interaction mechanisms including lipopolysaccharides, outer membrane proteins, flagella, pili, and enterobacterial common antigen^76^. Viral protein clustering and taxonomic classification were performed using vContact2 with Diamond similarity search and MCL clustering to identify viral protein families and evolutionary relationships^18,66,67^. Provirus detection and genomic context analysis employed geNomad to classify genomic components as viral, bacterial, or plasmid-derived, enabling identification of horizontally transferred elements and prophage regions ^24^. Protein localization was predicted using DeepLocPro, to classify predictive proteins into cellular localizations “Cell wall & surface”, “Periplasmic”, “Outer membrane”, “Cytoplasmic membrane”, “Extracellular”, and “Cytoplasmic”.

Predictive features were validated against known mediators of phage-host interactions characterized through RB-TnSeq, DubSeq, and CRISPRi experiments, which identify host genes affecting phage infection through transposon mutagenesis, overexpression, and inactivation screens^7^. This comparison revealed that the workflow successfully identified multiple screening hits, related regulators, or genes within the same operon or gene cluster as predictive features. This validation demonstrates that the approach captures biologically relevant determinants of phage infection, suggesting features with unknown relevance to phage infection may indicate novel mediators.

### Gene Neighborhood and Protein-Protein Interaction Network Analysis

To identify functional relationships between RB-TnSeq hits and computationally identified predictive features, genomic neighborhoods were analyzed based on chromosomal proximity. Genes were sorted by scaffold identifier and genomic start position within each bacterial strain. For each predictive feature, neighboring genes were identified as those within a 3-gene window (±3 genes) on the same scaffold. Distances were calculated both as the number of intervening genes and as genomic distance in base pairs. This approach accounts for operonic organization and co-regulated gene clusters that may share functional roles in phage-host interactions.

For visualization of gene neighborhoods, extended regions spanning 5 genes upstream and 5 genes downstream of genes of interest were extracted from sorted genomic coordinates. Synteny plots visualizing gene arrangement and conservation across strains were generated using clinker through the CAGECAT (Comparative Gene Cluster Analysis Toolbox) web server (https://cagecat.bioinformatics.nl/)^77^.

Functional relationships between proteins were assessed using STRING-DB (v12.0) protein-protein interaction networks ^53^. Custom interaction networks for *E. coli* strain BW25113 were generated by uploading complete proteomes to the STRING web application, enabling network construction using strain-specific locus identifiers. Network confidence scores were calculated from individual evidence channels (neighborhood, fusion, cooccurrence, homology, co-expression, experimental evidence, and database annotations) using STRING’s Bayesian integration framework with prior probability of 0.041:

For each evidence channel: score_adjusted = (score - prior) / (1 - prior)

Combined score: 1 - ∏(1 - score_adjusted_i) × (1 - prior) + prior

Protein interactions were filtered to retain high-confidence associations (combined score ≥ 600). To link RB-TnSeq hits with predictive features, networks were restricted to include only interactions where at least one partner was a computationally identified predictive feature. Features were considered functionally related to RB-TnSeq hits if they showed direct STRING interactions (combined score ≥ 600) with genes identified in genetic screens.

### Statistical Analysis

Statistical comparisons throughout employed non-parametric tests appropriate for the distribution of performance metrics. For comparisons of multiple configurations across datasets, each configuration was evaluated using 20-fold cross-validation, and configurations were ranked by mean performance on each dataset independently. The Friedman test assessed overall significance of rank differences across datasets. Pairwise comparisons of configurations used two-tailed Mann-Whitney U tests on performance scores, with effect sizes quantified using Common Language Effect Size (CLES = U/(n₁×n₂)) and Cliff’s delta to characterize the magnitude of observed differences. Correlations between dataset characteristics and model performance were assessed using Pearson correlation coefficients with two-tailed significance tests. Statistical significance was defined as p < 0.05.

Parameter optimization involved systematic evaluation of modeling configurations through multiple training iterations. Final parameter selection was based on rank-ordered performance across datasets (assessed via Friedman test), integrated with multiple independent validation criteria: (1) consistency across phylogenetically diverse datasets, (2) stability across 20-fold cross-validation, and (3) biological interpretability of predictions. This multi-criteria approach, combined with independent experimental validation (1,240 novel interactions and RB-TnSeq screening), provides robust evidence for selected parameters.

### Software Implementation and Compute Resources

Machine learning modeling was performed using the GenoPHI package available on GitHub (https://github.com/Noonanav/GenoPHI). Downstream analyses were conducted in Jupyter notebooks also available in the GitHub repository. The following software versions were used:

Core Bioinformatics Tools:

- MMSeqs2: 15.6f452
- DefenseFinder: 2.0.0
- eggNOG-mapper: 2.1.12 (with Diamond 2.1.9, eggNOG DB 5.0.2)
- vConTACT2: 0.11.3
- geNomad: 1.11.0
- PPanGGOLiN: 2.0.3
- Prodigal: 2.6.3
- Prodigal-gv: 2.11.0
- Pharokka: 1.7.3
- FastTree: 2.1.11
- DeepLocPro: 1.0.0

Python Environment:

- Python: 3.12.5
- pandas: 2.2.2, numpy: 1.26.4, scipy: 1.14.1
- scikit-learn: 1.5.2, CatBoost: 1.2.7, SHAP: 0.46.0, umap: 0.5.6, hdbscan: 0.8.33
- matplotlib: 3.9.2, seaborn: 0.13.2, plotnine: 0.13.6
- biopython: 1.84

Machine learning modeling was performed on the Lawrencium high-performance computing cluster at Lawrence Berkeley National Laboratory. Jobs were executed on lr7 partition nodes, each equipped with dual Intel Xeon Gold 6330 28-core processors (56 cores per node) and 256-512 GB memory, interconnected via HDR InfiniBand. Memory allocations ranged from 40-240 GB RAM with 8-32 CPU threads depending on dataset size and computational requirements.

Downstream analysis of results was conducted on local computational servers equipped with AMD processors (64-core AMD Opteron 6376 and 64-core AMD EPYC systems) for data processing, visualization, and statistical analysis using Jupyter notebooks and custom Python scripts.

## Acknowledgments

The authors thank members of the BRaVE Phage Foundry team, the Arkin lab, Simon Roux, Morgan N. Price, and Aeron T. Hammack for feedback and discussion, and Alexander Harms and Petr Leiman for providing phages used in experimental validation.

## Author Contributions

**A.J.C.N.**: Conceptualization, Methodology, Software, Formal Analysis, Data Curation, Writing— Original Draft, Writing—Review & Editing

**L.M.:** Conceptualization, Methodology, Investigation, Validation, Writing—Review & Editing

**E.O.R.:** Investigation, Validation, Writing—Review & Editing

**K.P.:** Software

**M.P.:** Investigation, Validation

**M.S.:** Investigation, Validation

**A.K.:** Software, Data Curation

**A.D.:** Validation, Writing—Review & Editing, Supervision

**E.G.D.:** Writing—Review & Editing, Supervision

**V.K.M.:** Conceptualization, Methodology, Writing—Review & Editing, Supervision

**A.P.A.:** Conceptualization, Methodology, Writing—Review & Editing, Supervision

## Data and Code Availability

All data and code are available through related publications and the GenoPHI GitHub repository at https://github.com/Noonanav/GenoPHI, including published interaction matrices, modeling workflows in a pip-installable Python package, analysis scripts, RB-TnSeq data, and the validation phage-host interaction matrix. The GenoPHI package is installable using ‘pip install genophì.

