## Supplementary Materials for "Phylogeny-agnostic strain-level prediction of phage-host interactions from genomes"

### 1 Supplementary Information

#### 1.1 Extended Results

##### 1.1.1 Dataset overview

The six datasets used in this study span six bacterial genera and a broad range of interaction densities and assay formats, providing a diverse foundation for workflow development and validation. The following section describes the composition, phylogenetic structure, and experimental characteristics of each dataset, and provides context for how dataset-specific properties influenced subsequent modeling decisions.

The six published phage-host interaction datasets used for model development and validation totaled 115,037 interactions, across 949 bacterial strains and 518 phages (*Table 1* / *Supp. Fig. 1-8*). The datasets needed to include both phage-host interaction matrices, where interactions were reported as, or can be converted to, binary phenotypes (infection or no-infection), and the genome sequences of associated phages and host strains (*Fig. 1A*). Four datasets were used for workflow development: an *E. coli* dataset (with a subset used for workflow development) (*Fig. 1B*)[16], a mixed *Klebsiella* spp. dataset[40], a *Klebsiella pneumoniae* dataset[17,41], and a *Pseudomonas aeruginosa* dataset (*Supp. Fig. 1*)[42]. The remaining two, a mixed *Vibrionaceae* dataset (242 *Vibrio*, 8 *Enterovibrio*, and 6 *Shewanella* strains) and a *Klebsiella* dataset, were excluded from the workflow optimization and were reserved for validation[38]. The four datasets used for workflow development totaled 27,081 interactions, including 4,304 positive interactions, and range from 23 bacterial strains by 19 phages in the *Pseudomonas* dataset to 177 strains by 94 phages in the *E. coli* dataset subset (*Fig. 1B*). The percentage of infectious (positive) interactions ranged from 2.4% to 36.2% (*Table 1*).

The *E. coli* interaction matrix was assayed on solid medium, where susceptibility was qualitatively assessed based on the clearance achieved by three phage dilutions, whereas all other datasets were assayed in liquid medium and relied on the area under the optical density curve to quantify susceptibility. These assay conditions can have a significant impact on observed interaction profiles. Plaquing assays use lawn clearance as a proxy for productive lysis but results can be confounded through enzymatic or mechanical disruption-mediated lysis (processes often referred to as lysis-from-without), abortive infection, or bacterial growth inhibition. Liquid-based assays are impacted by limit-of-detection challenges, where background limits resolution at low optical densities, and increased likelihood of phage host-range mutation[71,78]. Physical and chemical properties of assay environments also impact observed interactions, with dense biofilm-like environments of plaquing assays often resulting in increased interaction rates, in comparison to more dilute and dynamic liquid assays. It has been observed that interactions can vary based on the medium composition in liquid assays[47], but these environment-dependent phage-host interaction data are limited. For the purposes of this study, positive interactions, or those resulting in bacterial lysis or growth inhibition, were defined as any non-zero interaction in the *E. coli* dataset and based on criteria specified by the publishing authors for *Klebsiella*, *Pseudomonas*, and *Vibrionaceae* datasets.

Phylogenetic trees based on amino acid alignments of conserved genes were generated for each dataset[61,62]. These show host-strain diversity within individual datasets and shared phylogenetic clades between the two Klebsiella datasets, which are the only datasets covering the same host phylogenies (*Supp. Fig. 2-6*). A phage gene-sharing network representing all datasets showed that phages across datasets and host phylogenies were interspersed within phage clusters (*Supp. Fig. 7-8*)[18]. This suggested that phages with similar gene content, possibly representing structural or mechanistic similarities, are capable of infecting diverse hosts. These results suggest that gene content-based features could likely be shared across datasets and that pooling of datasets may improve model performance.

##### 1.1.2 Workflow development and optimization

Developing a phylogeny-agnostic prediction workflow required systematic evaluation of a large number of interdependent modeling decisions, spanning genomic representation, feature selection, model architecture, and training strategy. The following section describes the rationale and outcomes of this optimization process, with a focus on the key decisions that most substantially influenced final model performance.

Our objective was to develop a modeling workflow for strain-level prediction of phage-host interactions applicable to phage therapy design, industrial biotechnology, and biological research. The workflow needed to meet three key criteria: 1) predict interactions between individual bacterial strain-phage pairs; 2) be independent of host and phage phylogeny without requiring prior knowledge of genetic determinants of phage-host interaction; 3) enable predictions for novel bacterial strains or phages based on genomic information alone. These criteria distinguish our workflow from existing strategies that frequently assign host taxonomies to phages from an existing host database, often at higher taxonomic ranks, preventing these models from being used to evaluate specific phage-host interactions[18,22,25-36]. Modeling strategies that do predict one-to-one phage-host interactions: 1) use prior mechanistic knowledge, limiting applicability to single species with known mediators of phage infection[16,17], 2) do not directly address feature assignment and prediction for new strains, impacting their use in targeting new host strains[16], or 3) generate phage-specific models, enabling only the prediction of infection by an existing set of phages[13,16]. Additionally, we considered secondary criteria, including biological interpretability of predictive features and the applicability of resulting predictions to phage selection and phage cocktail design.

We performed an extensive modeling parameter optimization experiment, evaluating over 200 parameter settings and training more than 13.2 million individual predictive models. This involved optimization of feature engineering and selection strategies, predictive model training workflows, and feature assignment to novel bacterial strains and phages. Specifically, workflow optimization involved testing genomic representations including protein-family clustering strategies (*Supp. Fig. 9-10*) and amino acid (AA) *k*-mer lengths (*Supp. Fig. 11*)[64,66], modeling algorithms (*Supp. Fig. 12-13*)[68,69], feature selection algorithms (*Supp. Fig. 14*)[55,69], ensemble-learning approaches (*Supp. Fig. 15-19*), feature table filtering and engineering strategies, and training strategies (*Supp. Fig. 20*)[69], such as class weights and clustering strategies for train-test splits[48,69,79].

###### *1.1.2.1 Genomic Representation Strategies*

To generate phylogeny-independent numerical features representing bacterial and phage genomes, we used binary presence/absence matrices of either protein families, in a pangenome-like structure, or amino acid *k*-mers, allowing us to capture sequence variants and key residues. These feature generation workflows were run independently, with whole bacterial strain and phage proteomes represented as protein families or *k*-mers, or sequentially, representing only proteins belonging to predictive protein families as *k*-mers. We tested MCL and MMSeqs2 clustering algorithms for protein-family construction, including varied inflation values (the MCL argument controlling clustering stringency) and MMSeqs2 sequence identity and coverage thresholds (*Supp. Fig. 9-10*)[64,66]. We then compared eight machine learning algorithms across four development datasets, identifying CatBoost gradient-boosted decision trees as the most consistently high-performing model (*Supp. Fig. 12-13*). We implemented recursive feature elimination (RFE) to identify predictive subsets of feature tables of up to 18,236 protein family-based features or 165,938 *k*-mer features and benchmarked this strategy against five alternative feature selection methods (*Supp. Fig. 14* / *Supp. Table 1 and 5*). Feature selection was chosen over dimension reduction to enable both simplified feature assignment to novel bacterial strains and phages, and biological interpretation of predictive features.

###### *1.1.2.2 Feature Selection and Phylogenetic Filtering*

We tested whether performance was improved by filtering phylogenetically linked features prior to feature selection, decreasing the risk of overfitting and increasing the likelihood of identifying features linked to mechanism, rather than phylogeny. To accomplish this, bacterial strains and phages are clustered based on feature content (hierarchical or HDBSCAN clustering[48]) and features unique to a single cluster were removed. This filtering highlights a trade-off between generalizability and the capacity to identify strain- or clade-specific genetic features that may impact phage infection. For example, genes acquired through horizontal gene transfer and maintained only in a small number of closely related bacterial strains were removed. Filtering of phylogenetically-linked features improved prediction performance in the *E. coli* and *Klebsiella*-2 datasets, while this filtering slightly decreased performance in the smaller *Pseudomonas* and *Klebsiella*-1 dataset. We ultimately chose to maintain this filtering as it improved performance of the most robust models and leads to the removal of phylogenetically-linked features that might be indicators of a specific clade, rather than mechanism. When applied on complete datasets, filtering removed from 0.27% to 18.23% of bacterial features and 3.45% to 40.41% of phage features. Removed features contained significantly more protein families per feature (Mann-Whitney U test; $p < 0.001$), including 5.44% to 42.84% of bacterial protein families and 20.98% to 72.39% of phage protein families (*Supp. Fig. 21-22* / *Supp. Table 5-6*). This trend is expected given that filtered features often represent groups of many shared protein families within a single clade, making remaining features easier to associate with specific protein families for biological interpretation. One of the results of this filtering is that mechanistically relevant features unique to a single clade, (i.e. acquired through horizontal gene transfer) were removed from the feature table, potentially limiting within-clade discriminatory power. However, even if mechanistically relevant, these protein families may be collapsed with many other clade-specific families, complicating biological interpretation without improving model performance in the largest datasets.

###### *1.1.2.3 Ensemble Learning and Training Strategy*

We also explored the impact of iterative feature selection and modeling across various train-test splits, selecting features based on recurrence across feature selection iterations and using an ensemble-learning approach for final predictions (*Supp. Fig. 15-19*). Train-test splitting based on bacterial strain or phage clusters was also tested, forcing models to learn features predictive of interaction in distinct phylogenetic clades. This also prevents very similar strains from being present in training and testing datasets, limiting overfitting during feature selection and model training.

###### *1.1.2.4 Final Configuration Testing*

Finally, we tested the impact of *k*-length on model performance for *k*-mer-based feature tables. To evaluate computational feasibility of *k*-mer-based representations of full proteomes, we quantified the number of resulting features (unique presence-absence patterns) from *k* lengths of 3 to 6. While the counts remain below the theoretical $20^k$ maximum, the number of features scales aggressively from between 56,592 and 165,938 at $k = 4$ to between 437,634 and 1,764,951 at $k = 5$, in the largest 4 datasets. As a result, *k* values above 4 create an untenable feature space for iterative modeling, leading us to select *k* values to 3-4 for full proteome representations (*Supp. Fig. 21-22* / *Supp. Table 5-6*). Values of *k* from 3-15 were tested when filtering to predictive proteins only, which significantly decreased *k*-mer diversity enabling larger *k* values.

To quantify the overall importance of each methodological decision, we calculated the performance delta (Δ) between the worst- and best-performing configuration within each parameter based on Matthews Correlation Coefficient (MCC). This showed that modeling algorithm (Δ MCC = 0.365), training strategies (Δ MCC = 0.203), and feature selection method (Δ MCC = 0.184) had the largest impact on performance, while genomic representation parameters, including protein-family clustering thresholds (Δ MCC = 0.072) and *k*-mer length (Δ MCC = 0.020) showed limited effects (*Table 2* / *Supp. Fig. 20*).

To select a final parameter set, we employed rank-order analysis across datasets, with performance differences assessed using Friedman tests and pairwise Mann-Whitney U comparisons. This identified configurations that consistently maximized Matthews Correlation Coefficient (MCC) across phylogenetically diverse bacterial genera. The optimized workflow uses MMSeqs2-based protein family clustering (sequence identity 0.4, coverage 0.8), CatBoost gradient-boosted decision tree modeling, 25 rounds of RFE-based feature selection, 50 modeling iterations for ensemble-learning, and a *k*-mer length of 4 in the standalone *k*-mer workflow and 6 when combining with predictive protein families. To address dataset imbalance (2.1 %-36.2% positive interactions) and minimize overfitting, we implemented inverse-frequency-based phage-specific class weights, hierarchical clustering of bacterial strains (20 clusters) during train-test split, and feature filtering that maintains only features present across multiple strain clusters, removing features with strong phylogenetic linkage (*Supp. Fig. 20*).

##### 1.1.3 Model Performance

Having established an optimized modeling workflow, we evaluated predictive performance across all datasets and model configurations through nested cross-validation experiments. The following section describes performance across three biologically motivated prediction scenarios, the impact of dataset size and composition on model reliability, and the effect of combining datasets within and across bacterial genera.

When evaluating performance, we tested three model configurations that reflect possible applications in the laboratory or the clinic. These configurations dictate train-test splitting paradigms during feature selection and training, and were tested in 20-fold nested cross-validation experiments. The first configuration represents the most likely application, where users want to select a phage from an existing phage set (“phage bank”) that infects a novel bacterial strain, such as in the context of phage therapies or microbiome engineering. For cross-validation of these models, 10% of bacterial strains are excluded from the entire feature selection and modeling workflow and features are assigned to these strains for prediction with resulting models. During feature selection and training, strains are split across train and test datasets, whereas phages are maintained. The second mirrors this strategy, leaving out 10% of phages from the modeling workflow and predicting infection of a characterized set of bacterial strains by previously unseen phages. This reflects a scenario where researchers want to select a suitable host-strain for a target phage. Here phages are split across train and test datasets, with strains maintained. The final cross-validation workflow leaves out 10% of strains and 10% of phages, enabling predictions in completely unseen bacteria-phage pairs. For all validation sets, features were assigned to held-out genomes using similarity search against training protein clusters (MMSeqs2, sequence identity ≥0.4, coverage ≥0.8), ensuring feature assignment reflected realistic inference conditions. Model performance varied across datasets for all model configurations (*Supp. Fig. 23*), but with consistent relationships between dataset size and model performance (*Supp. Fig. 24-26*).

First, we evaluated the performance impact of genomic representation strategies by comparing 20-fold nested cross-validation performance of protein-family and *k*-mer-based models using the primary model configuration. Here we observed no significant differences in performance between models, suggesting that all representations captured sufficient biological information for predicting phage-host interactions (*Fig. 2A*). Although *k*-mer-based representations provide sequence-level detail and may offer insights into specific functional residues or motifs, they present competing constraints associated with specificity and multidimensionality at increasing values of *k*. While biological sequence space is sparse, the unique presence-absence pattern space for $k > 4$ remains extremely high-dimensional (*Supp. Fig. 21 / Supp. Table 5*). Excluding the small Pseudomonas dataset, feature counts range from 437,634 in the *Klebsiella*-1 dataset to 1,764,951 in the *Vibrionaceae* dataset at $k = 5$. This presents significant computational challenges for iterative ensemble-learning workflows. At $k = 4$, where feature counts in the four largest datasets are in a computationally feasible range (56,592 to 165,938), *k*-mer-based features frequently identify sequences shared across multiple genes. The median number of genes represented by each 4-mer is 6-7 in bacterial strains, compared to a median of 1 in phages, across all five datasets. While $k = 4$ represents a practical compromise between specificity and dimensionality, this multi-gene redundancy complicates the association of predictive features with specific genes and functional domains (*Supp. Fig. 22 / Supp. Table 6*) without offering a performance advantage over protein families (*Fig. 2A*). For these reasons, we focused on protein family-based models, which achieved comparable performance and provided a more streamlined path to biological interpretation and validation. Despite this, we believe that alternative representations could provide value in other contexts, such as with small genomes or when a specific subset of relevant genes has already been identified, and should be considered when training predictive models.

When predicting phages infecting novel bacterial strains, simulating the process of designing phage therapies, area under the receiver operating characteristic curve (AUROC) ranges from 0.67 to 0.94, Matthews Correlation Coefficient (MCC) from 0.13 to 0.54, and normalized area under the precision-recall curve (AUPR) of 0.07 to 0.60 (*Fig. 2B / Supp. Fig. 23A / Table 2*). For *E. coli*, our models achieved an AUROC of 0.87, closely matching the performance reported by the original study (AUROC = 0.86) despite fundamental differences in approach. Notably, the published method required phage-specific models and host features based on known *E. coli* mediators of phage infection[16], while our approach achieved comparable performance without these constraints, enabling application to novel phage-host systems. Similarly, when predicting hosts for novel phages, AUROC ranged from 0.69 to 0.93, MCC from 0.18 to 0.68, and normalized AUPR of 0.10 to 0.87 (*Supp. Fig. 23B / Supp. Table 3*). The final model configuration, predicting interaction between unseen bacteria-phage pairs, was most sensitive to dataset size, yielding AUROC values of 0.58 to 0.92, MCC of 0.07 to 0.44, and normalized AUPR of 0.03 to 0.58 (*Supp. Fig. 23C / Supp. Table 4*).

Across all cross-validation experiments, we observed significant positive correlations between model performance and dataset characteristics, including both total and infectious interaction counts (*Supp. Fig. 24-26*). When predicting phages infecting unseen hosts, performance increased with dataset size, showing Pearson correlation coefficients of $r = 0.60$ for AUROC and $r = 0.47$ for MCC (both $p < 1\times10^{-6}$). Positive interaction counts also correlated with performance, with $r = 0.28$ for AUROC ($p = 0.0057$) and $r = 0.36$ for MCC ($p = 2.0\times10^{-4}$). By contrast, the proportion of positive interactions showed a marginally significant negative correlation with AUROC ($r = -0.26$, $p = 0.0081$) and no correlation with MCC ($r = -0.20$, $p = 0.472$). This slight decrease in performance with an increased proportion of infectious interactions likely reflects characteristics of our specific datasets, including the high performance of the large *Vibrionaceae* dataset (63,488 interactions; AUROC = 0.94 and MCC = 0.54) with only 2.2% positive interactions and the variable performance of the small *Pseudomonas* dataset (437 interactions; AUROC = 0.75 and MCC = 0.33) with 36.2% positive interactions (*Supp. Fig. 24*). The *Klebsiella*-1 dataset was the lowest performing across all model configurations, with AUROC of 0.67, 0.69, and 0.58 in configurations 1, 2, and 3, respectively. This is the second smallest dataset used in model development, with only 3,658 interactions and 4.9% positive interactions. This seems to indicate a data structure threshold, where models fail to generalize at this sparsity and size. *Klebsiella*-2, with 6,348 total interactions and only 2.4% positive interactions, performs significantly better (AUROC of 0.88, 0.81, and 0.68), suggesting a threshold of near 5000 interactions for low density datasets. Overall, these results suggest that a sufficient total interaction count can mitigate the negative performance impact of dataset imbalance. Standard deviation in MCC and AUROC across rounds of cross-validation also decreased with dataset size, showing increasing stability and reliability in larger datasets (*Supp. Fig. 23*).

###### *1.1.3.1 Cross-Genus Model Performance*

A key question for the practical utility of this framework is whether models trained on one set of host-phage pairs can inform predictions for phylogenetically distinct systems. The following section evaluates the impact of combining datasets within and across genera, and tests the limits of cross-genus generalizability through leave-one-group-out cross-validation.

We next evaluated the impact of training models on combined datasets both across phylogenetically diverse hosts and within single host genera, including two categories of performance test. The first involved training models on combined datasets with validation sets representing fractions of each dataset, allowing us to investigate whether including additional data from unrelated datasets improves within-dataset prediction. Notably, this performance test enables phages to be maintained across training and validation sets, with predictions being made on unseen bacterial strains. The cross-genus combination included *E. coli*, *Klebsiella*, and *Pseudomonas* datasets (*Supp. Fig. 28*), while the within-genus combination merged the two *Klebsiella* datasets (*Supp. Fig. 27*). Combining *Klebsiella* datasets led to slight but significant performance improvements in the *Klebsiella*-2 dataset ($p = 0.024$) and a non-significant improvement in the *Klebsiella*-1 dataset ($p = 0.398$) compared to single-dataset models. This represents MCC increases of 0.04 and 0.07 over single-dataset models, despite the relatively small 10,006 interaction combined dataset (*Fig. 2D* / *Supp. Fig. 27A-B*). In these combined *Klebsiella* models, there was no significant correlation between the strain-level increase in predictive performance and the proximity of the closest relative added by the additional data, suggesting that this performance improvement cannot be explained by the addition of highly similar bacterial strains (*Supp. Fig. 27C-D*). This indicates a more complex relationship between genotype and phage-host interaction phenotype than phylogenetic relatedness, or may highlight the trade-off between generalizability through filtering of phylogenetically linked features and within-clade discrimination based on narrowly conserved clade-specific features. The AUROC of 0.812 of this combined model also nearly matches the performance of models reported by a previous study (AUROC = 0.818), but whose method is heavily dependent on *Klebsiella*-specific knowledge using only *K*-locus genes and phage tail fibers[17]. This again indicates our workflow can provide comparable performance to alternative workflows without requiring phylogeny-specific knowledge of the mediators of phage-host interactions. Unlike within-genus combinations, merging cross-genus datasets did not significantly improve performance of any dataset (*Supp. Fig. 28A*). This response likely relates to the sharing of predictive features across datasets and the requirement for a minimum set of shared features for accurate predictions. Jaccard distance analysis based on feature content revealed clear separations between host phylogenies (*Supp. Fig. 28B*), with more overlap observed between phages (*Supp. Fig. 28C*), suggesting that sufficient feature overlap is required for performance gains through dataset combination. The fraction of predictive features that were shared across genera were often associated with either core microbial metabolic processes or mobile genetic elements (MGEs), suggesting horizontal transmission of phage defense systems or previous cross-genera phage infection (*Supp. Fig. 29*). These shared features may represent interesting conserved mechanisms of phage infection and defense, but their apparent limited impact on predictive performance indicates that phylogeny-specific genetic features may play a more critical role.

The second category of combined-dataset performance tests involved leave-one-group-out (LOGO) cross-validation at the genus-level. In this context, models were trained on combinations of *E. coli*, *Klebsiella*, and *Pseudomonas* datasets, with one of these taxonomic groups reserved for validation in each cross-validation run. This cross-validation format presents an additional challenge over combined dataset models described above, as predictions are made on unseen bacterial strains and phages, representing the third model configuration, showing the lowest performance within datasets (*Supp. Fig. 23C / Supp. Table 4*). Here we observed significantly decreased performances, with AUROC only slightly above random for all datasets, with 0.60, 0.55, and 0.57 in *E. coli*, *Klebsiella*, and *Pseudomonas* datasets, respectively (*Supp Fig. 30*). This likely results either from the biological reality that phage-host determinants are largely genus-specific or the same limitation of protein-family-based features described previously, where limited feature overlap was observed across host phylogenies. This is combined with the additional predictive challenge of requiring models to learn cross-phylogeny phage representations. Despite limited improvement above random predictions, ROC curves indicated that the highest confidence positive predictions are slightly enriched in true positive predictions (*Supp Fig. 30*). This observation led us to evaluate whether these models could still be useful in selecting phages infecting bacterial strains in genera for which we have no knowledge of phage-host interaction profiles. For this analysis, we compared the proportion of bacterial strains for which the top 5 phages ranked by predicted infection probability included at least one infectious phage to the proportion of bacterial strains infected by at least one phage from a random selection of 5 phages. This comparison is valid, as we are assuming no knowledge of phage host range in the left-out dataset. Here, we observed a significant increase in the proportion of suitable phages identified by models in *E. coli* and *Klebsiella* datasets, allowing the identification of at least one infectious phage in 87% of the *E. coli* dataset and 52% of the *Klebsiella* dataset, representing 1.45 and 1.74-fold increases over random, respectively (*Supp Fig. 30*). No increase was observed in the *Pseudomonas* dataset, with at least one infectious phage being identified for 90% of strains by both model and random selection. The results highlight both the limited cross-genus applicability of these models, but also their potential for increasing the rate of identifying suitable phages for a given target, even out of distribution. This could decrease the experimental overhead for researchers or clinicians attempting to identify phages targeting under-studied host phylogenies.

##### 1.1.4 Modeling Error Analysis

Understanding where and why models fail is as important as characterizing overall performance, particularly for a workflow intended to guide experimental decisions. The following section systematically characterizes prediction errors in the *E. coli* dataset, examining bias patterns, the relationship between phylogenetic isolation and strain- or phage-level performance, and the fidelity of predicted versus observed interaction similarities between strain and phage pairs.

###### *1.1.4.1 Prediction Bias and Strain Susceptibility*

First, an analysis of overall model bias revealed consistent over-prediction bias, with 58 bacterial strains showing systematic over-prediction (>20% deviation from observed infection rates) compared to only 11 under-predictors. This bias correlated negatively with strain susceptibility ($r = -0.207$, $p = 8.3 \times 10^{-5}$), being most pronounced in bacterial strains with narrow susceptibility profiles, consistent with the positive correlation between strain susceptibility and model performance ($r = 0.310$, $p = 2.3 \times 10^{-9}$) (*Supp. Fig. 31-32*). This indicates that a larger proportion of interactions are falsely predicted to be infectious in bacterial strains with narrow susceptibility profiles. Several factors likely contribute to this behavior. First, by nature of their narrow susceptibility, the dataset includes few phages capable of infecting similar host strains, preventing models from learning relevant feature sets. Whether these host strains are truly only susceptible to a small set of phages, possibly through extensive defense mechanisms or distinct receptor variants, cannot be distinguished from the possibility that the tested phage set is not representative of appropriate phage diversity for these strains. This highlights the importance of including a phylogenetically broad set of phages in experimental datasets.

###### *1.1.4.2 Phylogenetic Isolation and Performance*

To explore this further, we investigated the impact of phylogenetic distribution of bacterial strains and phages on prediction accuracy. For all bacterial genomes, we constructed phylogenetic trees based on core-genome concatenated marker genes and examined how prediction performance varied across phylogenetic space (*Supp. Fig. 2-6*). For each strain, we calculated the mean distance to its five closest phylogenetic neighbors as a measure of phylogenetic isolation. Surprisingly, we found no consistent relationship between phylogenetic isolation and strain-level prediction performance (MCC) ($r = -0.055$, $p = 0.60$), indicating that the presence of closely related bacterial strains in the training data does not necessarily improve prediction accuracy (*Supp. Fig. 32*). This further supports our observations when modeling combined *Klebsiella* datasets, suggesting that mechanisms of phage susceptibility are not directly linked to phylogeny or are not necessarily vertically inherited. In contrast to this, a significant negative correlation was observed between phage performance and phylogenetic isolation ($r = -0.425$, $p = 1.9 \times 10^{-5}$), where proteomic equivalence was used as a metric of phage genomic similarity (*Supp. Fig. 32*). This relationship indicates that models performed less well when predicting infection by more distantly-related phages, highlighting the importance of phage diversity in interaction datasets and the possible performance benefit of phage phylogenetic redundancy. This could also be a limitation of the modeling dataset that included 402 *E. coli* strains, in comparison to only 94 phages.

##### *1.1.4.3 Interaction Profile Similarity Analysis*

To systematically quantify where models fail to recognize distinguishing genetic features dictating bacterial infection patterns, we calculated pairwise Jaccard similarities between bacterial strain pairs and phage pairs based on their experimental interaction profiles and compared these to model-predicted interaction similarities across 66,795 bacterial strain pairs and 4,371 phage pairs. This calculation evaluates whether bacterial strain pairs and phage pairs with similar interaction profiles are predicted to behave similarly. The result is a sensitive metric of strain- or phage-level predictive performance, where a one-to-one relationship indicates perfect accordance and a large distance from this line indicates discordance and the inability of the model to capture critical information. Given our bacterial strain-based cross-validation design, where all phages appear in both training and test sets, phage-phage similarity correlations were expectedly high ($r = 0.893$, $p < 1 \times 10^{-12}$), indicating models effectively capture phage-specific infectivity patterns (*Supp. Fig. 35C-D*). In contrast, strain-strain similarity correlations were moderate ($r = 0.387$ ($p < 1 \times 10^{-12}$) for all bacterial strains, $r = 0.493$ ($p < 1 \times 10^{-12}$) for strains infected by ≥20 phages) (*Supp. Fig. 35A-B*). The improved correlation for broadly susceptible bacterial strains matches the observation that models better capture similarity patterns among these strains. Analysis of extreme discordance cases, where shared interactions differed significantly from shared predicted interactions, revealed fundamental model failures in capturing relevant biological information. We identified strain pairs where models predicted nearly identical interaction patterns (similarity > 0.9) despite minimal biological overlap (true similarity < 0.3), exemplified by *E. coli* BL21 comparisons to *E. coli* strains ECOR22, ECOR42 and ECOR54. This indicates the model's failure to capture BL21's distinct phage susceptibility patterns, which result from its laboratory-adapted genetic background including truncated lipopolysaccharide and altered outer membrane protein composition compared to wild-type bacterial strains. Conversely, strain pairs like *E. coli* IAI45 compared to *E. coli* strains ECOR57, ECOR59, and ECOR71 showed high biological similarity (>0.7) but low predicted similarity (<0.3), indicating failure to recognize shared mediators of infection. This information may help identify bacterial strains and phages with abnormal interaction patterns, possibly uncovering novel mediators or sequence variants of biological interest.

Despite these limitations, high-confidence errors (false-negatives with predicted likelihood of infection of 0-0.1 and false-positives with predicted likelihood of infection of 0.9-1) were rare, totaling only 1.5% of predictions (*Supp. Fig. 31*). This suggests most failures occur in uncertain boundary cases rather than confident misclassifications. Calibration curves showing predicted vs. observed infection rates (*Supp. Fig. 34*) and Brier scores (*Table 1* / *Supp. Table 3-4*) support this observation, with scores under 0.15 for all datasets and model configurations except the 437-interaction *Pseudomonas* dataset. In all but the low-performing *Klebsiella*-1 dataset, high-confidence predictions are suitable targets for experimental validation, with observed infection rates between 80-95% when model prediction confidence ≥ 0.9. High AUROC values further demonstrate that models successfully rank phages by infection probability (*Fig. 2B / Supp Fig. 23*). Together, these results demonstrate that this workflow identifies predictive genetic features from whole genomes and that prediction confidence can effectively guide phage selection for engineering and therapeutic applications.

##### 1.1.5 Phage Selection and Phage Cocktail Design

Translating probabilistic model outputs into actionable phage selection decisions requires an explicit cocktail design strategy. This section compares candidate cocktail design approaches across datasets and cocktail sizes, evaluates their performance relative to promiscuity-based baselines, and identifies a strategy suitable for downstream applications.

Previous studies have demonstrated that effective phage cocktail design requires balancing phage infectivity with phage diversity, such that selected phages maximize strain coverage while reducing redundancy and potentially limiting escape [13,15,16,37]. Guided by these principles, we evaluated cocktail design approaches that combined two phage ranking strategies with three high-level selection frameworks. Phages were ranked either by model-predicted interaction confidence or by phage promiscuity, defined as the fraction of training strains infected by each phage. These rankings were then used in one of three selection frameworks: (1) rank-based selection of the top *n* phages, (2) hierarchical clustering followed by selection of the highest-ranked phage from each of *n* clusters, and (3) density-based clustering using HDBSCAN followed by selection of the highest-ranked phage from each cluster and construction of cocktails from the *n* cluster representatives with the highest ranking scores, with remaining positions filled by the next-highest-ranked phages when fewer than *n* clusters were identified (*Fig. 3A-B / Supp. Fig. 36-37*)[48]. For clustering-based frameworks, phages were grouped in one of two representation spaces. First, phages were clustered by activity group using strain-phage interaction profiles, following the "activity group" framework proposed by Keith et al. (2024)[13]. Second, phages were clustered using pangenome-derived feature representations containing predictive features identified through the feature selection workflow. Together, these factors yielded 10 unique cocktail design conditions.

Phage selection and cocktail performance was measured as the proportion of target bacterial strains for which at least one selected phage showed activity against the target strain. Although this metric does not directly test whether a phage combination suppresses resistance emergence or prevents escape in combination therapy, it does quantify whether cocktail design improves the probability of selecting at least one active phage while mitigating uncertainty in model-guided prioritization. We evaluated single-phage selections alongside 3-phage and 5-phage cocktails, comparing model-guided strategies against baseline cocktails composed of the most promiscuous phages. Performance was evaluated across 20-fold cross-validation, maintaining the same strain holdout strategy used for individual interaction predictions.

Cocktail performance varied across datasets, but with model-based ranking outperforming promiscuity-based ranking in all cases. Within ranking strategies, clustering phages into “activity groups” outperformed pangenome-based representations in 9 of 12 design workflows with model-based ranking and 10 of 12 workflows based on promiscuity, suggesting interaction matrix-based activity groups better capture phage functional clades. Focusing on these interaction matrix-based phage representations, clustering using HDBSCAN outperformed hierarchical clustering in all model-based ranking strategies and 4 of 6 promiscuity-based ranking strategies (*Supp. Fig. 36-37*). In select cases, selection of highest-ranked phages without clustering led to the highest coverage, but with a small increase over clustering-based methods. We maintained clustering in our workflow, as our current metric does not evaluate the possibility of cross-resistance, meaning that matching performance with phage diversity is preferable for downstream application. Finally, we compared the published promiscuity-based design, using hierarchical clustering into activity groups, with model-based design using HDBSCAN clustering of the interaction matrix (the best-performing strategy). For single-phage selection, models identified active phages for 66.9% of E. coli strains, 66.7% of Klebsiella strains, and 39.4% of Vibrionaceae strains, equating to 1.1-fold, 3.1-fold, and 2.7-fold increases over promiscuity-based cocktails, respectively. For 5-phage cocktails, models selected active phages for 97.5%, 87.8%, and 57.5% of E. coli, Klebsiella, and Vibrionaceae strains, respectively, representing 1.2-fold, 1.7-fold, and 1.5-fold improvements over promiscuity-based cocktails. Despite strong performance, the relative increase in bacterial strain coverage over promiscuity-based selection in the *E. coli* dataset was smaller than in *Klebsiella* and *Vibrionaceae*. This reflects dataset sparsity and structure, where *E. coli* phage DIJ07_P2 shows activity against 65.4% of tested strains, setting the minimum performance of promiscuity-based cocktails, in comparison to the most broadly active phages in *Klebsiella* and *Vibrionaceae* datasets that interact with 13.8% and 14.8% of bacterial strains, respectively. Overall, the results demonstrate that model-guided cocktail design outperforms promiscuity-based methods and that incorporating mechanistic diversity into model-guided phage selection maintains high performance while potentially reducing the risk of resistance evolution in a cocktail setting.

##### 1.1.6 Experimental Validation

Cross-validation performance demonstrates that models learn generalizable features within a given dataset, but external validation in independently generated experimental data is necessary to confirm that predictions reflect genuine biological signal rather than dataset-specific artifacts. The following section describes three complementary validation experiments: a 1,300-interaction spotting assay matrix testing model predictions on previously unseen phages, a genome-wide RB-TnSeq genetic screen linking computationally identified predictive features to experimentally confirmed host determinants of phage infection, and a cross-dataset *Klebsiella* validation testing model generalizability outside of *E. coli*.

###### *1.1.6.1 E. coli Interaction Matrix Validation*

We selected 25 *E. coli* strains belonging to the ECOR collection that were included in the published dataset (*E. coli* ECOR13 - ECOR38) and predicted interaction of these strains with a set of 52 previously unseen phages belonging to the BASEL phage collection[5,16,49-51]. Spotting assays were then performed with these phages on all 25 *E. coli* strains, resulting in 1,300 tested interactions of which 60 (4.6%) did not show a clear phenotype (*Fig. 4A*). This assay format mimics experimental protocols used during generation of the published training dataset, allowing models to predict analogous phenotypes. However, this proxy of phage infection is known to result in observed clearance phenotypes that may not represent full productive lysis cycles, through enzymatic or mechanical disruption-based lysis, or growth inhibition. To further validate our observed phenotypes, we selected a representative 20 phages and 12 *E. coli* strains (18.5% of dataset) (*Supp. Fig. 39*), ensuring each phage showed activity against at least two of the selected bacterial strains, and performed an efficiency of plating (EOP) experiment by plaquing eight 10-fold serial dilutions of each phage on all strains (*Supp. Fig. 40*). We observed that only 4 of 145 interactions (2.8%) scored as "0" showed a clearance phenotype at any dilution, with these representing four of the lowest observed EOP values, and all 82 interactions scored as "1" showed a clearance phenotype on at least one dilution. The 13 filtered interactions with unclear phenotypes in single-plaquing assays, showed mixed results in EOP experiments with 7 of 13 showing clearance phenotypes (*Supp. Fig. 41*). These results confirmed that our assigned scores reproducibly represent true clearance phenotypes across phage dilutions.

###### *1.1.6.2 Cross-Dataset Klebsiella Validation*

To evaluate model generalizability outside of *E. coli*, we identified a published *Klebsiella* dataset including 3,318 interactions across 68 *Klebsiella* strains and 54 phages. Models were trained combining *Klebsiella*-1 and *Klebsiella*-2 datasets and used to predict interactions on this third dataset. This model configuration already demonstrated low performance in small *Klebsiella*-1 and *Klebsiella*-2 datasets, with AUROC of 0.58 and 0.68, respectively, indicating that the requirement to learn both conserved bacterial strain and phage representations likely makes models more sensitive to dataset size. Despite this, models showed an AUROC of 0.63 when predicting on the new *Klebsiella* dataset, showing that comparable performance is maintained even in this more challenging predictive task (*Supp. Fig. 30*).

###### *1.1.6.3 Predictive Feature Validation with RB-TnSeq*

To validate the biological relevance of the predictive features, we performed an RB-TnSeq genetic screening experiment in *E. coli* ECOR27, which was included in both the modeling dataset and our interaction matrix described above. In these experiments, pooled genome-wide mutant libraries are subjected to phages, allowing us to identify host factors associated with phage infection through the increased fitness of their mutants (*Supp. Fig. 44*)[7]. Although transposon mutagenesis-based libraries only enable screening of non-essential genes, preventing the identification of essential genes associated with phage infection, this still represents a valuable set of experimentally confirmed genes for cross-comparison against predictive features. The RB-TnSeq comparison was conducted at three levels of stringency to account for the co-inheritance of functionally related genes and the limitations of protein family-based feature assignment. Gene neighborhood (3 genes upstream and downstream) and STRING-DB-based functional relationships were included because functionally related genes are often co-inherited or co-maintained within genomes[80,81] making models incapable of distinguishing them during feature selection, as these have equal predictive power. This analysis demonstrated that the predictive modeling workflow was particularly effective at identifying features associated with cell-wall biosynthesis pathways, including LPS, O-antigen, cellulose, and capsule biosynthesis, whereas porins were often represented by regulators or adjacent genes.

To further characterize the biological relevance of predictive features, we compared their annotations to published RB-TnSeq, DubSeq, and CRISPRi experiments[7]. This revealed multiple additional screening hits or related genes successfully identified as predictive features, including *aes*, *bcsA* (*bcsA* / *bcsE* / *bcsF* / *bcsQ*), *ycgF* (*bluR* anti-repressor), *uvrY* (*csrA* regulator), *cyaA*, *dsbB* (*dsbA*), *fhuA*, *ydiV* (negative regulator of *flhD*), *gadC*, *glmS*, *macB* (*macA*), *malT* (*malX* / *malQ*), *mdtB* (*mdtB* / *mdtF*), *nfrA*, *ompC*, *ompF*, *ompT*, *ptsG*, *rpiR* (negative regulator of *rpiB*), *cytR* (negative regulator of *tsx*), *waaY*, *yhbX*, and *yphG*. Of these, multiple are porins known to be the primary phage receptors (*fhuA, nfrA*, *ompC*, *ompF*, *ompT*) or important contributors to *E. coli* outer-membrane structure (*bcsA*, *waaY*).

##### 1.1.7 Biological Interpretation

Beyond predictive accuracy, a key goal of this framework is to identify specific genetic features that mechanistically explain phage-host interaction outcomes. The following section examines the top predictive features in the *E. coli* dataset, linking them to known and potentially novel determinants of phage susceptibility through functional annotation, gene neighborhood analysis, and STRING-DB-based pathway inference.

Analysis of the top 25 most predictive features in the *E. coli* dataset revealed various functional categories associated with known phage-host interaction mechanisms (*Fig. 5A*). Three of these features were directly associated with RB-TnSeq data in *E. coli* ECOR27, relating to cellulose, O-antigen, and LPS biosynthesis. Of the unvalidated features, the two that related directly to cell wall biosynthesis showed varied effects, with presence of the O-antigen biosynthesis gene *rfbB* increasing infection probability and the absence of the peptidoglycan biosynthesis regulator *agaR* decreasing infection probability. Two features associated with genes of viral origin could either promote or inhibit infection, potentially reflecting either superinfection exclusion mechanisms or indicators of host susceptibility. Seven features were related to mobile genetic elements and two with components of RM systems, showing varying impacts on infection likelihood. A total of eight features showed no clear relationship to known phage-host interaction mechanisms, potentially representing novel mediators requiring further investigation.

To explore possible phage-infection mechanisms associated with the top two predictive features in the *E. coli* dataset, we employed the same STRING-DB and gene neighborhood analysis used to link predictive features to RB-TnSeq hits. We found that *goaG* (also referred to as *puuE*) and *gabT* are associated with putrescine catabolism (*Fig. 5B*). Putrescine has been shown to prevent bacteriophage replication in *Pseudomonas aeruginosa* and may act as a host signaling molecule during phage infection[58], indicating that putrescine-mediated inhibition of phage replication may be conserved in *E. coli*. A gene neighborhood analysis showed that *ybdK*, which is encoded in 392 of 402 strains, is located adjacent to the outer membrane protein *fepA*, a known phage receptor (*Fig. 5C*)[59]. Although *fepA* is still present in strains missing *ybdK*, this may indicate changes in regulation that impact receptor expression. Although these phenotypes would require experimental validation, these analyses show that modeling features are potentially valuable tools in hypothesis generation and the discovery of novel mediators of phage infection.

### 2 Supplementary Tables

*Supplementary Table 1.* Filtering of phylogenetically-linked features based on strain and phage clustering.

| Dataset | Type | Features Retained | Features Removed | % Features Removed | Protein Families Maintained | Protein Families Removed | % PFs Removed |
|---|---|---|---|---|---|---|---|
| E. coli | Strain | 18052 | 1744 | 8.81 | 23736 | 4686 | 16.49 |
| E. coli | Phage | 184 | 102 | 35.66 | 657 | 874 | 57.09 |
| Klebsiella-1 | Strain | 3532 | 79 | 2.19 | 9320 | 1272 | 12.01 |
| Klebsiella-1 | Phage | 156 | 36 | 18.75 | 235 | 586 | 71.38 |
| Klebsiella-2 | Strain | 5416 | 221 | 3.92 | 11011 | 1122 | 9.25 |
| Klebsiella-2 | Phage | 116 | 35 | 23.18 | 198 | 494 | 71.39 |
| Pseudomonas | Strain | 1114 | 3 | 0.27 | 7335 | 422 | 5.44 |
| Pseudomonas | Phage | 28 | 1 | 3.45 | 162 | 43 | 20.98 |
| Vibrionaceae | Strain | 7988 | 1781 | 18.23 | 12668 | 9496 | 42.84 |
| Vibrionaceae | Phage | 634 | 430 | 40.41 | 759 | 1990 | 72.39 |

*Supplementary Table 2.* Impact of modeling parameters on predictive performance.

| Parameter | Δ MCC | Δ AUROC | Configurations per dataset |
|---|---|---|---|
| Modeling Algorithm | 0.365 | 0.148 | 8 |
| Training Strategies (class weights / feature filtering) | 0.203 | 0.110 | 14 |
| Feature Selection Algorithm | 0.184 | 0.086 | 5 |
| Modeling / Feature Selection Iterations | 0.124 | 0.069 | 156 |
| MMSeqs2 Clustering Threshold | 0.072 | 0.031 | 25 |
| *k* length | 0.066 | 0.020 | 13 |

*Supplementary Table 3.* Model performance across datasets when predicting infection of known bacterial strains by unseen phages.

| Dataset | AUROC | MCC | Normalized AUPR | Brier score |
|---|---|---|---|---|
| *Pseudomonas* | 0.925 | 0.682 | 0.869 | 0.115 |
| *Vibrionaceae* | 0.924 | 0.487 | 0.659 | 0.030 |
| *E. coli* | 0.870 | 0.510 | 0.563 | 0.146 |
| *Klebsiella*-2 | 0.814 | 0.294 | 0.312 | 0.037 |

| | | | | |
|---|---|---|---|---|
| *Klebsiella*-1 | 0.687 | 0.178 | 0.102 | 0.088 |

*Supplementary Table 4.* Model performance across datasets when predicting interactions between unseen bacteria-phage pairs.

| Dataset | AUROC | MCC | Normalized AUPR | Brier score |
|---|---|---|---|---|
| *Vibrionaceae* | 0.920 | 0.436 | 0.515 | 0.025 |
| *E. coli* | 0.812 | 0.410 | 0.381 | 0.134 |
| *Pseudomonas* | 0.766 | 0.283 | 0.576 | 0.238 |
| *Klebsiella*-2 | 0.675 | 0.194 | 0.147 | 0.022 |
| *Klebsiella*-1 | 0.577 | 0.068 | 0.032 | 0.064 |

*Supplementary Table 5.* Whole-proteome *k*-mer feature counts at *k* values of 3-6.

| Feature Type | Dataset | k3 | k4 | k5 | k6 |
|---|---|---|---|---|---|
| Strain | *E. coli* | 299 | 27,332 | 1,174,637 | 1,501,337 |
| | *Klebsiella*-1 | 2 | 17,802 | 419,335 | 326,402 |
| | *Klebsiella*-2 | 4 | 16,298 | 432,756 | 335,268 |
| | *Vibrionaceae* | 11 | 37,900 | 1,569,348 | 2,420,929 |
| | *Pseudomonas* | 2 | 5,025 | 34,900 | 29,445 |
| Phage | *E. coli* | 6234 | 68,650 | 29,279 | 7,277 |
| | *Klebsiella*-1 | 6091 | 53,426 | 18,299 | 6,021 |
| | *Klebsiella*-2 | 6087 | 40,294 | 10,669 | 3,657 |
| | *Vibrionaceae* | 7815 | 128,038 | 195,603 | 42,901 |
| | *Pseudomonas* | 1888 | 4,074 | 805 | 242 |
| Total | *E. coli* | 6533 | 95,982 | 1,203,916 | 1,508,614 |
| | *Klebsiella*-1 | 6093 | 71,228 | 437,634 | 332,423 |
| | *Klebsiella*-2 | 6091 | 56,592 | 443,425 | 338,925 |
| | *Vibrionaceae* | 7826 | 165,938 | 1,764,951 | 2,463,830 |
| | *Pseudomonas* | 1890 | 9,099 | 35,705 | 29,687 |

*Supplementary Table 6.* Gene specificity in whole-proteome *k*-mer features at *k* values of 3-6.

| Feature Type | Dataset | k-length | Mean Genes | Median Genes | Std |
|---|---|---|---|---|---|
| Strain | E. coli | k3 | 170.93 | 126 | 154.28 |
| | E. coli | k4 | 9.94 | 6 | 11.61 |
| | E. coli | k5 | 1.63 | 1 | 1.26 |
| | E. coli | k6 | 1.07 | 1 | 0.33 |
| | Klebsiella-1 | k3 | 187.17 | 131 | 182.94 |
| | Klebsiella-1 | k4 | 10.91 | 6 | 14.12 |
| | Klebsiella-1 | k5 | 1.74 | 1 | 1.57 |
| | Klebsiella-1 | k6 | 1.09 | 1 | 0.46 |
| | Klebsiella-2 | k3 | 187.87 | 132 | 184.09 |
| | Klebsiella-2 | k4 | 10.94 | 6 | 14.18 |
| | Klebsiella-2 | k5 | 1.74 | 1 | 1.53 |
| | Klebsiella-2 | k6 | 1.08 | 1 | 0.36 |
| | Vibrio | k3 | 168.51 | 125 | 149.78 |
| | Vibrio | k4 | 9.91 | 6 | 11.04 |
| | Vibrio | k5 | 1.61 | 1 | 1.16 |
| | Vibrio | k6 | 1.06 | 1 | 0.30 |
| | Pseudomonas | k3 | 226.24 | 137 | 263.86 |
| | Pseudomonas | k4 | 13.61 | 7 | 21.99 |
| | Pseudomonas | k5 | 2.08 | 1 | 2.43 |
| | Pseudomonas | k6 | 1.14 | 1 | 0.52 |
| Phage | E. coli | k3 | 3.51 | 2 | 3.38 |
| | E. coli | k4 | 1.18 | 1 | 0.49 |
| | E. coli | k5 | 1.01 | 1 | 0.12 |
| | E. coli | k6 | 1.00 | 1 | 0.04 |
| | Klebsiella-1 | k3 | 2.98 | 2 | 2.63 |
| | Klebsiella-1 | k4 | 1.14 | 1 | 0.42 |
| | Klebsiella-1 | k5 | 1.01 | 1 | 0.11 |
| | Klebsiella-1 | k6 | 1.00 | 1 | 0.03 |
| | Klebsiella-2 | k3 | 2.65 | 2 | 2.16 |
| | Klebsiella-2 | k4 | 1.12 | 1 | 0.38 |
| | Klebsiella-2 | k5 | 1.01 | 1 | 0.11 |
| | Klebsiella-2 | k6 | 1.00 | 1 | 0.04 |
| | Vibrio | k3 | 2.81 | 2 | 2.58 |
| | Vibrio | k4 | 1.13 | 1 | 0.41 |
| | Vibrio | k5 | 1.01 | 1 | 0.10 |
| | Vibrio | k6 | 1.00 | 1 | 0.04 |

| | | | | | |
|---|---|---|---|---|---|
| | Pseudomonas | k3 | 2.82 | 2 | 2.23 |
| | Pseudomonas | k4 | 1.14 | 1 | 0.40 |
| | Pseudomonas | k5 | 1.01 | 1 | 0.12 |
| | Pseudomonas | k6 | 1.00 | 1 | 0.05 |

*Supplementary Table 7*. Phage list used in experimental validation.

| Phage ID | Full Phage Name | Genome Accession | Genome Size (bp) | Interaction matrix | RB-TnSeq |
|---|---|---|---|---|---|
| T2 | Escherichia phage T2 | MH751506.1 | 163832 | | x |
| P1vir | Escherichia phage P1vir | NC_005856.1 | 94800 | | x |
| phi92 | Escherichia phage phi92 | FR775895.2 | 148612 | | x |
| Bas01 | Escherichia phage AugustePiccard | MZ501051 | 50126 | x | |
| Bas02 | Escherichia phage JeanPiccard | MZ501080 | 47149 | x | |
| Bas03 | Escherichia phage JulesPiccard | MZ501087 | 47731 | x | |
| Bas04 | Escherichia phage FritzSarasin | MZ501069 | 51777 | x | |
| Bas05 | Escherichia phage PeterMerian | MZ501101 | 47775 | x | x |
| Bas06 | Escherichia phage KarlJaspers | MZ501090 | 51474 | x | |
| Bas07 | Escherichia phage JakobBernoulli | MZ501079 | 51129 | x | x |
| Bas08 | Escherichia phage DanielBernoulli | MZ501059 | 50731 | x | |
| Bas09 | Escherichia phage PaulSarasin | MZ501099 | 51285 | x | x |
| Bas10 | Escherichia phage IsaakIselin | MZ501077 | 49911 | x | x |
| Bas12 | Escherichia phage BrunoManser | MZ501053 | 49298 | x | |
| Bas13 | Escherichia phage LeonhardEuler | MZ501092 | 50192 | x | |
| Bas14 | Escherichia phage TheodorHerzl | MZ501107 | 43627 | x | |
| Bas15 | Escherichia phage PaulFeyerabend | MZ501097 | 44550 | x | |
| Bas16 | Escherichia phage GeorgBuechner | MZ501070 | 44295 | x | |
| Bas17 | Escherichia phage KarlBarth | MZ501088 | 43104 | x | |
| Bas18 | Escherichia phage Oekolampad | MZ501095 | 44882 | | x |
| Bas19 | Escherichia phage ChristophMerian | MZ501057 | 58073 | x | x |
| Bas20 | Escherichia phage FritzHoffmann | MZ501068 | 59834 | x | |
| Bas21 | Escherichia phage GottfriedDienst | MZ501071 | 56589 | x | x |
| Bas22 | Escherichia phage KurtStettler | MZ501091 | 58567 | x | |
| Bas25 | Escherichia phage VogelGryff | MZ501110 | 58342 | x | x |

| | | | | | |
|---|---|---|---|---|---|
| Bas27 | Escherichia phage TrudiGerster | MZ501108 | 114179 | x | |
| Bas28 | Escherichia phage IrmaTschudi | MZ501076 | 115446 | x | |
| Bas29 | Escherichia phage SuperGirl | MZ501105 | 110821 | x | |
| Bas31 | Escherichia phage DaisyDussoix | MZ501058 | 113532 | x | |
| Bas33 | Escherichia phage HildyBeyeler | MZ501074 | 111607 | x | |
| Bas34 | Escherichia phage SelmaRatti | MZ501103 | 107080 | x | |
| Bas35 | Escherichia phage WilhelmHis | MZ501113 | 166861 | x | |
| Bas36 | Escherichia phage Paracelsus | MZ501096 | 167732 | x | |
| Bas37 | Escherichia phage KarlGJung | MZ501089 | 167832 | x | x |
| Bas38 | Escherichia phage AugustSocin | MZ501052 | 167411 | x | x |
| Bas39 | Escherichia phage FriedrichMiescher | MZ501066 | 169509 | x | x |
| Bas41 | Escherichia phage FriedrichZschokke | MZ501067 | 165574 | x | |
| Bas42 | Escherichia phage AndreasVesalius | MZ501050 | 168255 | x | |
| Bas43 | Escherichia phage TadeuszReichstein | MZ501106 | 166115 | x | |
| Bas44 | Escherichia phage AdolfPortmann | MZ501046 | 166773 | x | |
| Bas45 | Escherichia phage PaulHMueller | MZ501098 | 170053 | x | x |
| Bas46 | Escherichia phage ChristianSchoenbein | MZ501056 | 169031 | x | |
| Bas47 | Escherichia phage AlbertHofmann | MZ501047 | 168968 | x | |
| Bas48 | Escherichia phage CarlMeissner | MZ501054 | 137623 | x | x |
| Bas49 | Escherichia phage EmilHeitz | MZ501062 | 139980 | x | |
| Bas51 | Escherichia phage WalterGehring | MZ501111 | 140659 | x | |
| Bas52 | Escherichia phage RudolfGeigy | MZ501102 | 137380 | x | |
| Bas54 | Escherichia phage MaxBurger | MZ501093 | 136346 | x | |
| Bas56 | Escherichia phage AlexBoehm | MZ501048 | 135345 | x | |
| Bas57 | Escherichia phage MaxTheCat | MZ501094 | 135061 | x | |
| Bas58 | Escherichia phage HeinrichReichert | MZ501073 | 136906 | x | |
| Bas59 | Escherichia phage EduardKellenberger | MZ501061 | 131526 | x | |
| Bas60 | Escherichia phage PaulScherrer | MZ501100 | 150425 | x | |
| Bas61 | Escherichia phage EmilieFrey | MZ501063 | 146666 | x | x |
| Bas63 | Escherichia phage JohannRWettstein | MZ501086 | 87100 | x | x |
| Bas64 | Escherichia phage JeanTinguely | MZ501081 | 39451 | x | |
| Bas65 | Escherichia phage JacobBurckhardt | MZ501078 | 39451 | x | |

| | | | | | |
|---|---|---|---|---|---|
| Bas67 | Escherichia phage ErnstBeyeler | MZ501064 | 39315 | x | x |
| Bas68 | Escherichia phage CarlSpitteler | MZ501055 | 39466 | x | x |
| Bas69 | Escherichia phage AlfredRasser | MZ501049 | 70849 | x | x |

### 3 Supplementary Figures

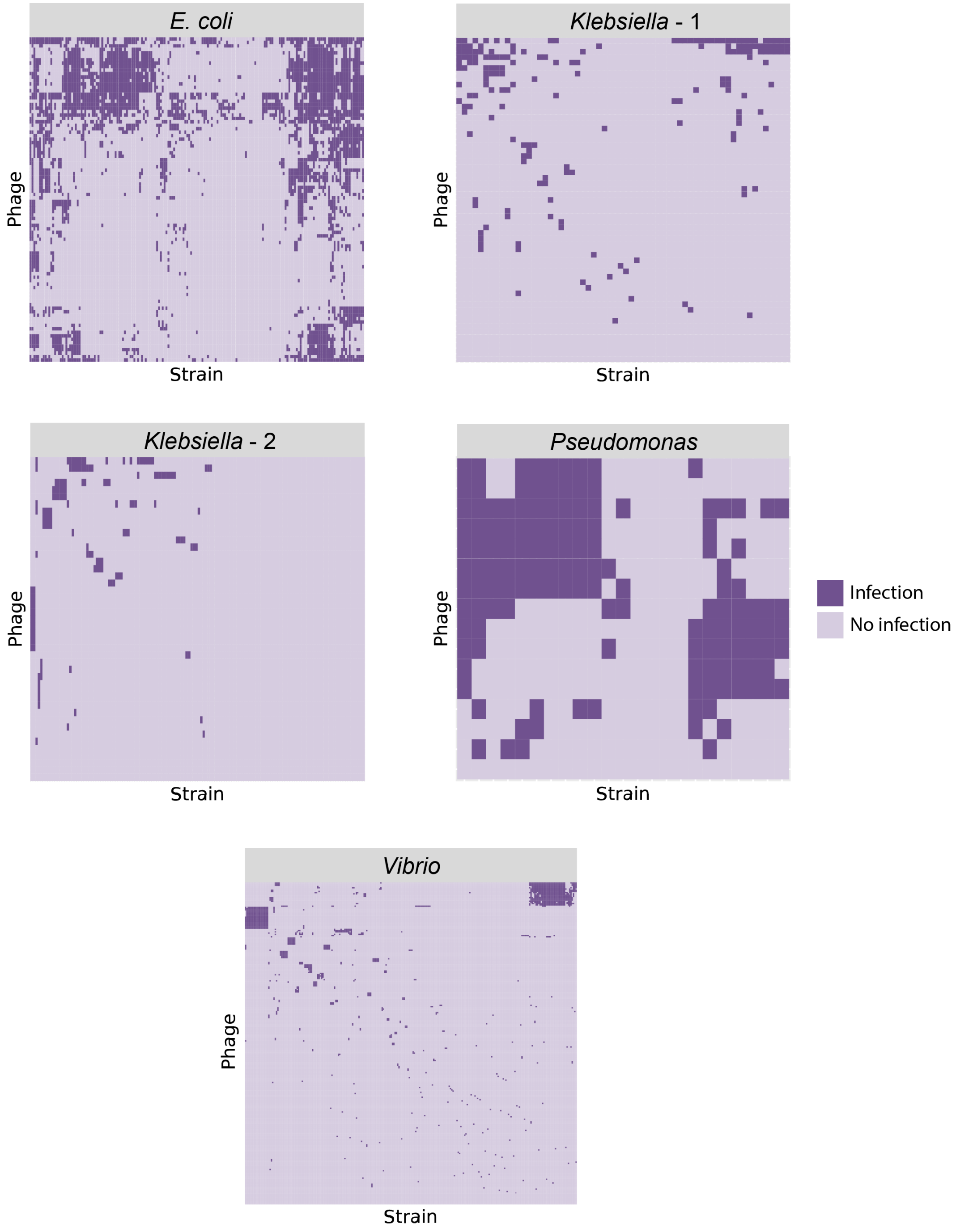

*Supplementary Figure 1. Phage-host interaction matrices.* The interaction matrices for five published datasets. Clusters of bacterial strains and phages indicate conserved infection patterns within these groups. Dark purple represents interactions resulting in infection and light purple indicates interaction that did not result in infection.

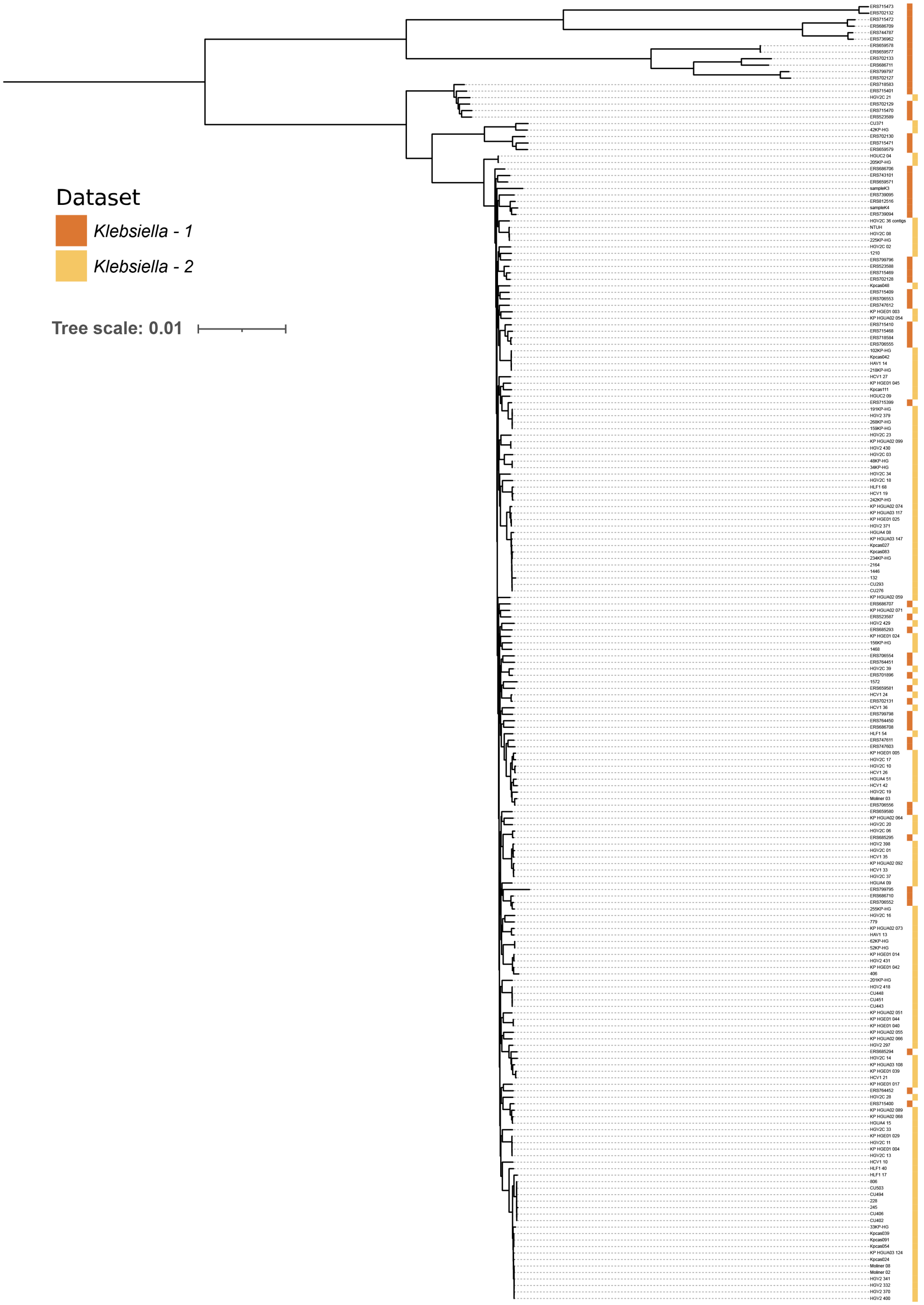

*Supplementary Figure 2. Klebsiella dataset tree.* Phylogenetic tree based on core-genome concatenated marker genes alignments. Dark orange indicates mixed *Klebsiella* spp. dataset and light orange indicates *Klebsiella pneumoniae* dataset.

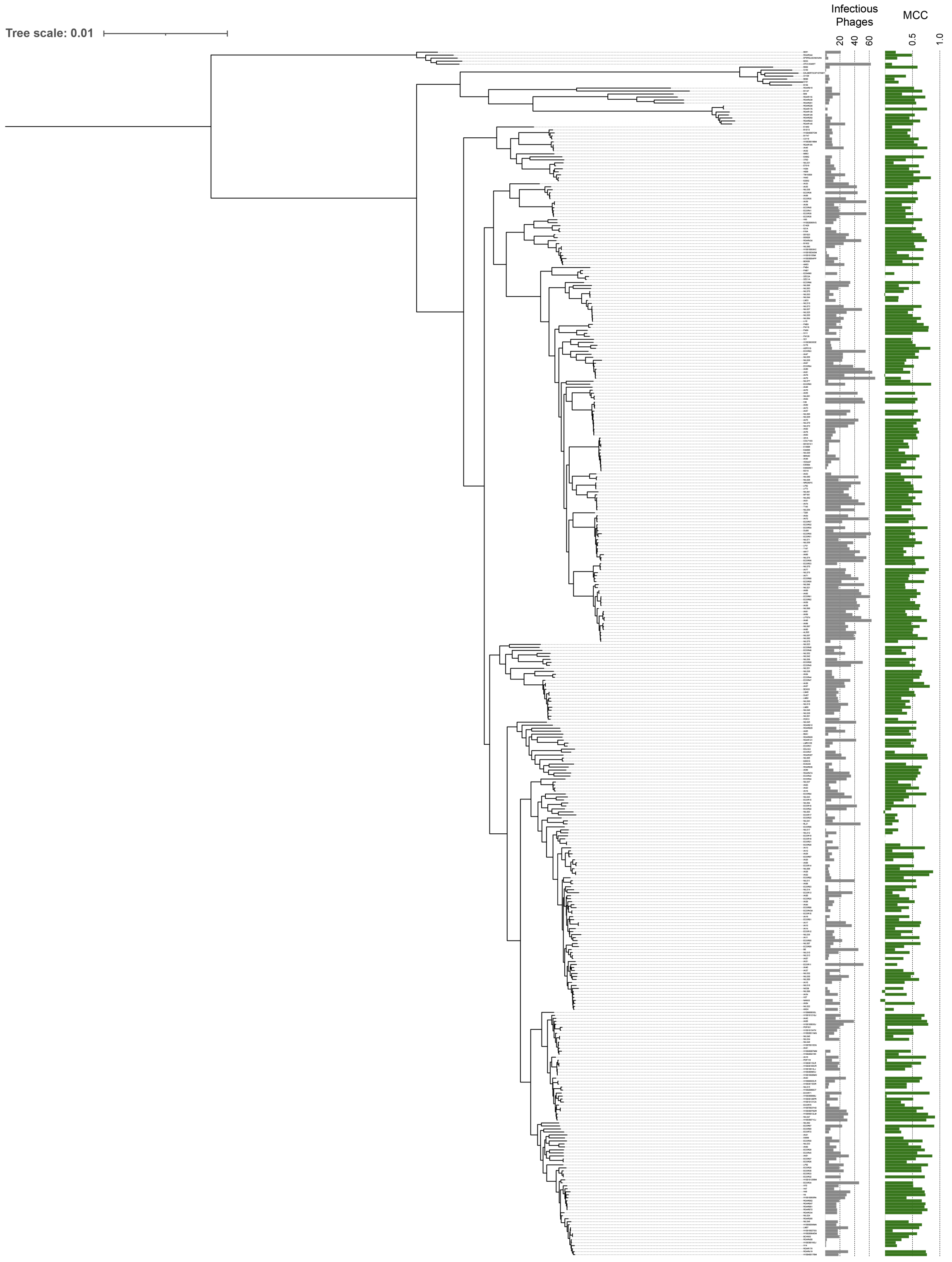

*Supplementary Figure 3. E. coli dataset tree.* Phylogenetic tree based on core-genome concatenated marker genes alignments. Grey bars indicate number of infectious phages out of 94-phage set. Green bars indicate prediction performance in 20-fold cross-validation for models with bacterial strain-based train-test splitting (unseen strains).

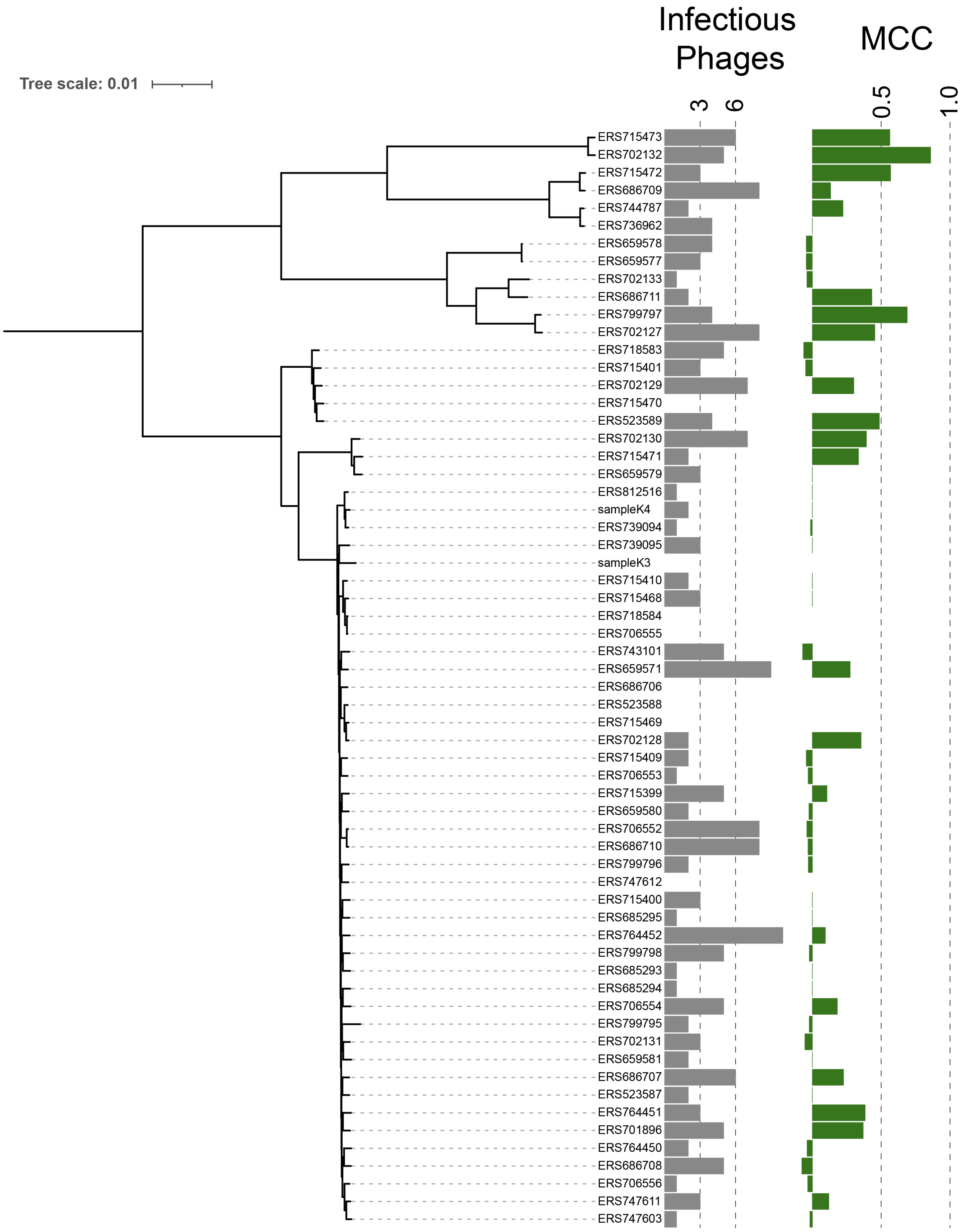

*Supplementary Figure 4. Klebsiella-1 dataset tree* Phylogenetic tree based on core-genome concatenated marker genes alignments. Grey bars indicate number of infectious phages out of 59-phage set. Green bars indicate prediction performance in 20-fold cross-validation for models with bacterial strain-based train-test splitting (unseen strains).

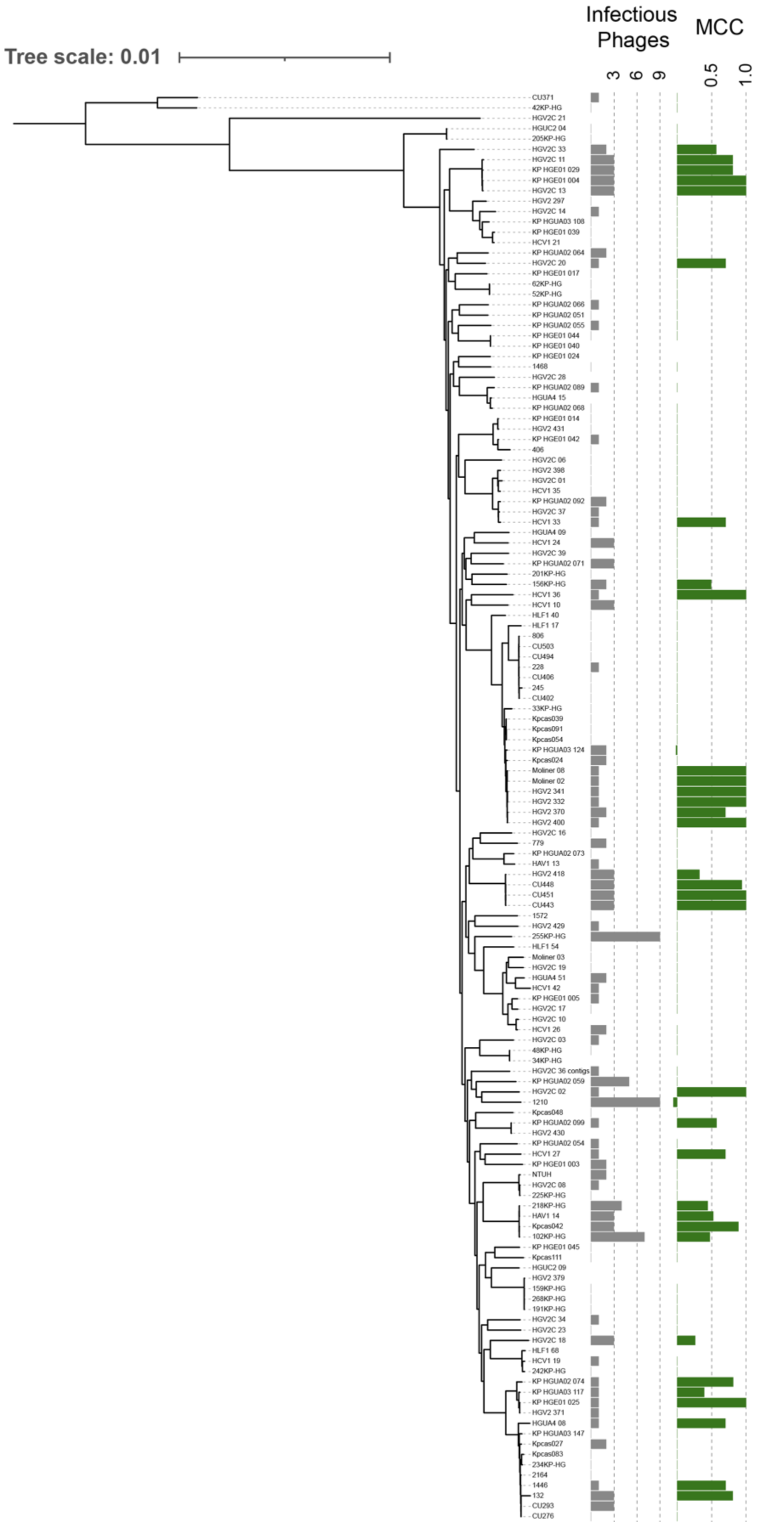

*Supplementary Figure 5. Klebsiella-2 dataset tree.* Phylogenetic tree based on core-genome concatenated marker genes alignments. Grey bars indicate number of infectious phages out of 46-phage set. Green bars indicate prediction performance in 20-fold cross-validation for models with bacterial strain-based train-test splitting (unseen strains).

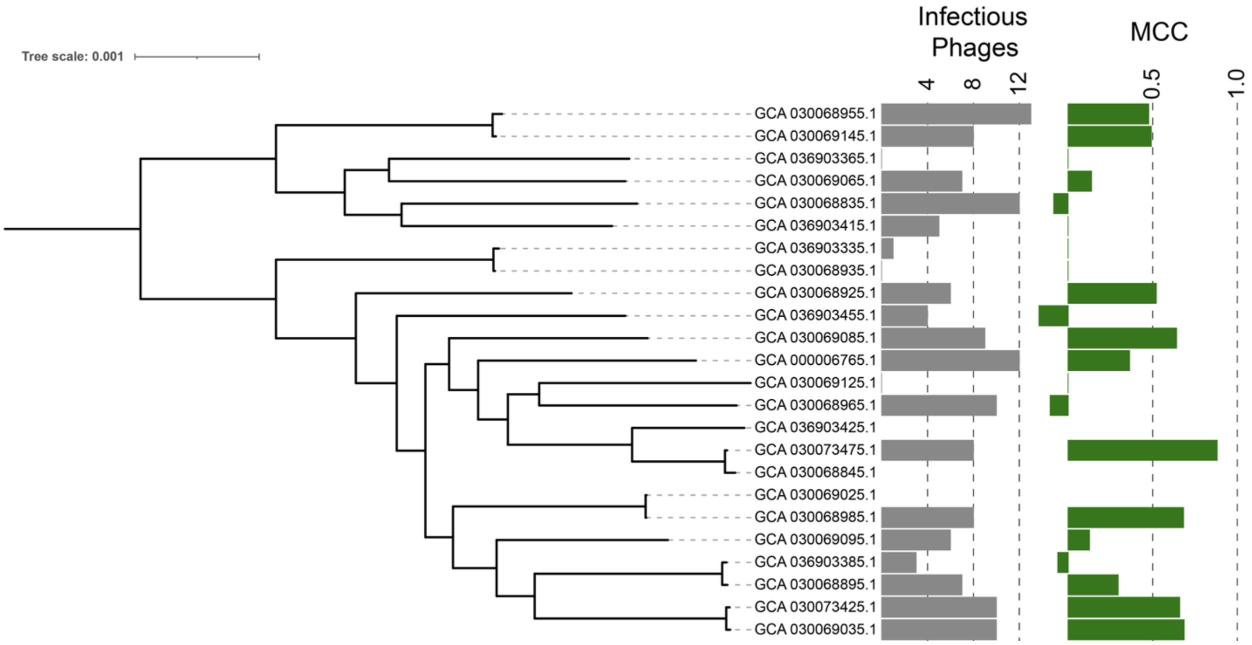

*Supplementary Figure 6. Pseudomonas aeruginosa dataset tree.* Phylogenetic tree based on core-genome concatenated marker genes alignments. Grey bars indicate number of infectious phages out of 19-phage set. Green bars indicate prediction performance in 20-fold cross-validation for models with bacterial strain-based train-test splitting (unseen strains).

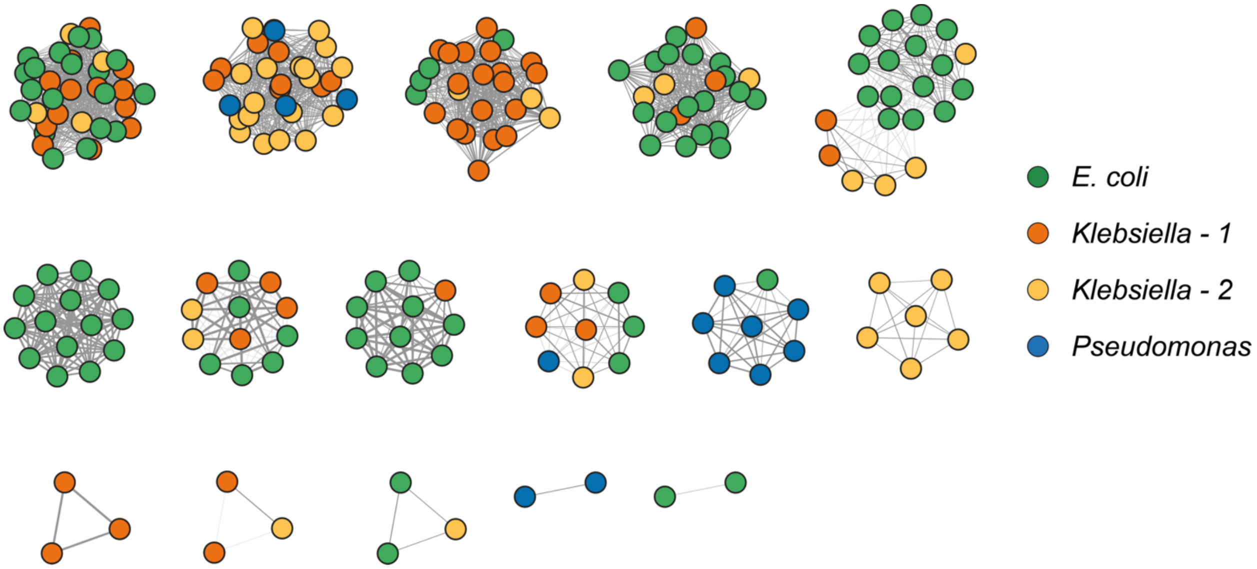

*Supplementary Figure 7. Phage gene sharing network for development datasets.* Gene sharing network shows distribution of phages across datasets (color). Edges indicate vConTACT2 gene sharing similarity metric.

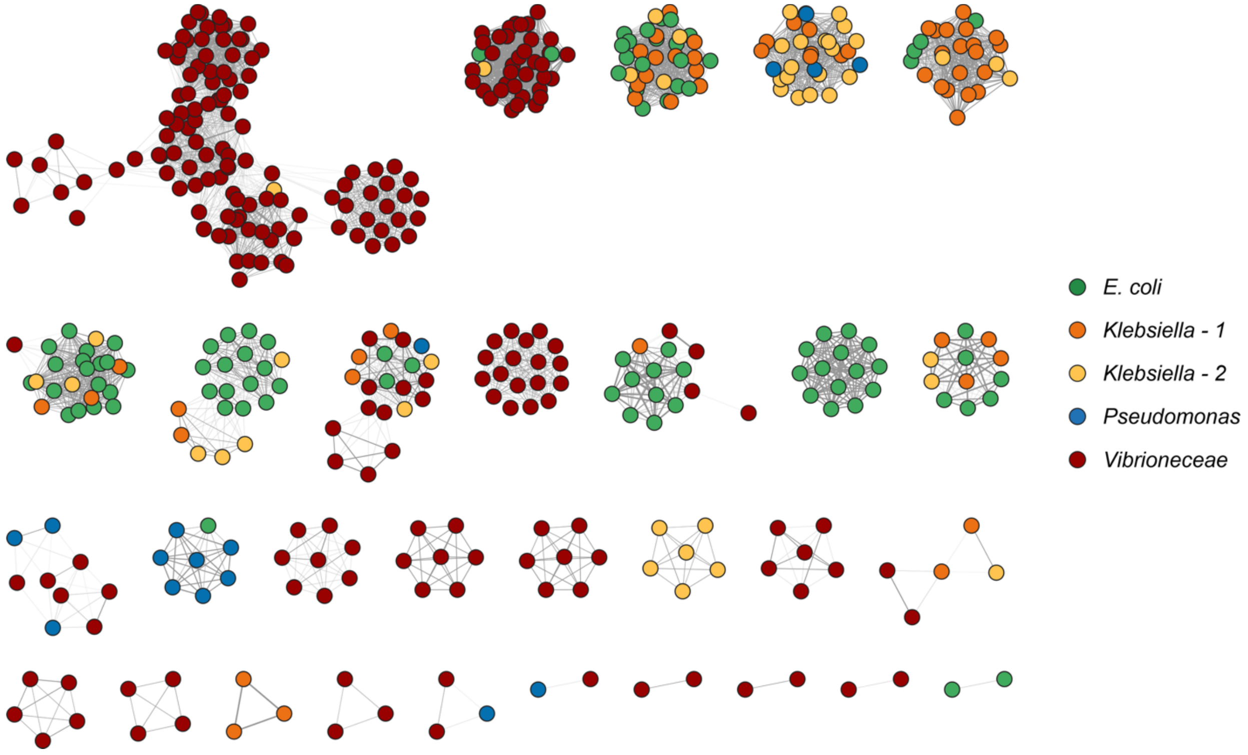

*Supplementary Figure 8. Phage gene sharing network for public datasets.* Gene sharing network shows distribution of phages across datasets (color). Edges indicate vConTACT2 gene sharing similarity metric.

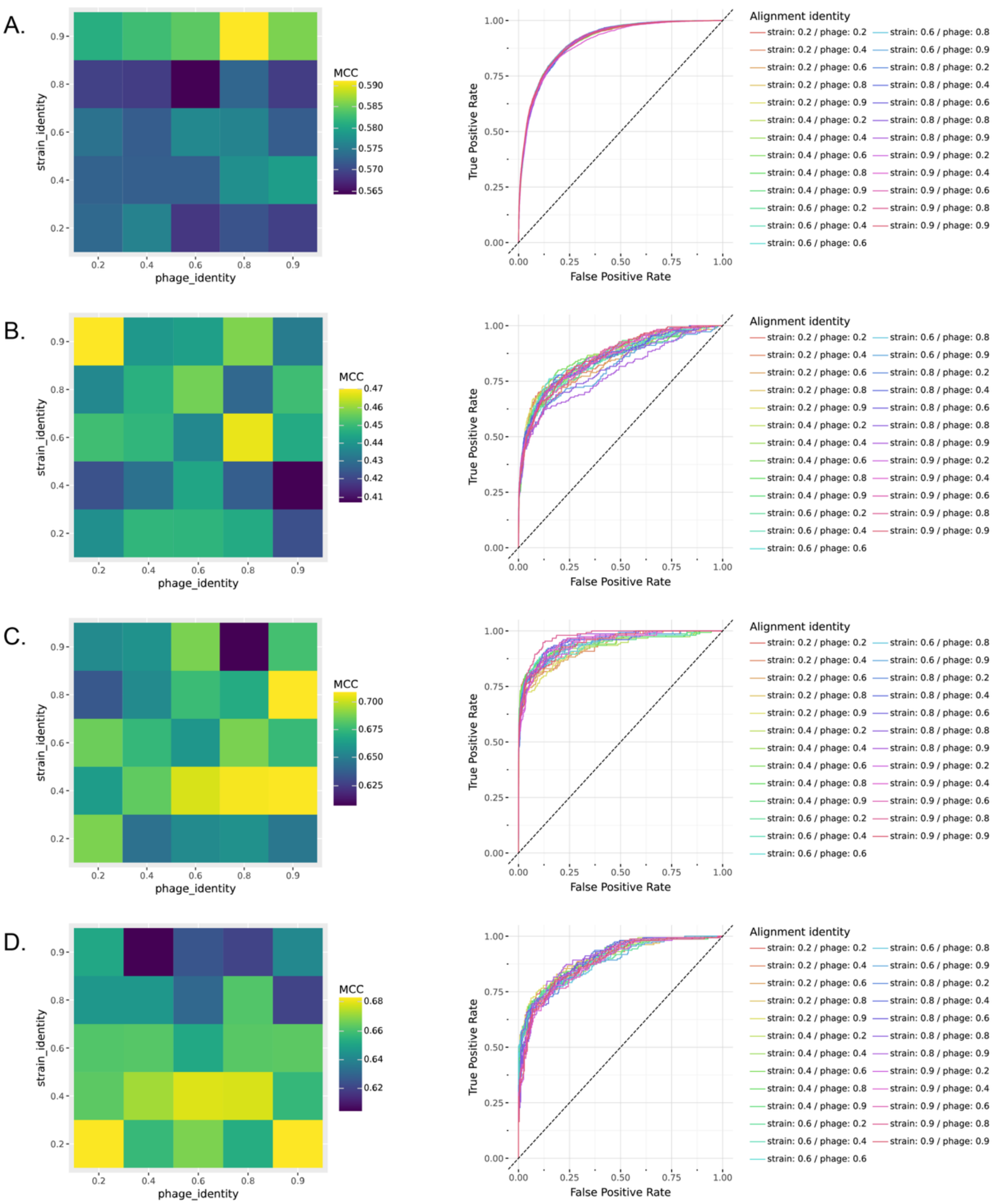

*Supplementary Figure 9. Impact of MMSeqs2 identity thresholds on model performance.* Heatmaps show model performance (MCC) across 25 MMSeqs2 identity threshold combinations. Receiver operating characteristic curves should model performance (AUROC) across 25 MMSeqs2 identity threshold combinations across *E. coli* (A), *Klebsiella*-1 (B), *Klebsiella*-2 (C), and *Pseudomonas* (D) datasets

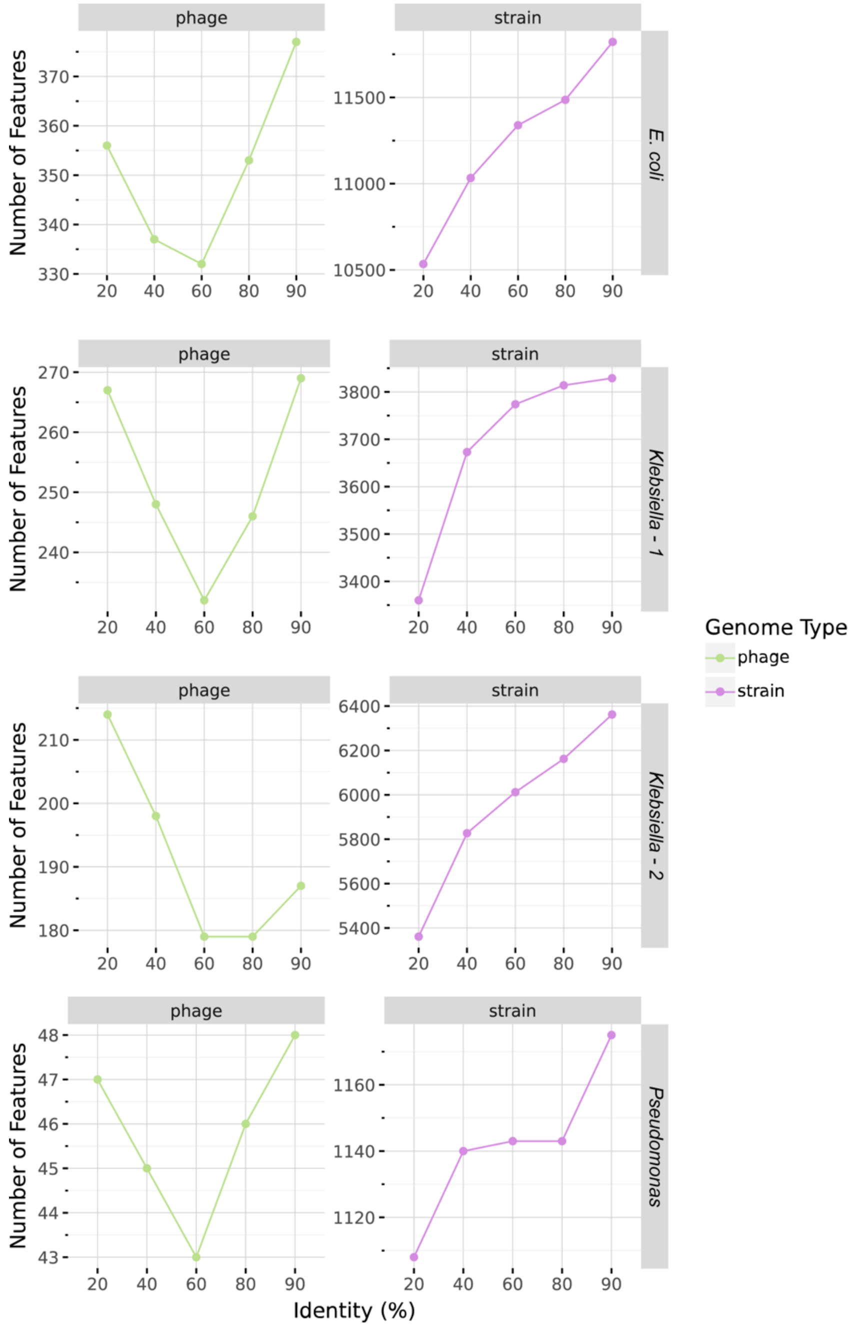

*Supplementary Figure 10. Feature counts for MMSeqs2 identity thresholds.* Lines show the impact of MMSeqs2 identity threshold on the total number of phage (green) and bacterial strain (purple) protein family features across development datasets.

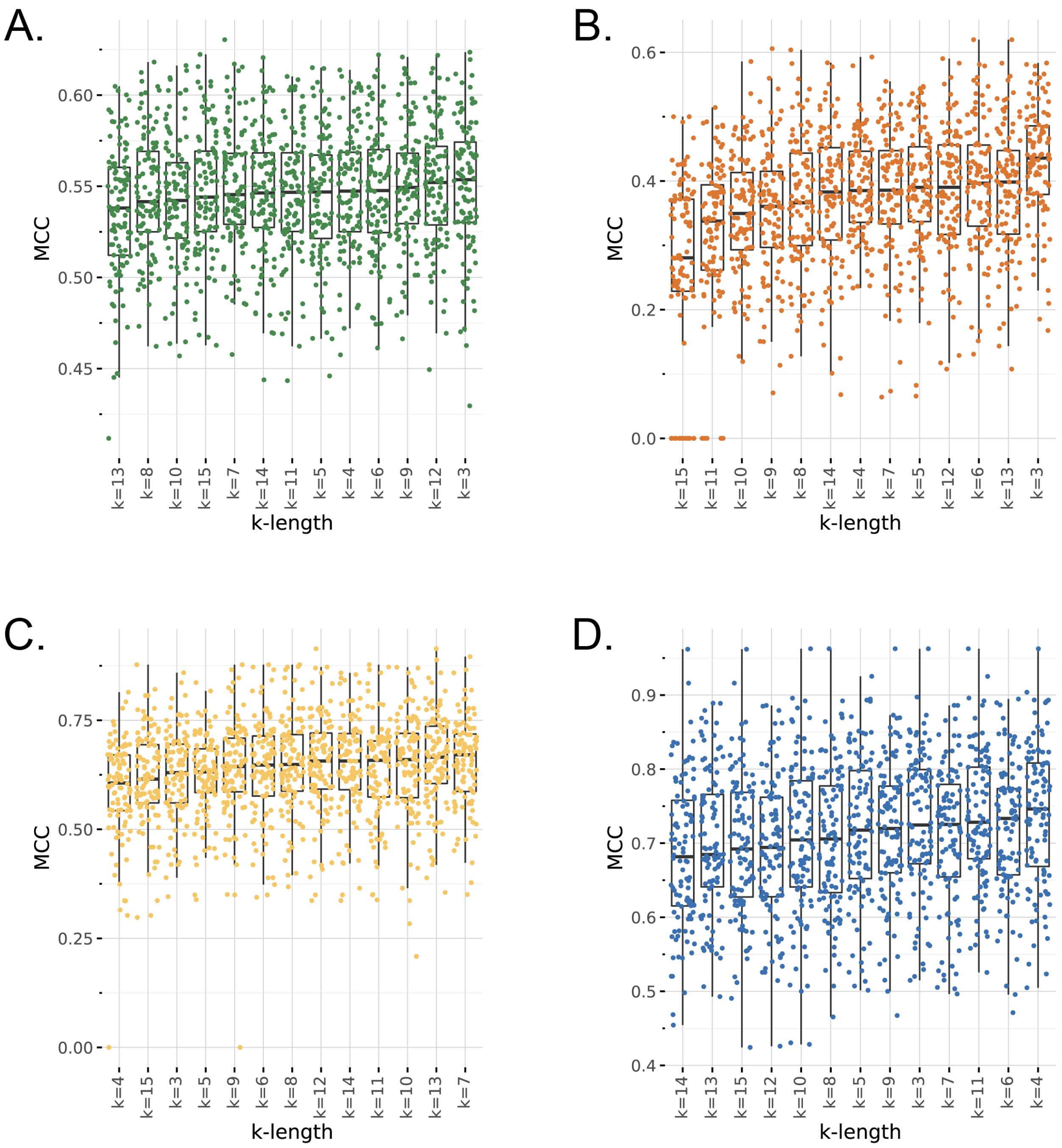

*Supplementary Figure 11. The impact of k-length in E. coli (A), Klebsiella-1 (B), Klebsiella-2 (C), and Pseudomonas (D) datasets.*

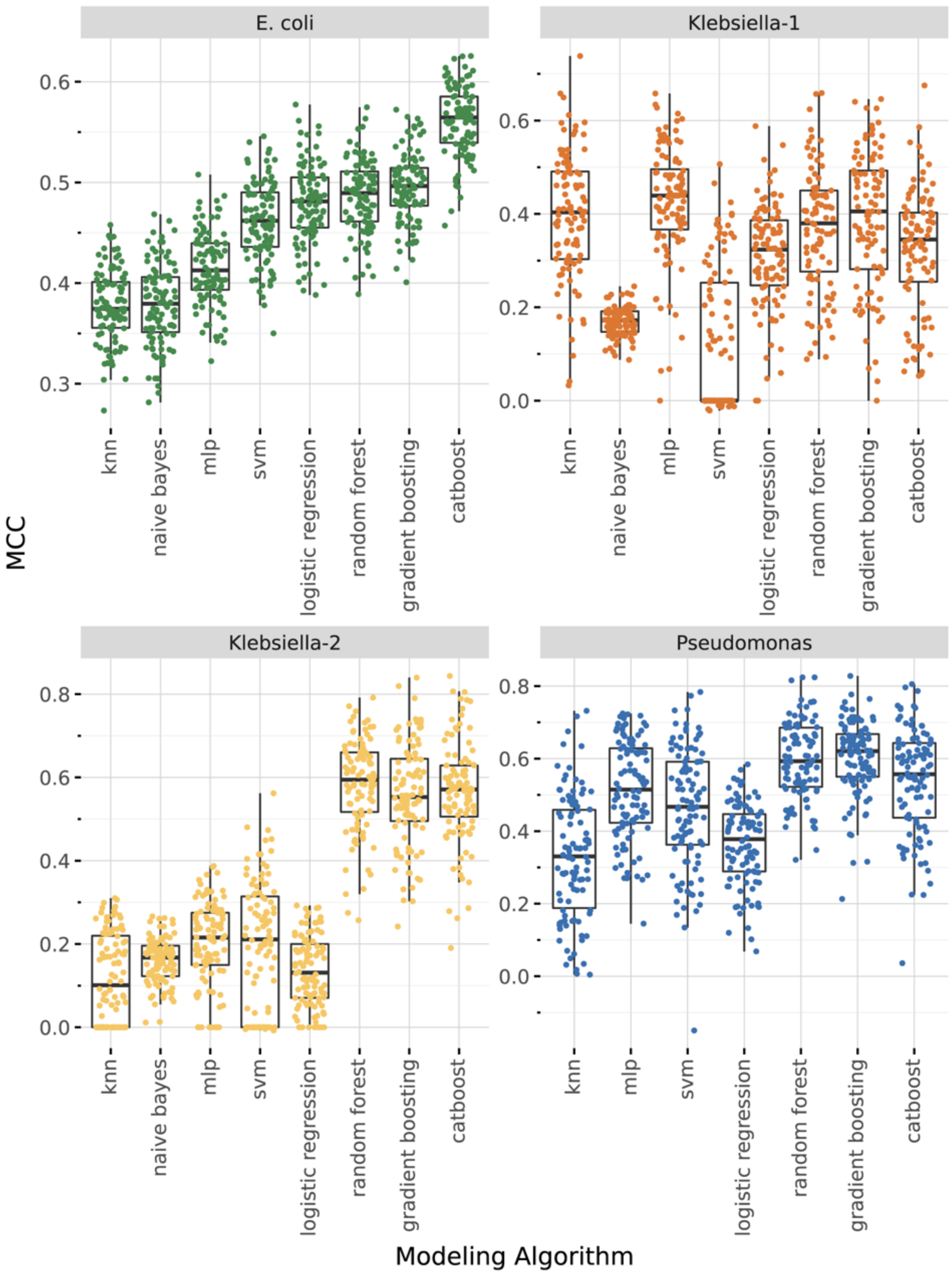

*Supplementary Figure 12. Modeling algorithm comparison.* Boxplots show performance (MCC) of 100 models used for ensemble learning approach across modeling algorithms and development datasets.

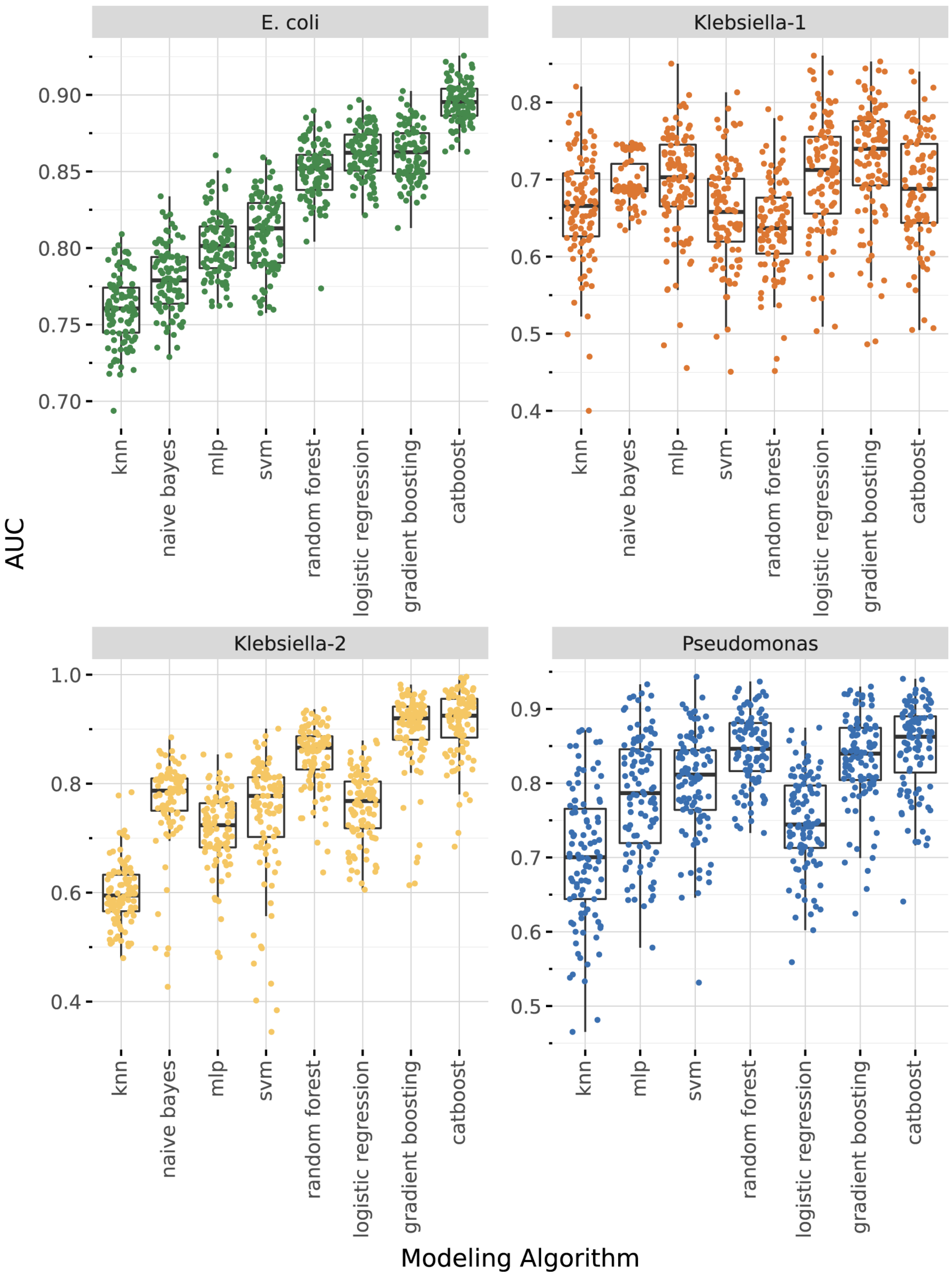

693

694 *Supplementary Figure 13. Modeling algorithm comparison.* Boxplots show performance (AUROC) of 100 models used for ensemble learning approach across modeling algorithms and development datasets.

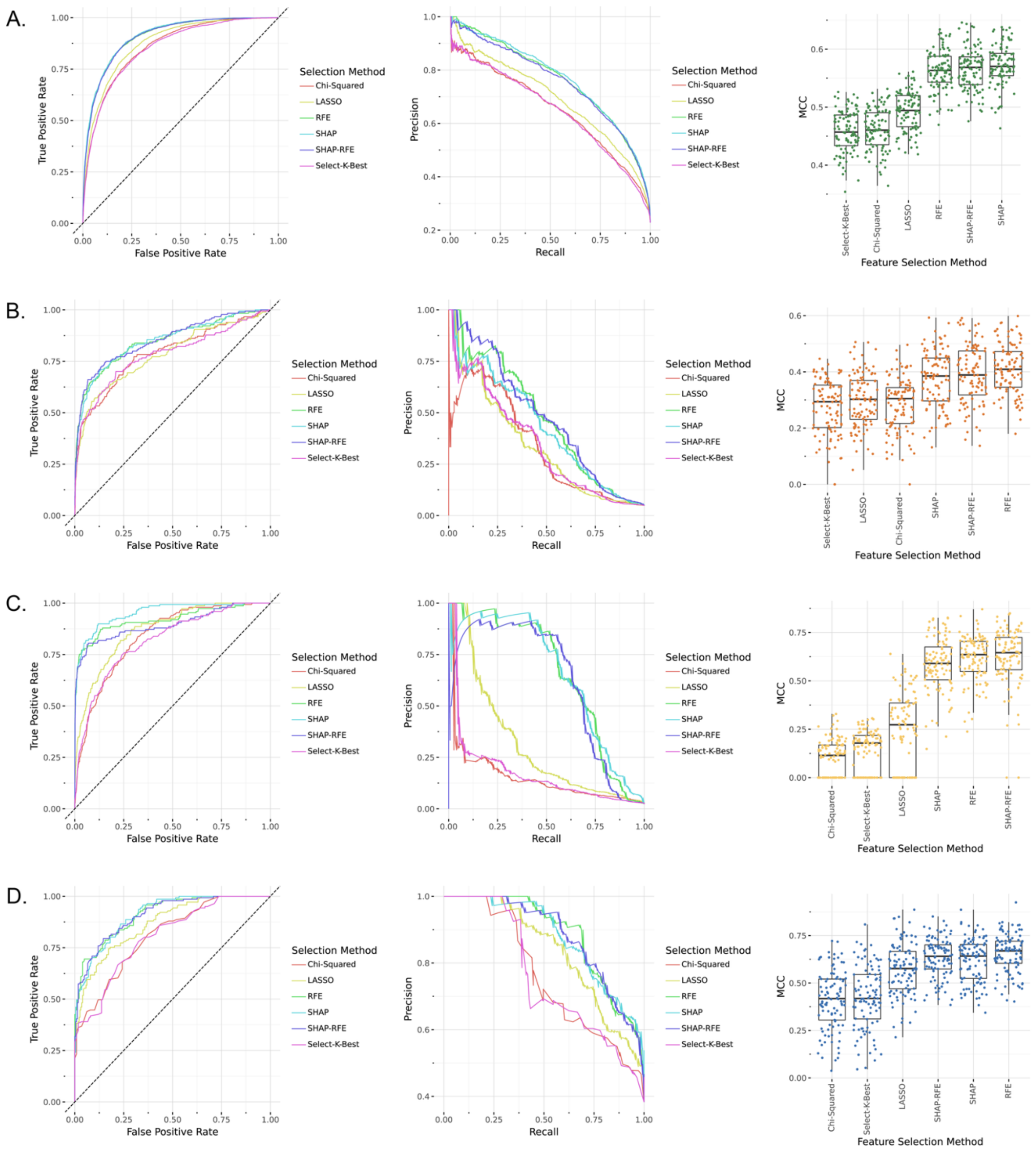

*Supplementary Figure 14. Impact of feature selection algorithm on model performance.* ROC curves, precision-recall curves and the distribution of model performances comparing 6 feature selection algorithms across *E. coli* (A), *Klebsiella*-1 (B), *Klebsiella*-2 (C), and *Pseudomonas* (D) datasets.

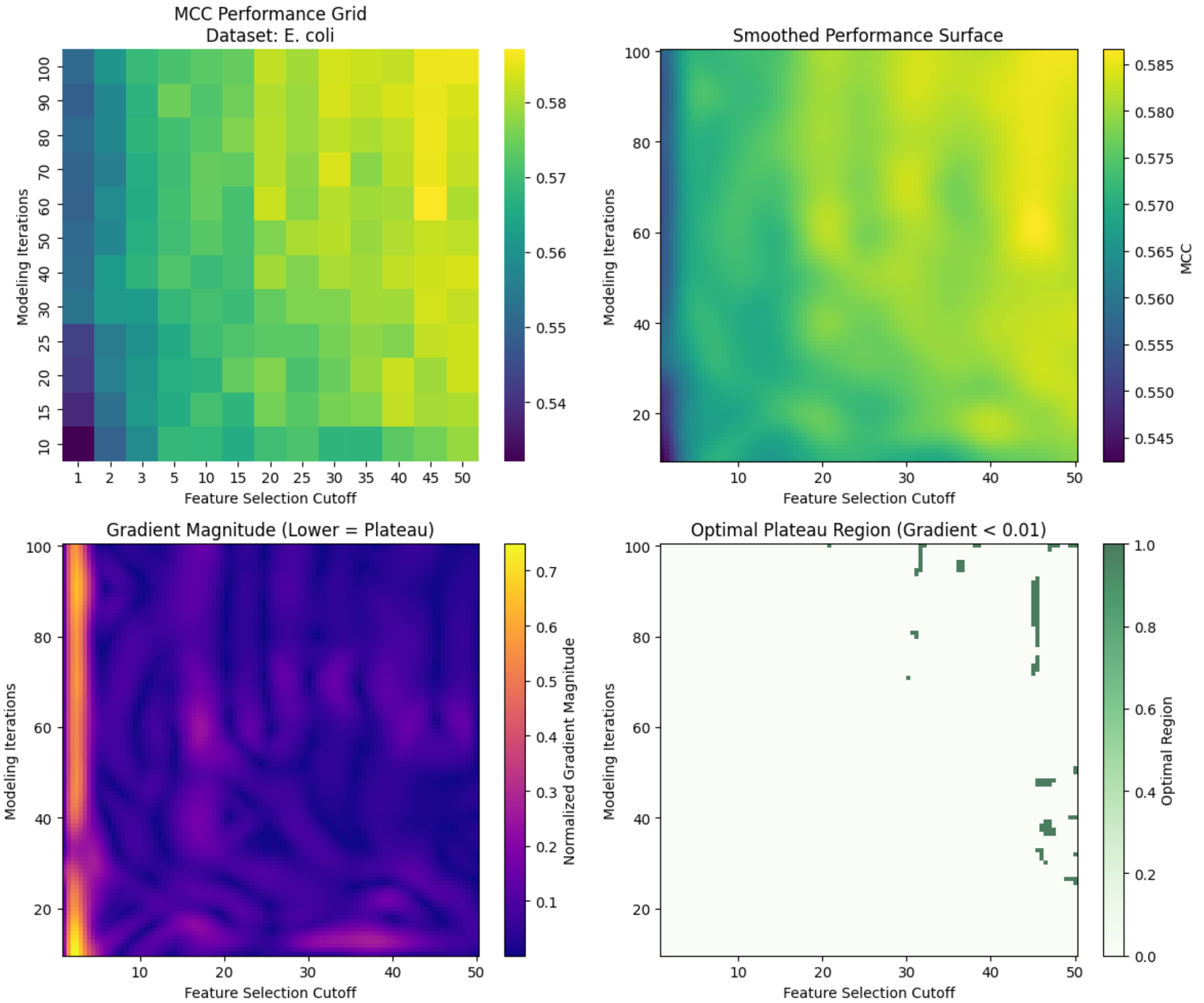

*Supplementary Figure 15. Impact of ensemble features selection and modeling approach iterations in E. coli dataset.*

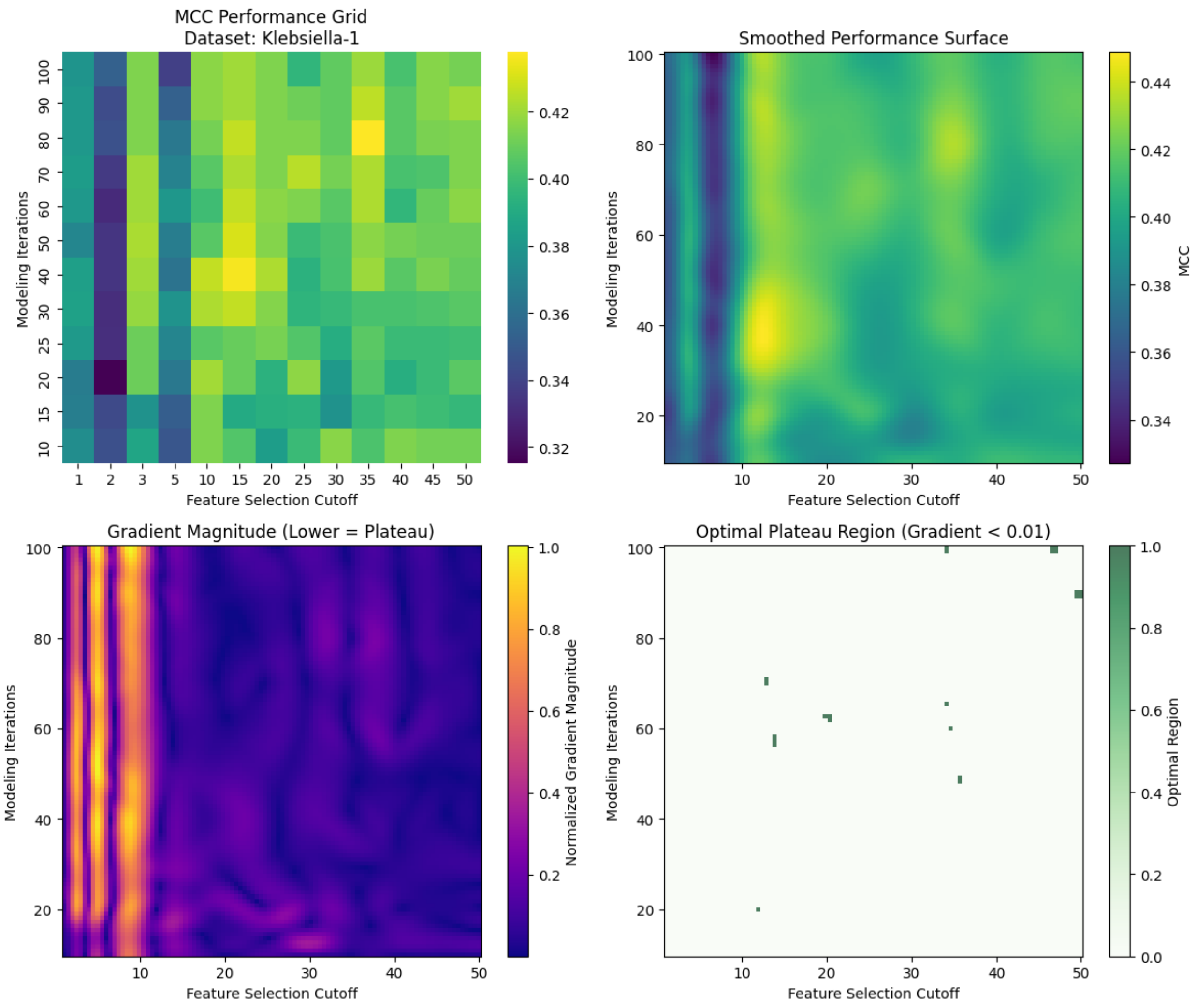

*Supplementary Figure 16. Impact of ensemble features selection and modeling approach iterations in Klebsiella-1 dataset.*

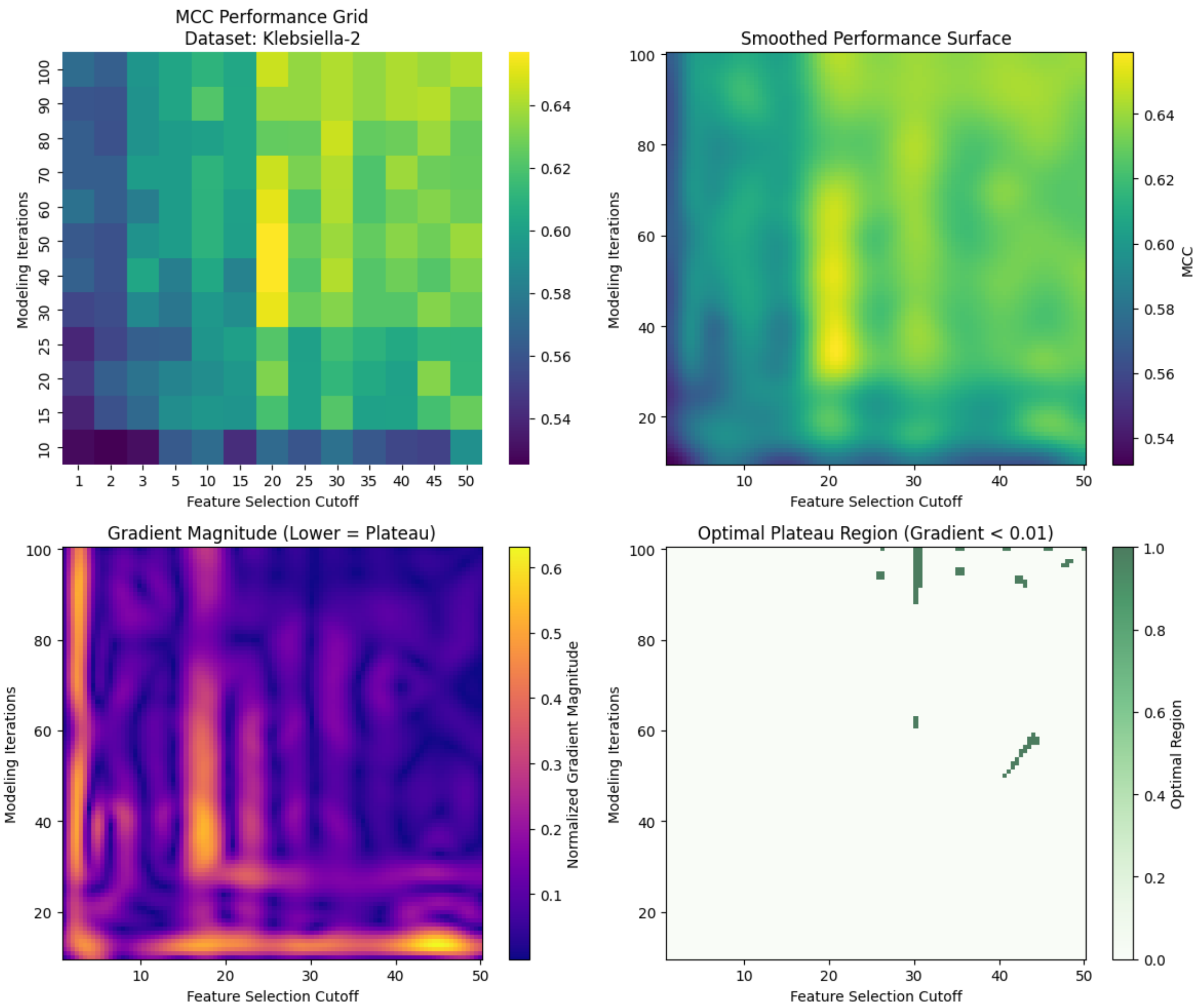

*Supplementary Figure 17. Impact of ensemble features selection and modeling approach iterations in Klebsiella-2 dataset.*

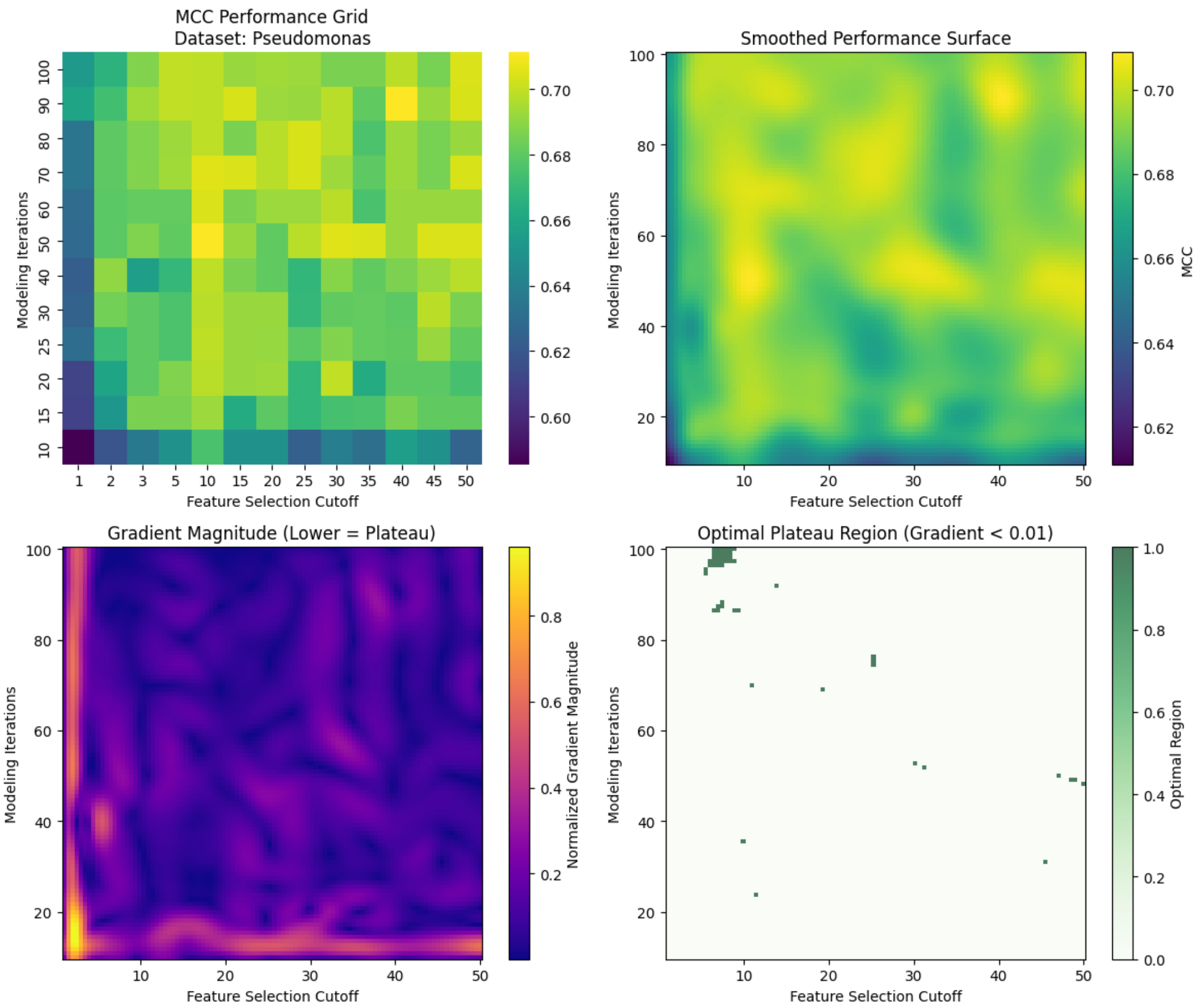

*Supplementary Figure 18. Impact of ensemble features selection and modeling approach iterations in Pseudomonas dataset.*

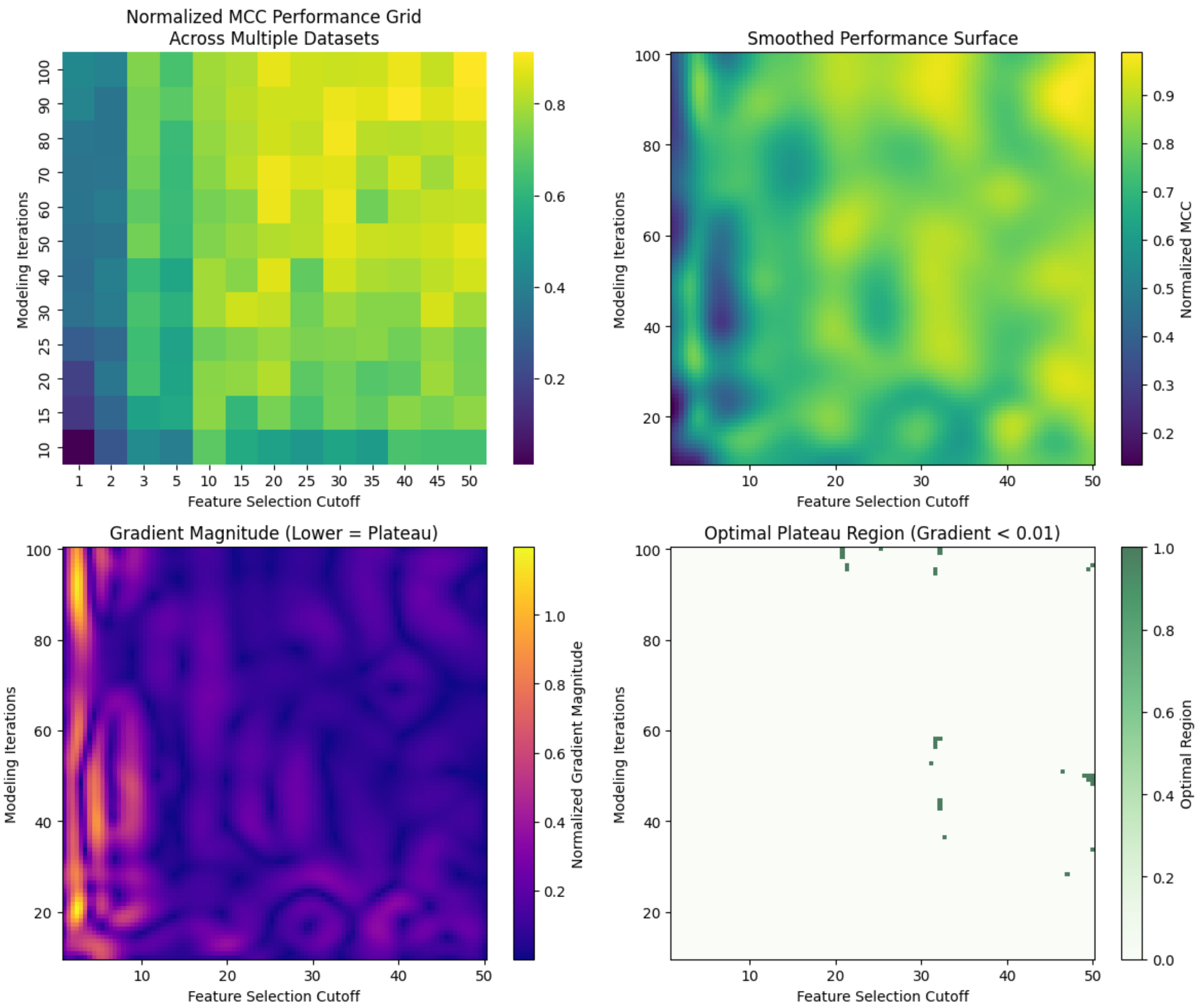

*Supplementary Figure 19. Combined analysis of the impact of ensemble features selection and modeling approach iterations in all development datasets.*

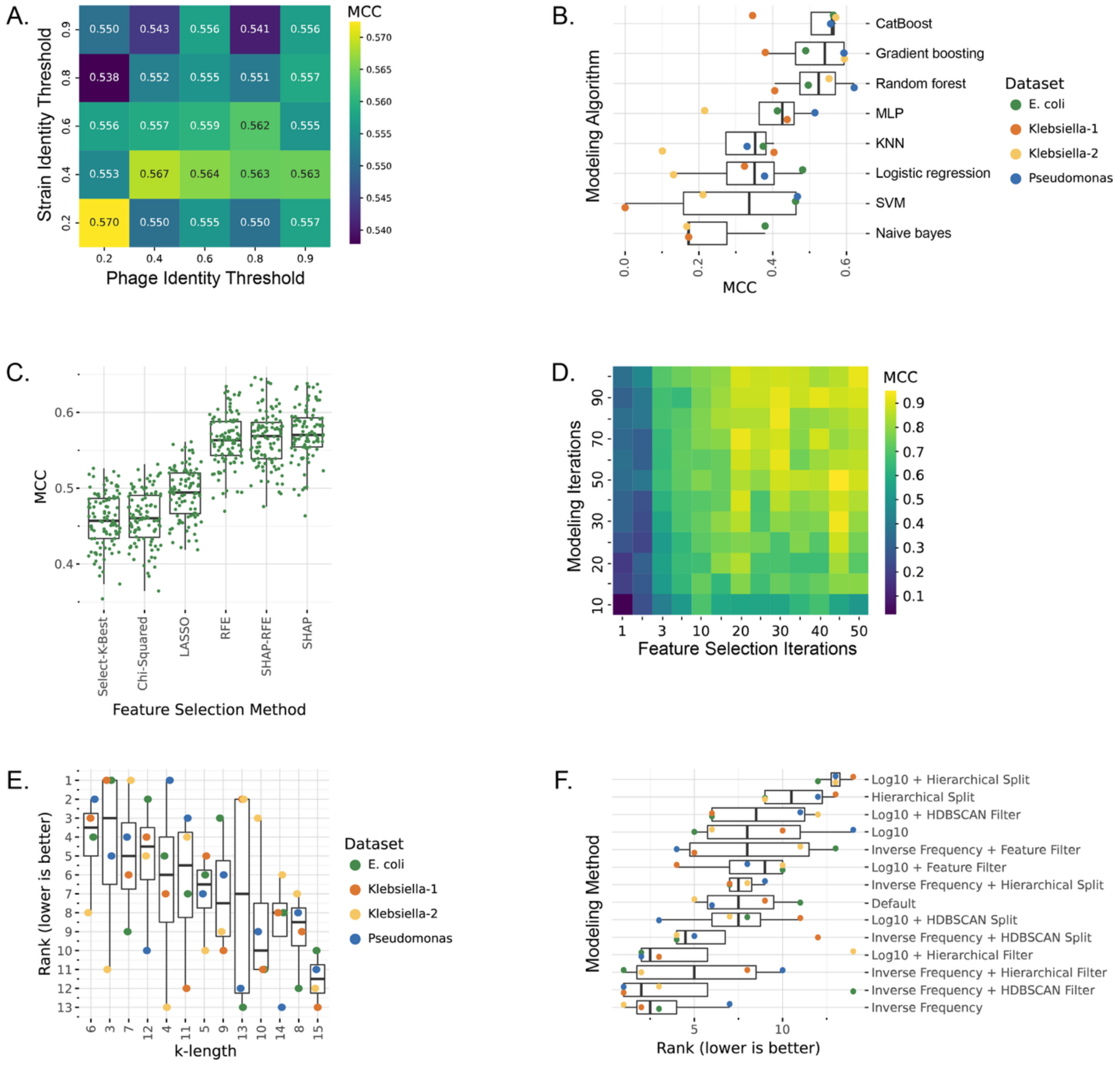

*Supplementary Figure 20. Modeling workflow optimization*. A) Mixed effect model of MMSeqs2 identity thresholds on model performance (MCC). B) Impact of modeling algorithm on model performance across datasets (MCC). C) Impact of feature selection method on training model performance (MCC) in the *E. coli* dataset. D) Impact of feature selection and modeling iterations on model performance (MCC) in the *E. coli* dataset. E) Rank order impact of *k* length on model performance (MCC) of unseen bacterial strains across datasets, based on 20-fold cross-validation F) Rank order impact of hyperparameters on model performance (MCC) of unseen strains across datasets, based on 20-fold cross-validation.

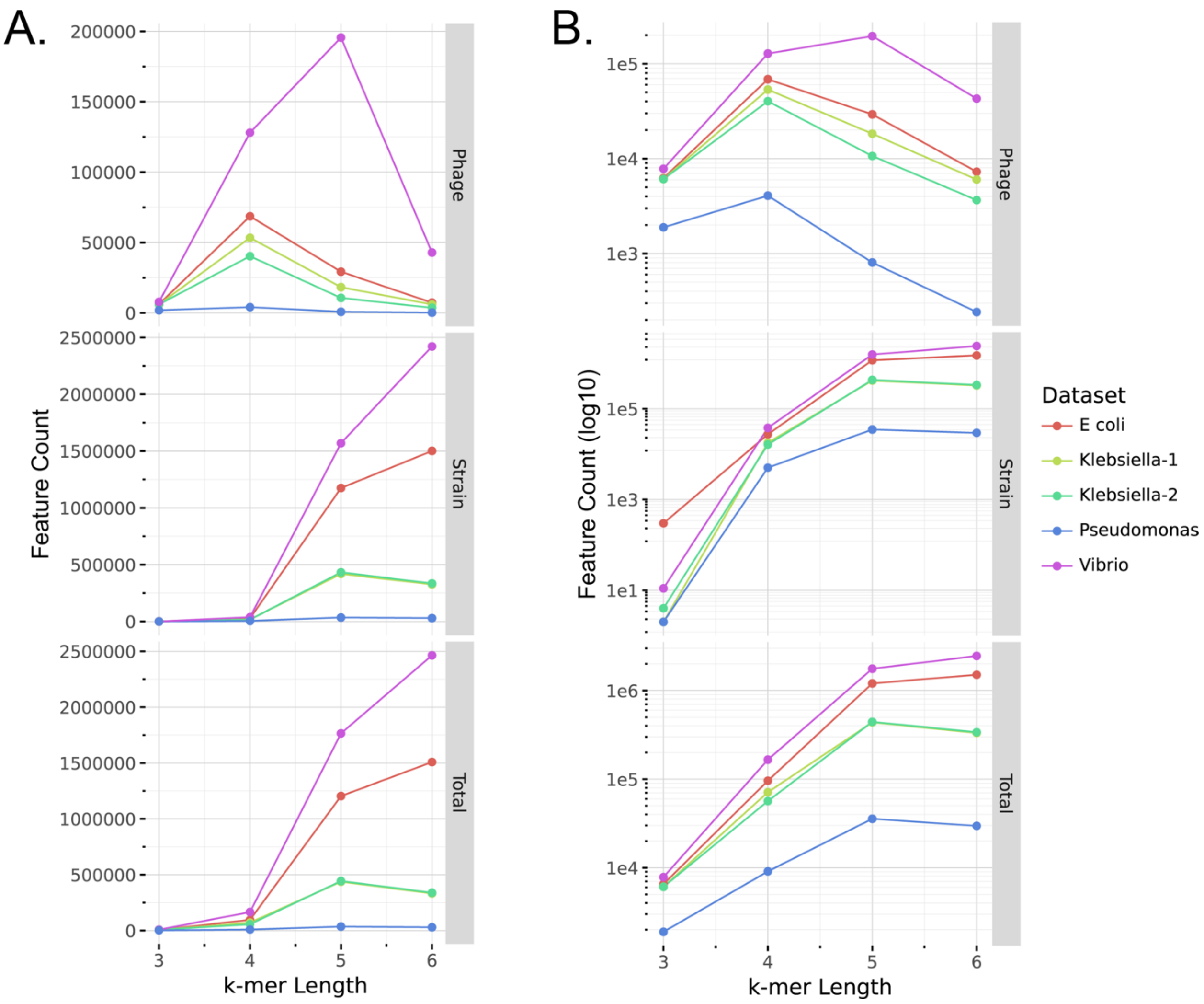

*Supplementary Figure 21. K*-mer feature counts by *k* values. Counts of *k*-mer-based features across *k* values from 3-6 for phage genomes, bacterial genomes, and combined (Total) feature counts. Colors indicate dataset, with Y-axis in linear scale (A) and log scale (B).

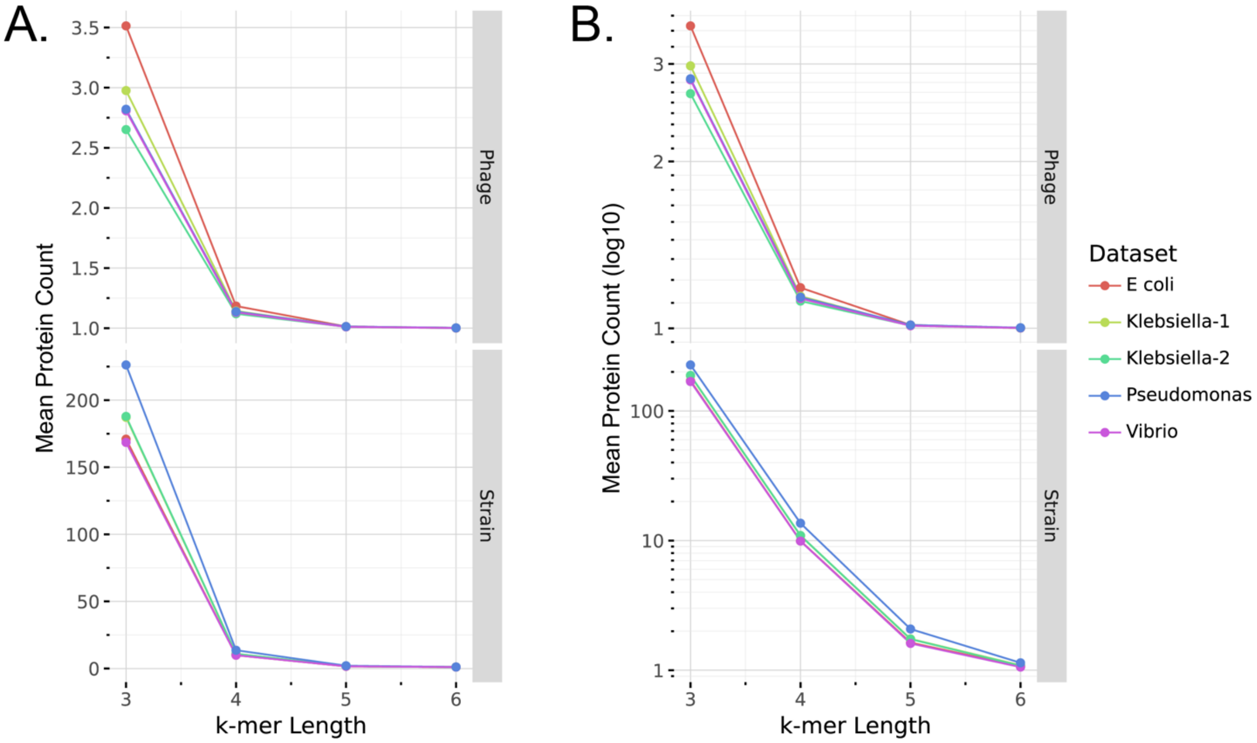

*Supplementary Figure 22.* Mean protein counts per feature per genome by *k* value. The mean protein counts represented by each *k*-mer-based features was calculated across *k* values from 3-6 for phage and bacterial genomes. Colors indicate dataset, with Y-axis in linear scale (A) and log scale (B).

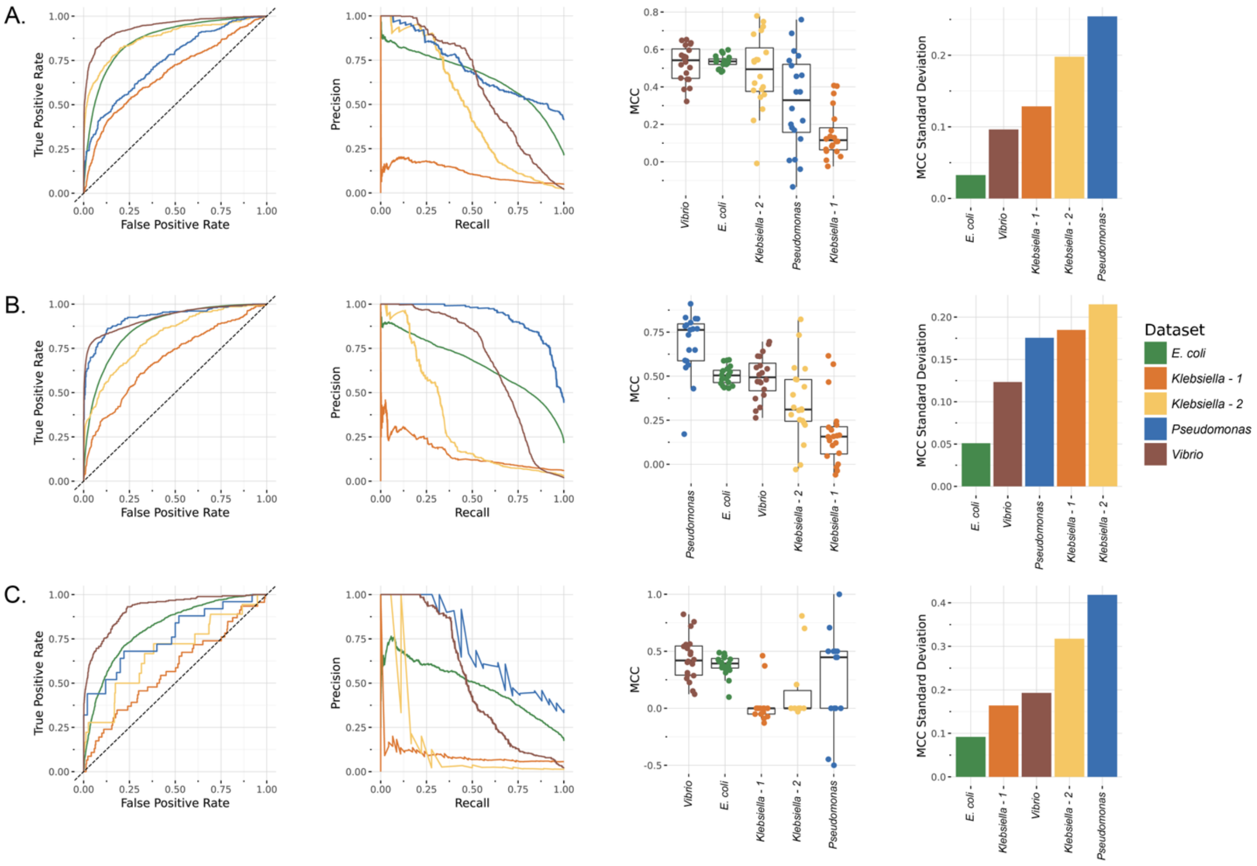

*Supplementary Figure 23. Model performance across datasets.* Receiver operating characteristic (ROC) curves show prediction performance variability across datasets, when predicting phages infecting previously unseen bacterial strains (A), hosts for previously unseen phages (B), and interactions between previously unseen bacteria-phage pairs (C).

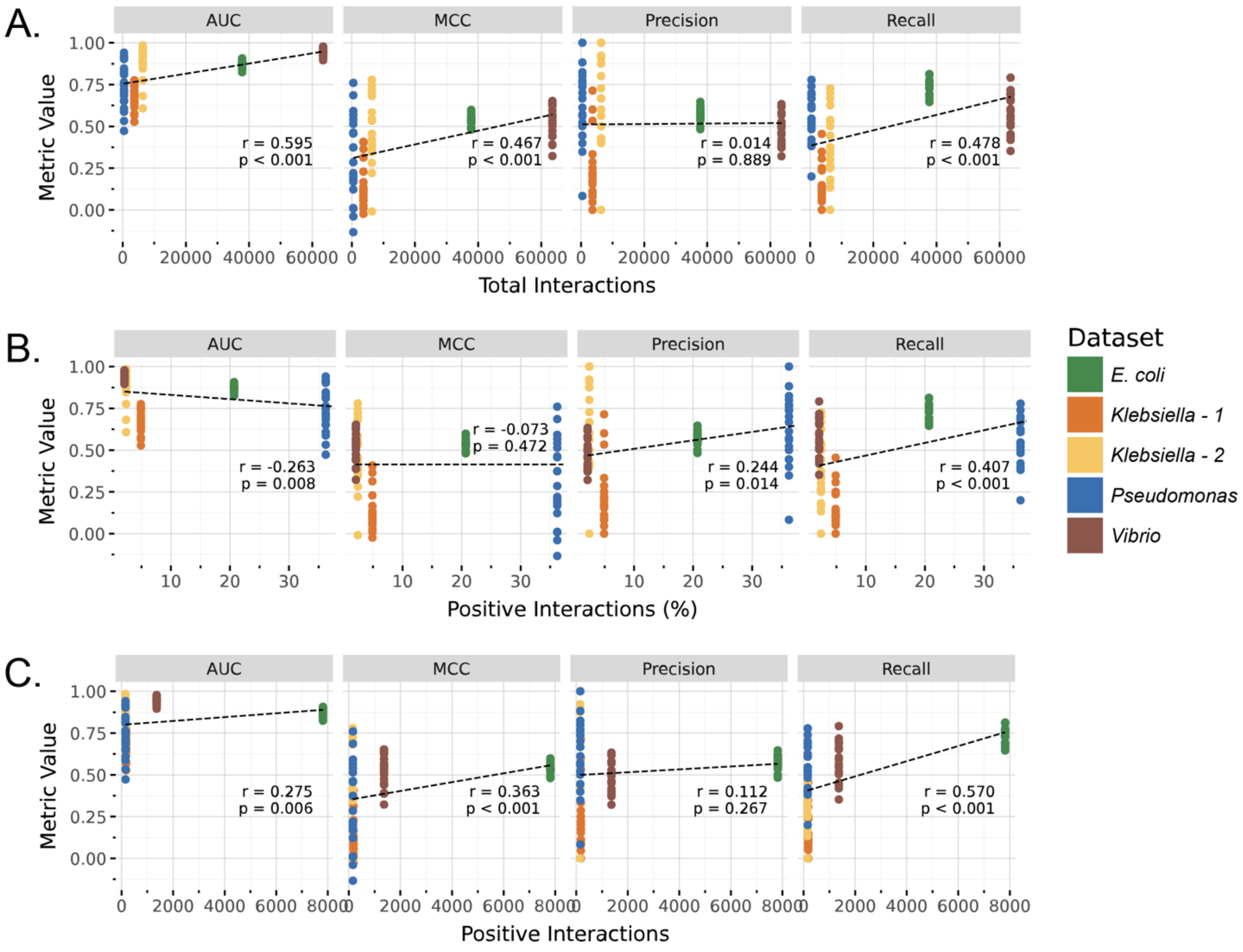

*Supplementary Figure 24. Relationship between model performance and dataset characteristics when predicting phages infecting previously unseen bacterial strains.*

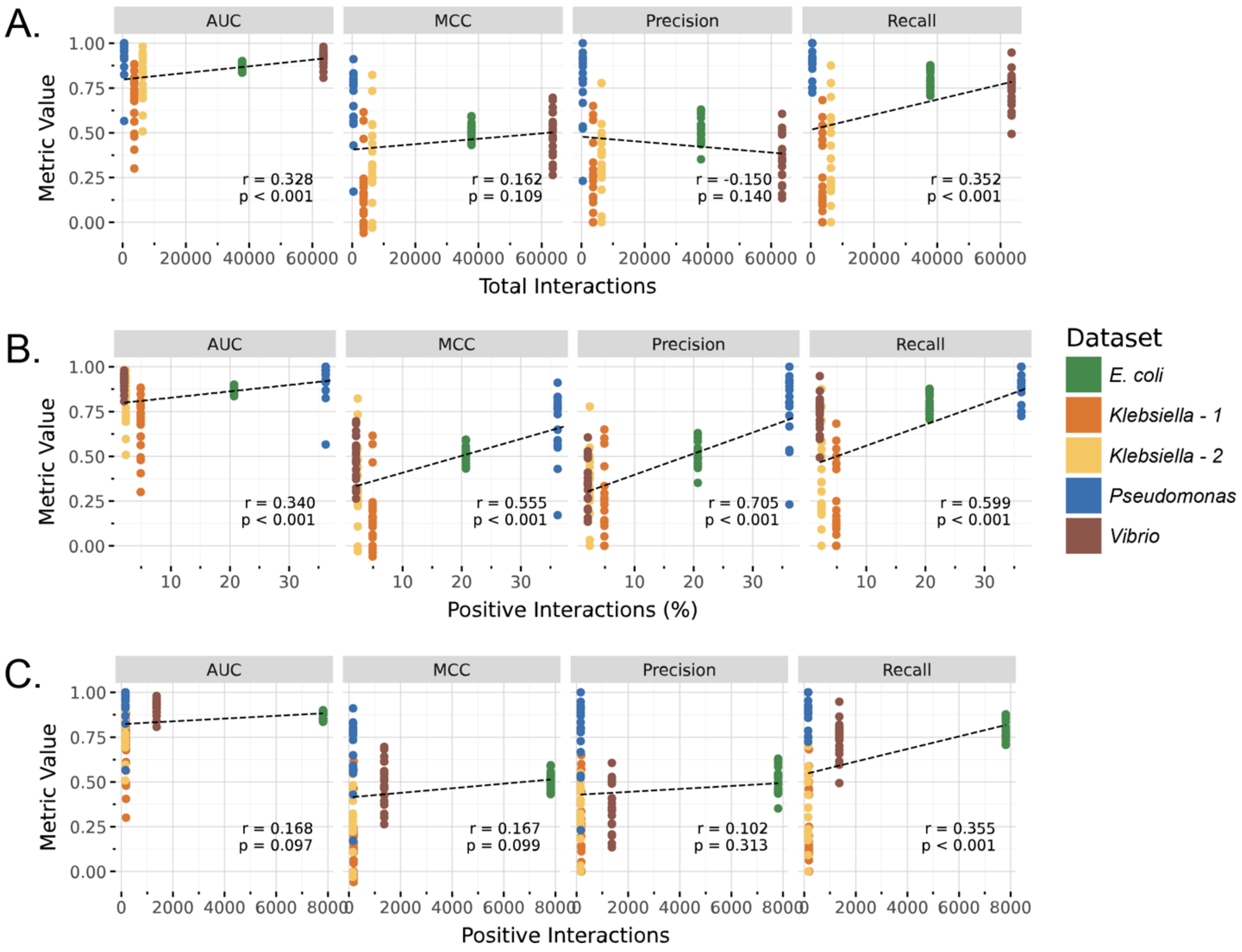

*Supplementary Figure 25. Relationship between model performance and dataset characteristics when predicting hosts for previously unseen phages.*

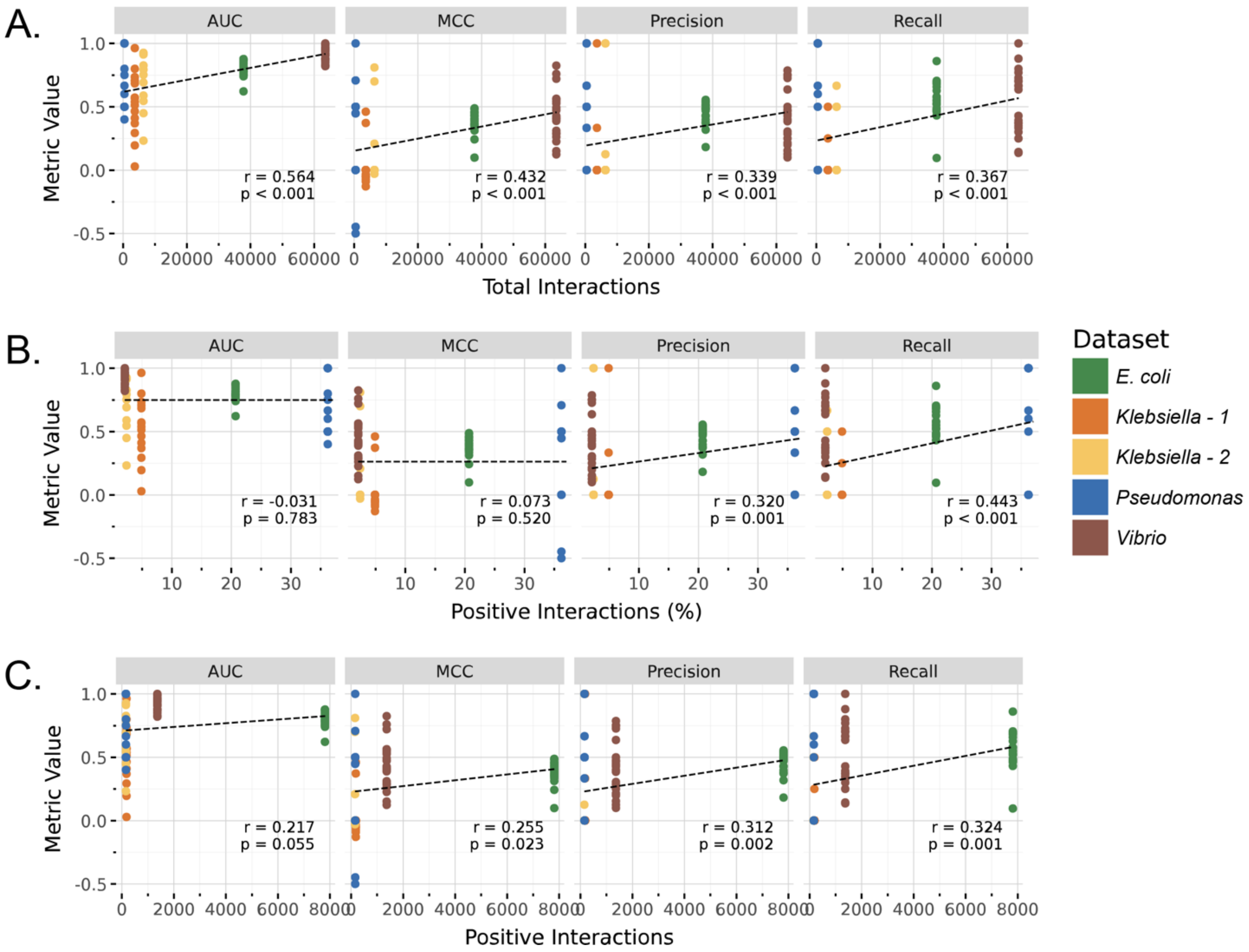

*Supplementary Figure 26. Relationship between model performance and dataset characteristics when predicting interactions between previously unseen bacteria-phage pairs.*

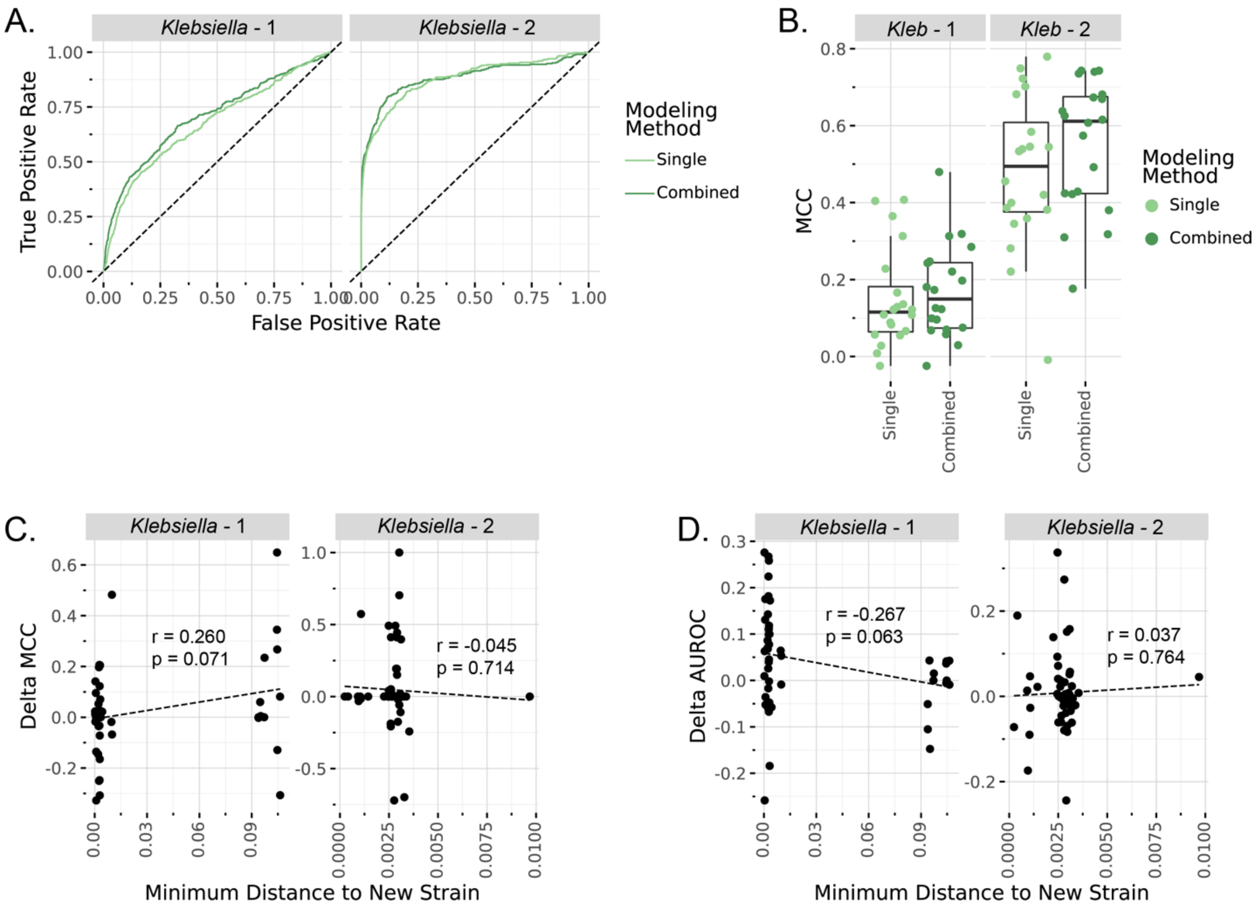

*Supplementary Figure 27. Combining datasets within the Klebsiella genus*. A) Models trained by combining both *Klebsiella* datasets (dark green) showed increased performance across both datasets, in comparison to models trained on single datasets (light green). Displayed AUROC curves are from 20-fold cross-validation. B) Boxplots show performance of individual rounds of cross validation when combining both *Klebsiella* datasets (dark green) in comparison to models trained on single datasets (light green). A significant increase in performance was observed in the *Klebsiella*-2 dataset (p = 0.024). C) The change in MCC of individual *Klebsiella* strains was compared to the phylogenetic distance to the nearest strain from the added dataset. D) The change in AUROC of individual *Klebsiella* strains was compared to the phylogenetic distance to the nearest strain from the added dataset.

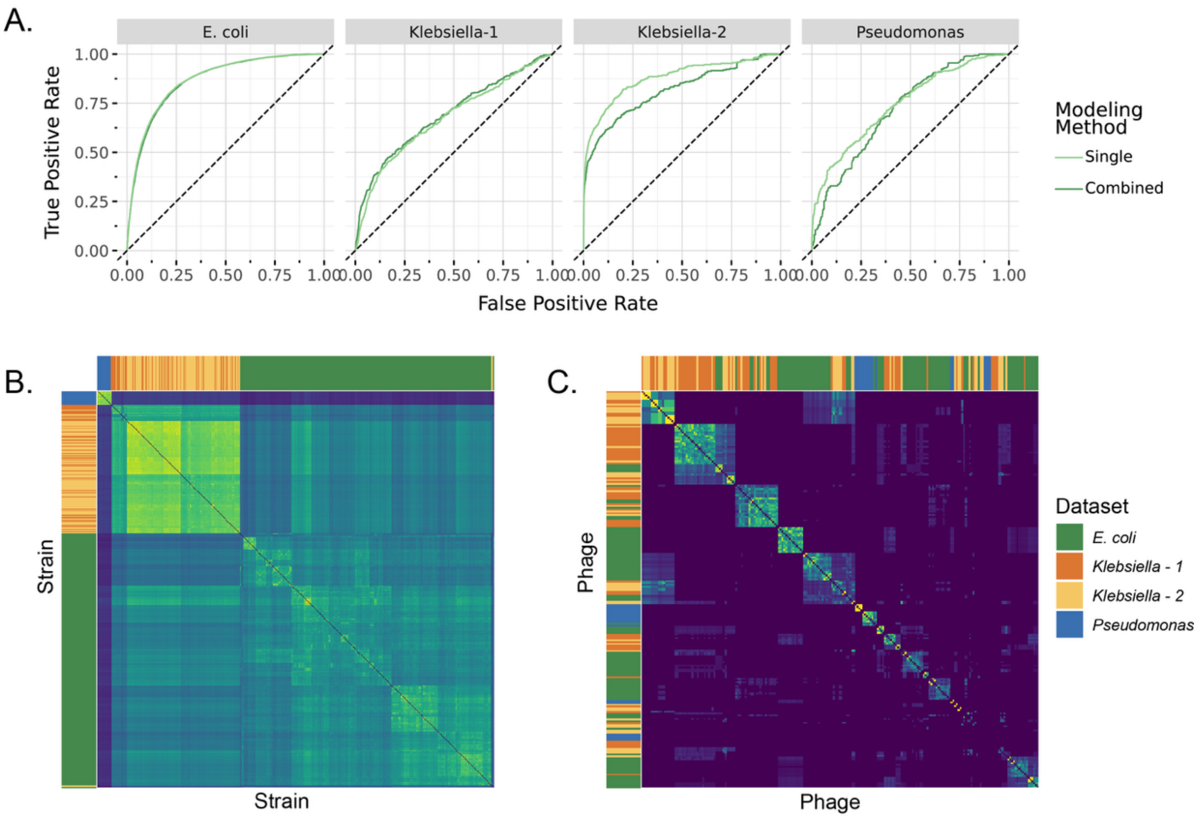

*Supplementary Figure 28. Combining datasets across genera*. A) Models trained by combining all datasets (dark green) showed increased performance in the *Klebsiella* datasets, and no change or a slightly decrease in performance in *E. coli* and *Pseudomonas* datasets, in comparison to models trained on single datasets (light green). Displayed results are from 20-fold cross-validation. B) Jaccard distances between bacterial strains, based on predictive feature content, shows show distinct clusters by genus. Colored bars show dataset. C) Jaccard distances between phages, based on predictive feature content, show more modularity and overlap between genera. Colored bars show phage dataset.

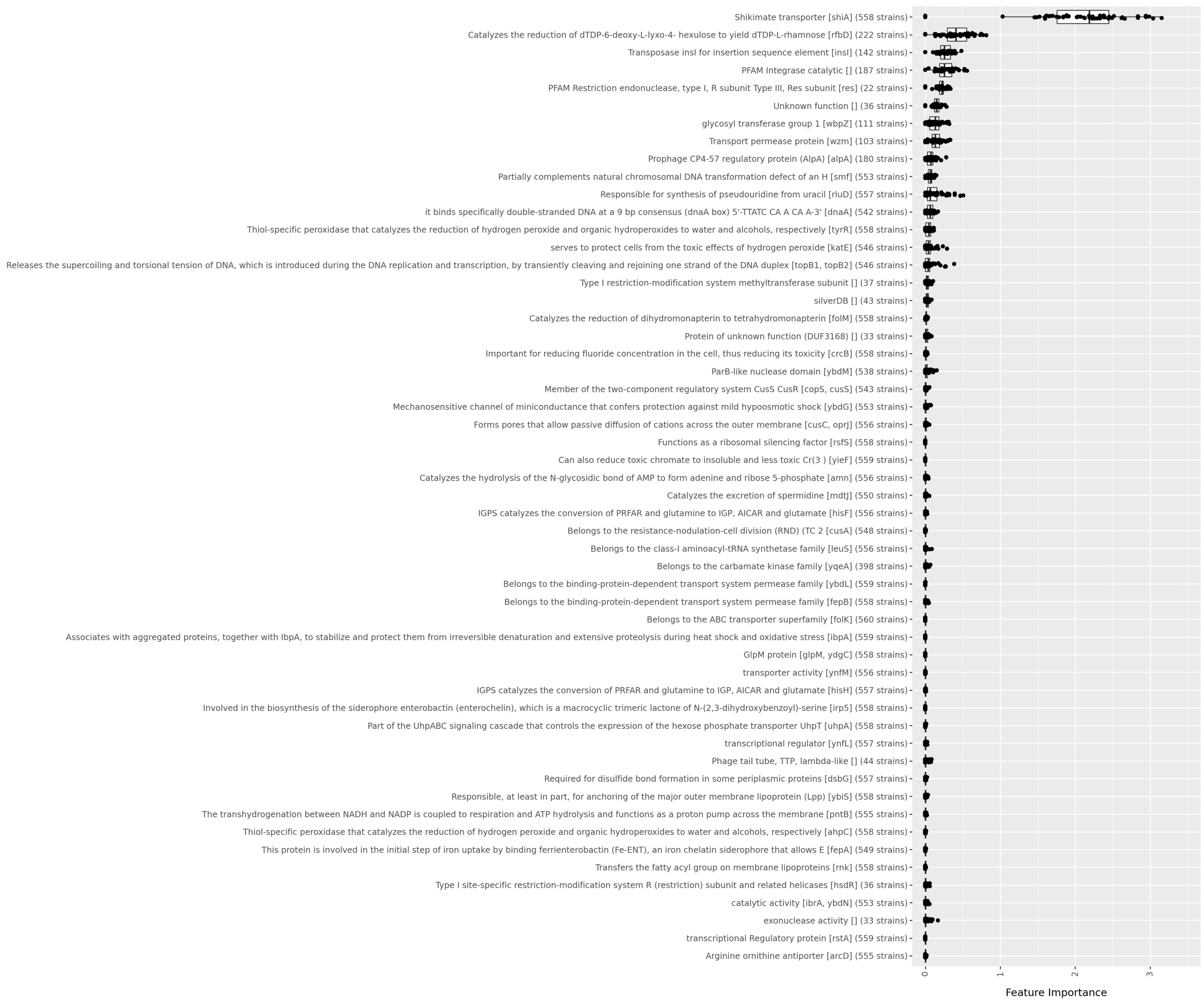

*Supplementary Figure 29. Annotations of predictive features shared across datasets.* Plot shows feature importance of 54 predictive features that were shared across *E. coli*, *Klebsiella*, and *Pseudomonas* datasets. Y-axis shows the most common annotation of genes associated with each predictive features and number of bacterial strains encoding that features. X-axis shows feature importance across 50 models used in ensemble learning.

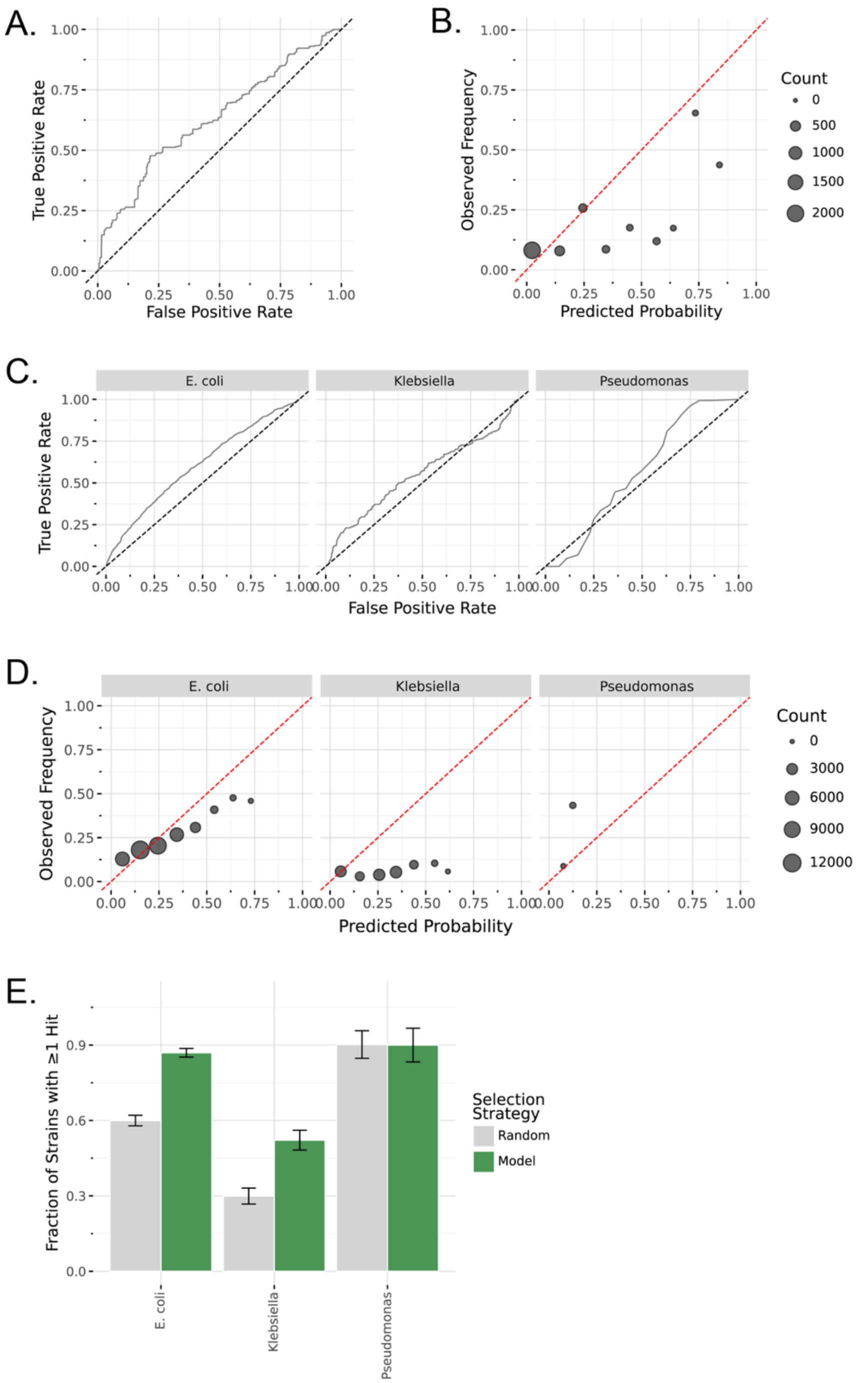

*Supplementary Figure 30. Model validation and LOGO cross-genus cross-validation.* Prediction performance on the *Klebsiella*-3 dataset using models trained on *Klebsiella*-1 and *Klebsiella*-2 datasets is shown using ROC curves (A) and calibration curves (B). Prediction performance in cross-genus cross-validation labelled with the left-out genus is shown using ROC curves (C) and calibration curves (D). Proportion of strains for which at least one interacting phage was identified in a group of 5, comparing model selection to random selection in the cross-genus cross-validation experiments.

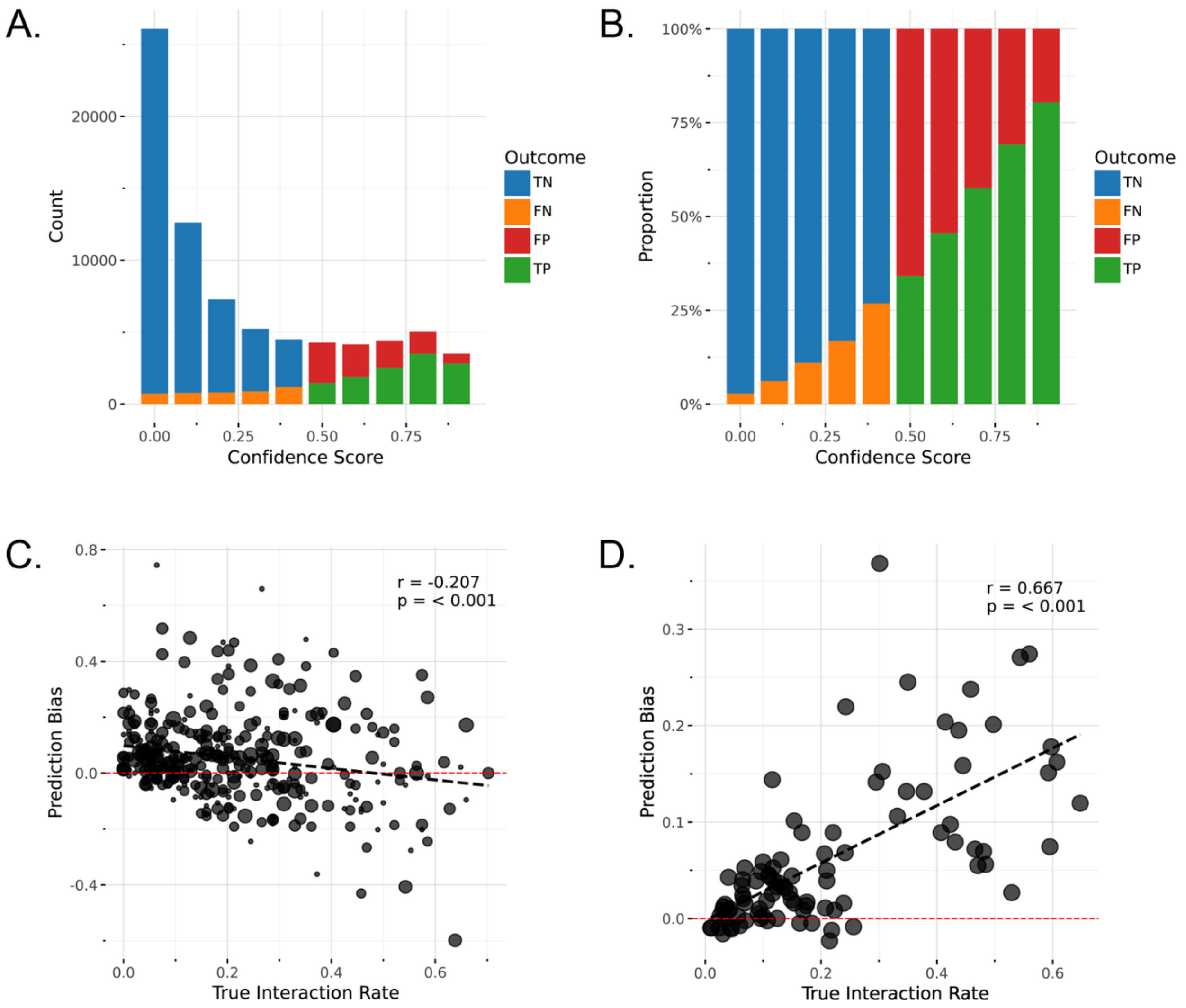

*Supplementary Figure 31. E. coli dataset prediction bias overview.* The distribution of prediction confidences across 20-fold cross-validation leaving out 10% of strains from the *E. coli* dataset. The majority of predictions were of no-infection, consistent with distributions observed in the experimental dataset (A). The proportion of true-positives (TP) and true-negatives (TN) increases as confidence scores move away from 0.5 classification threshold (B). Prediction bias compared to true infection rates show a negative correlation in strains (C) and a positive correlation in phages (D).

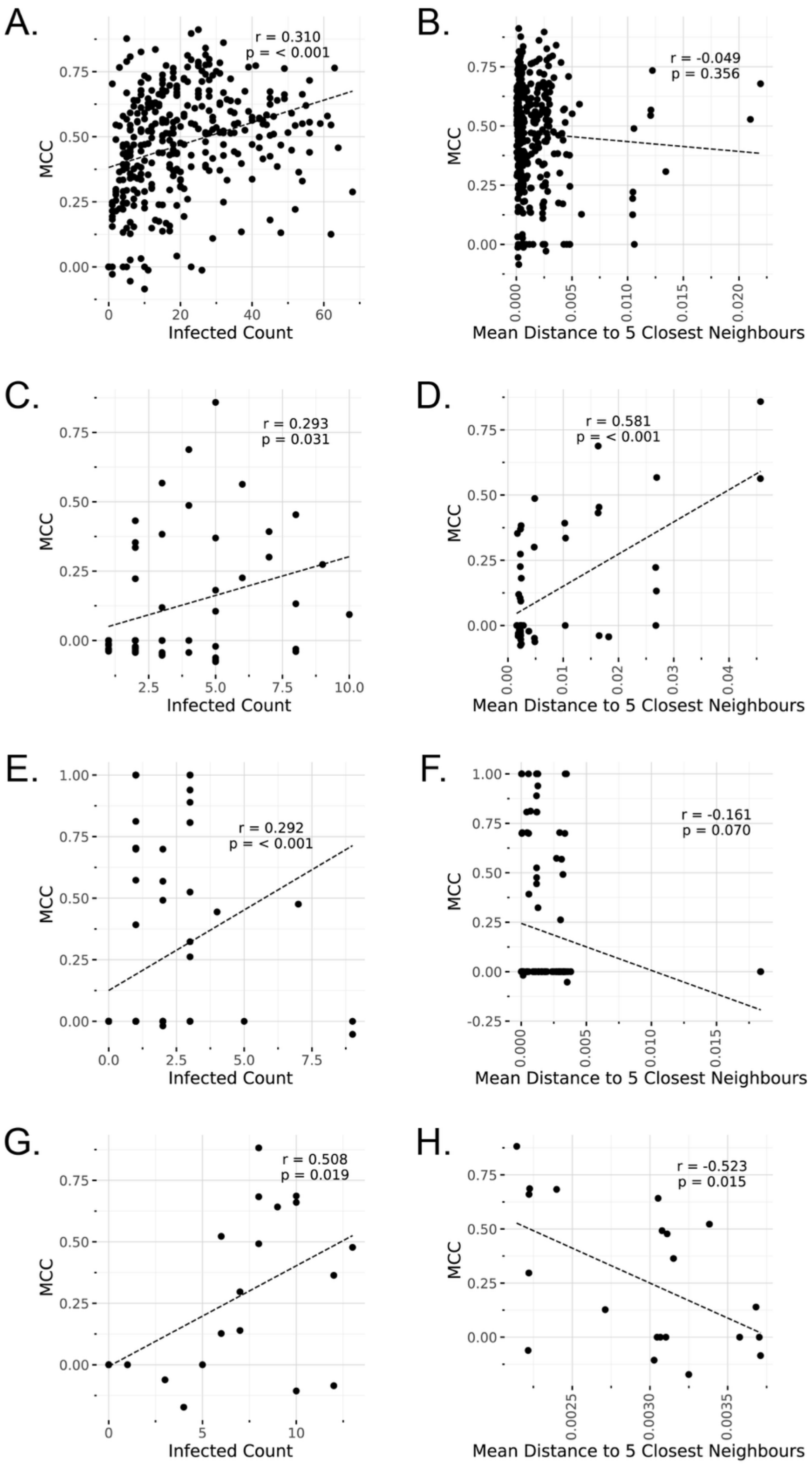

*Supplementary Figure 32. Impact of bacterial strain characteristics on prediction performance.* The relationship between strain susceptibility (number of infecting phages) and strain-level MCC (left) and phylogenetic isolation based on concatenated marker gene alignments and strain-level MCC (right) were evaluated for the *E. coli* (A / B), *Klebiella*-1 (C / D), *Klebiella*-2 (E / F), and *Pseudomonas* (G / H) datasets.

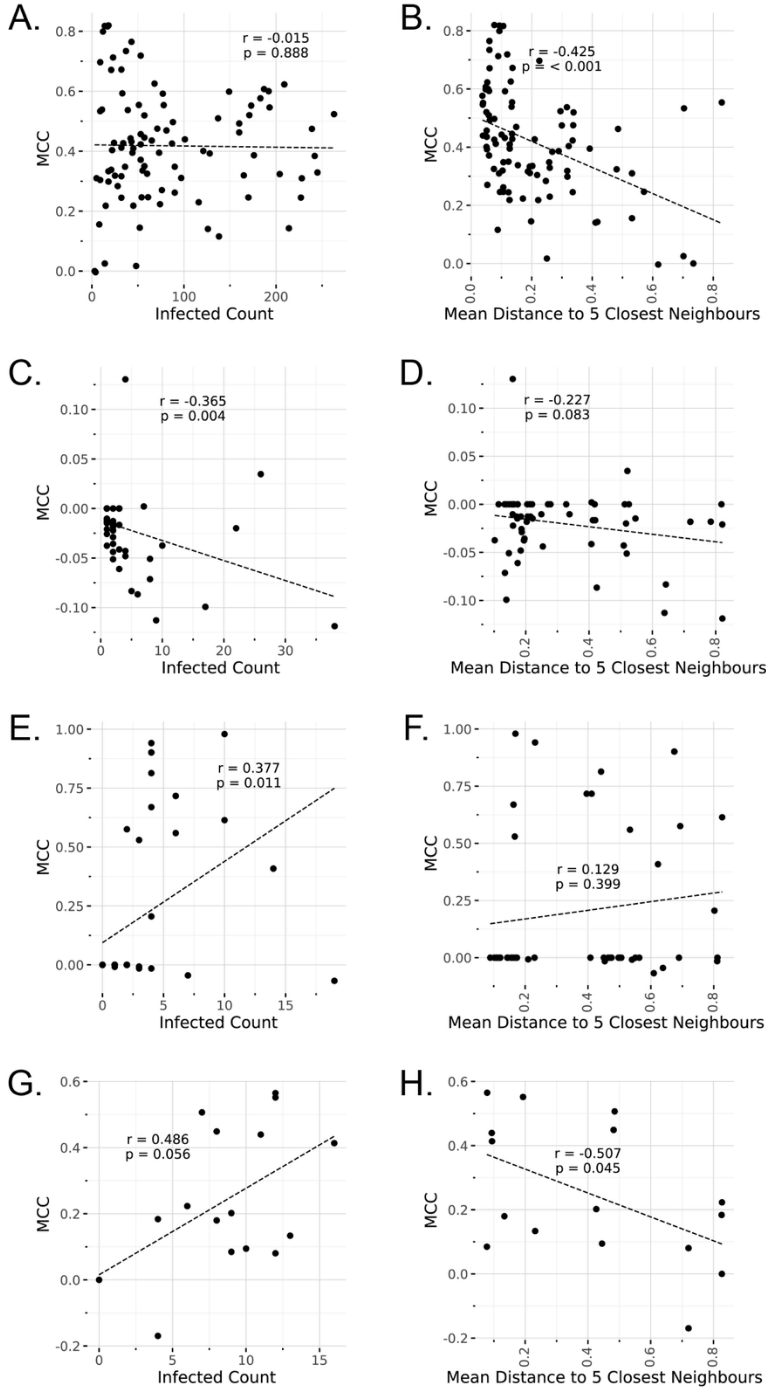

*Supplementary Figure 33. Impact of phage characteristics on prediction performance.* The relationship between phage infectivity (number of infected strains) and phage-level MCC (left) and phylogenetic isolation based on proteomic equivalence (PEG) and phage -level MCC (right) were evaluated for the *E. coli* (A / B), *Klebiella*-1 (C / D), *Klebiella*-2 (E / F), and *Pseudomonas* (G / H) datasets.

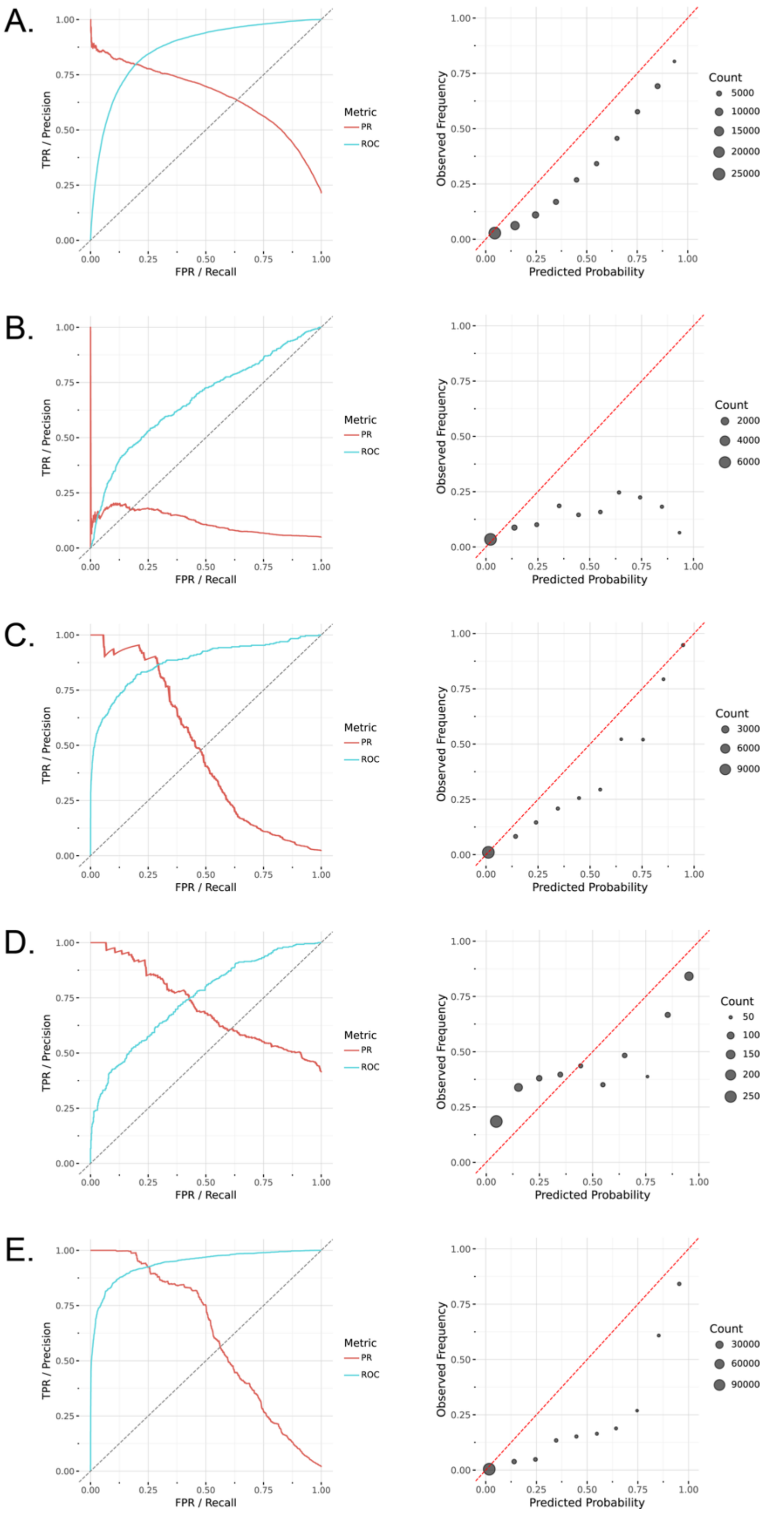

*Supplementary Figure 34. Predictive model performance and calibration curves.* Column 1 shows ROC (blue) and precision-recall (red) curves. Column 2 shows calibration curves comparing predicted vs. observed infection rates for the *E. coli* (A), *Klebiella*-1 (B), *Klebiella*-2 (C), *Pseudomonas* (D), and *Vibrionaceae* (E) datasets.

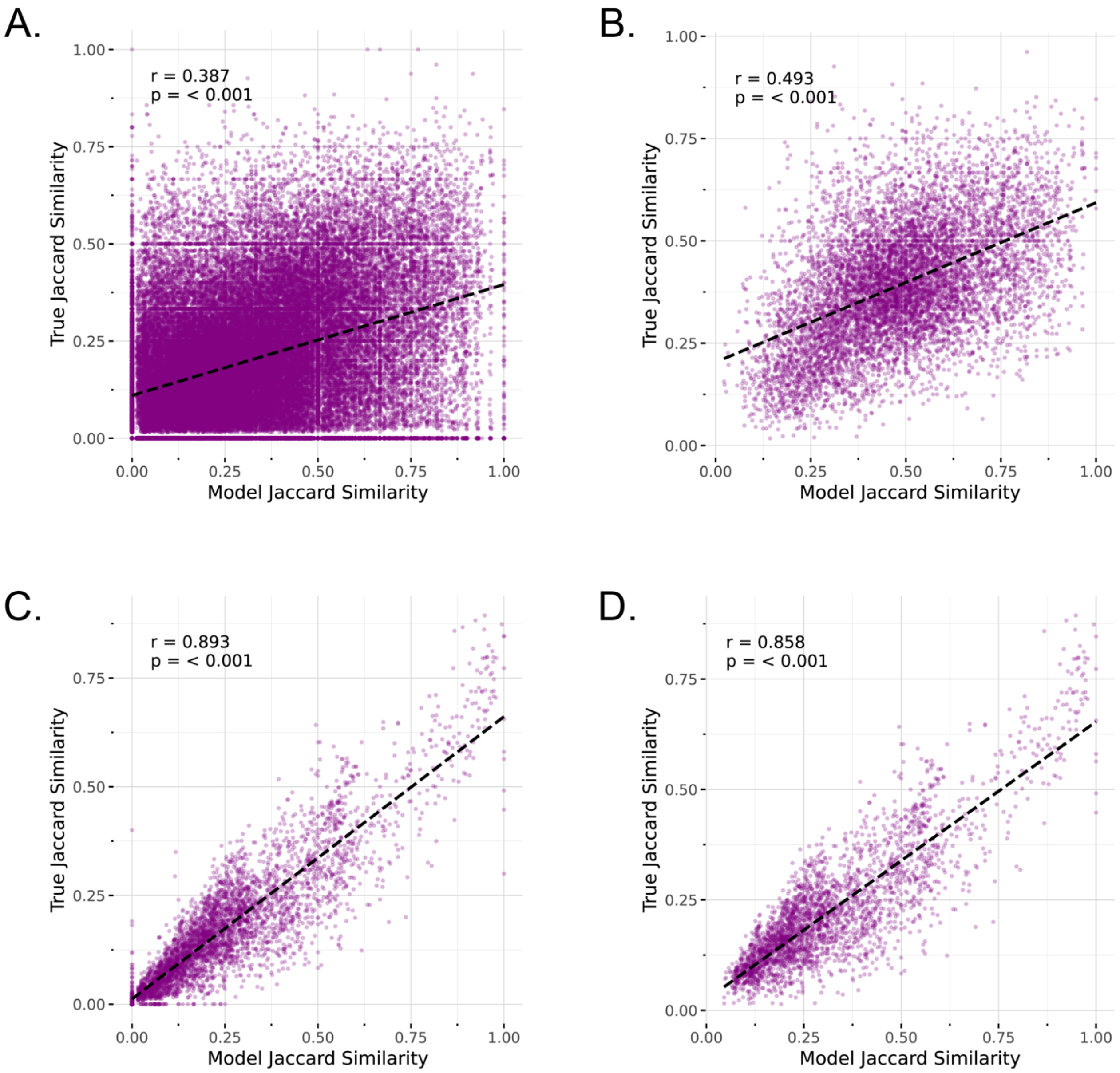

*Supplementary Figure 35. Comparing experimental and predicted interaction profiles.* Jaccard similarities between bacterial strain pairs and phage pairs based on their experimental interaction profiles and compared these to model-predicted interaction similarities across 66,795 strain pairs (A) and 4,371 phage pairs (C), and between broadly susceptible bacterial strains (infected by >= 20 phages) (B) and broadly infectious phages (infects >= 20 strains) (D).

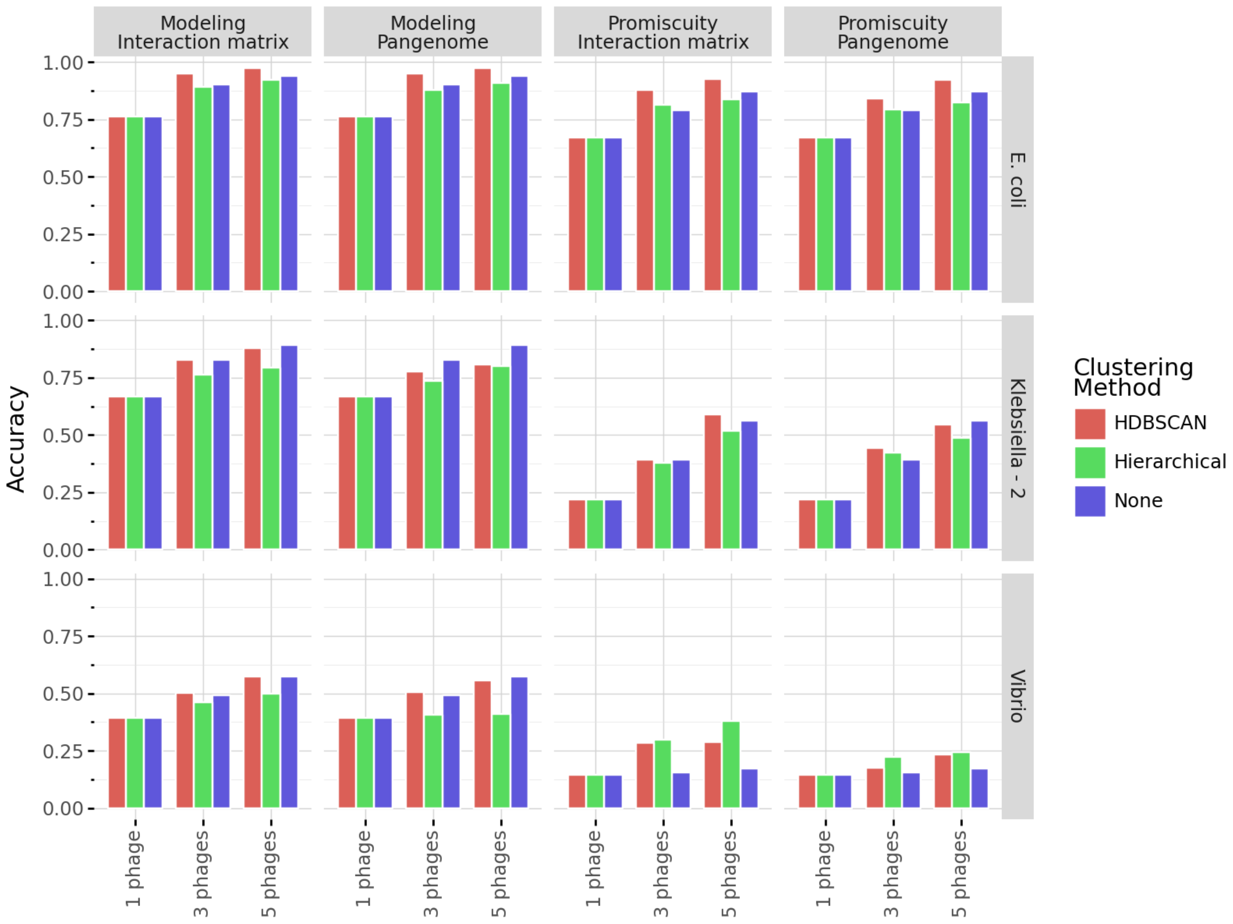

*Supplementary Figure 36. Phage cocktail design workflow clustering comparison.* Proportion of cocktails identifying at least 1 interacting phage across phage clustering algorithms (colors) for both Modeling-based (columns 1-2) and Promiscuity-based (columns 3-4) phage selection. Clustering of phages was based on the interaction matrix (columns 1 and 3) and the phage pangenome (columns 2 and 4).

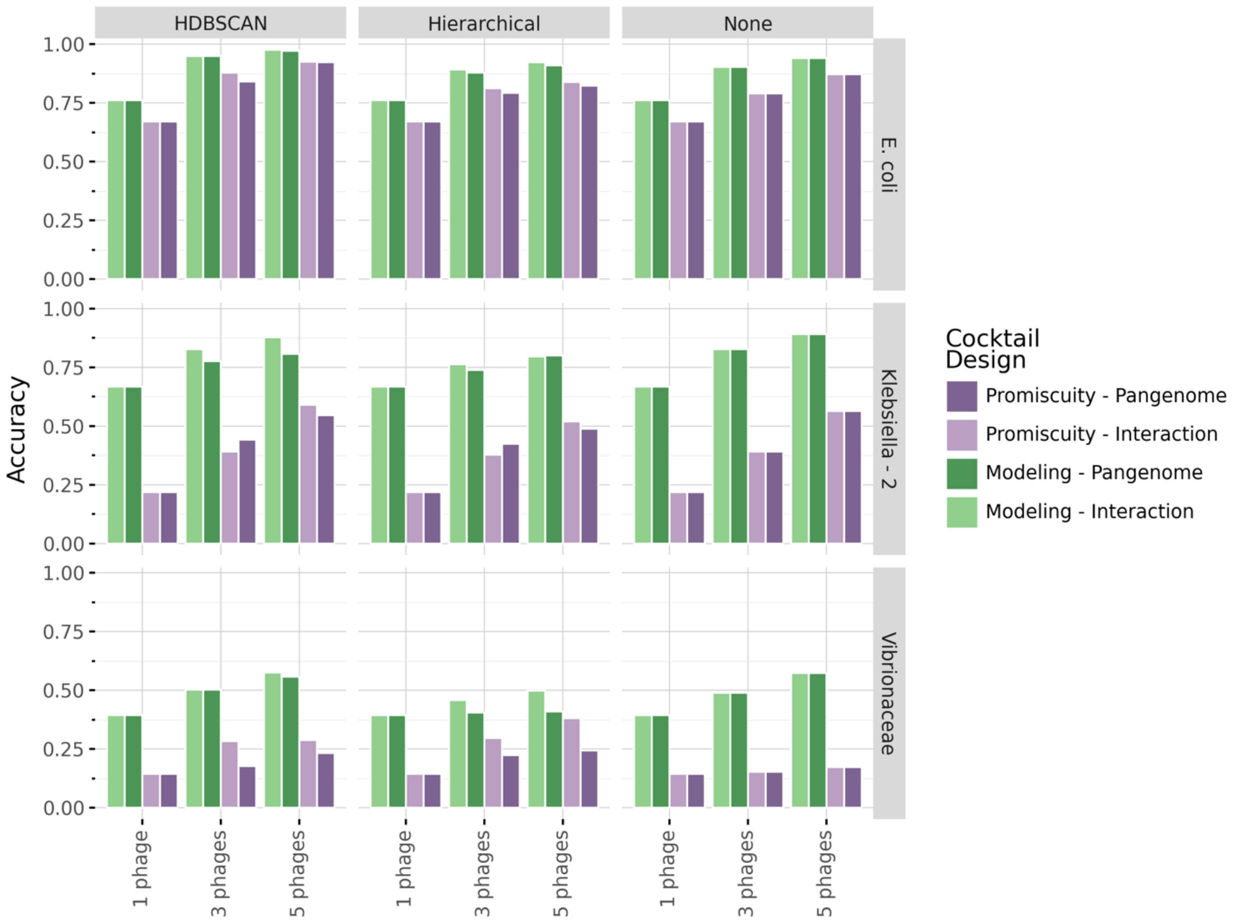

*Supplementary Figure 37. Phage cocktail design workflow ranking strategy comparison.* Proportion of cocktails identifying at least 1 interacting phage across phage clustering algorithms (columns) for both Modeling-based (purple) and Promiscuity-based (green) phage selection. Clustering of phages was based on the interaction matrix (light colors) and the phage pangenome (dark colors).

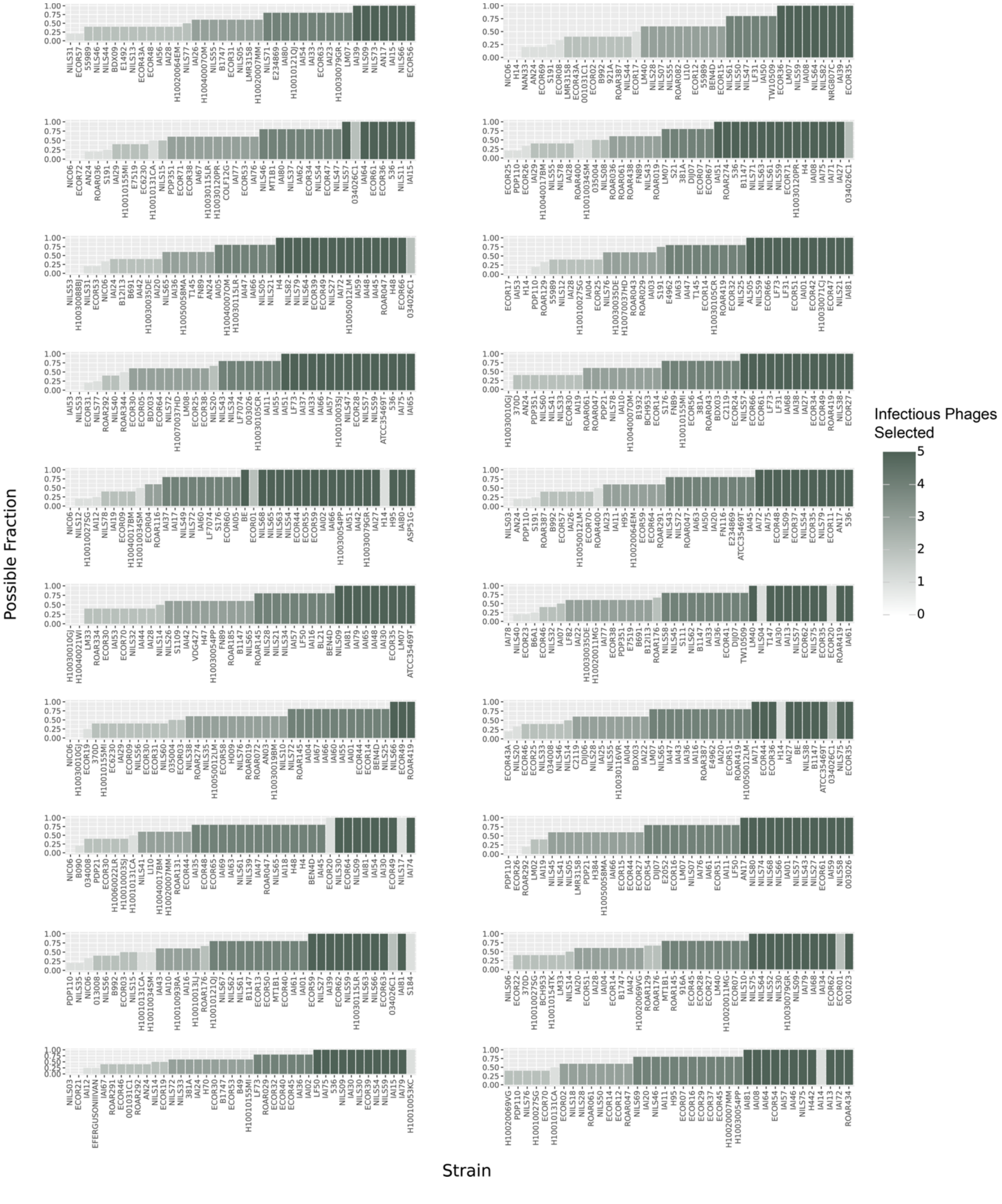

*Supplementary Figure 38. Phage cocktail design in E. coli over 20-fold cross-validation.* Bars show phage cocktail design success rate for all validation strains in 20-fold cross-validation. Bar height represents the fraction of possible phages selected in 5-phage cocktail designs, showing success rate regardless of how many infectious phages were present in the dataset. Bar color represents the total number of infectious phages selected in 5-phage cocktail.

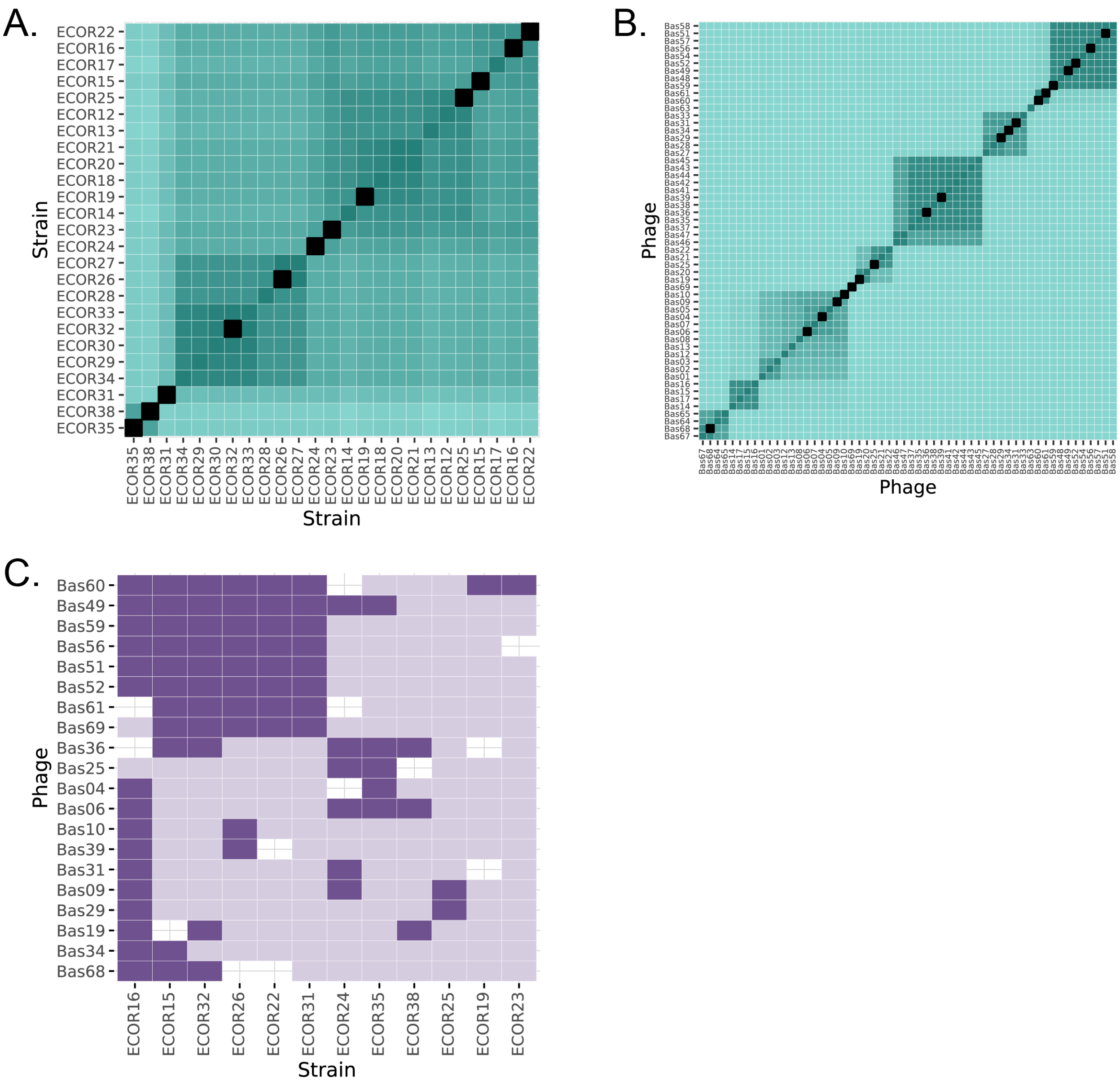

*Supplementary Figure 39. Representative strain and phage selection for EOP experiments.* (A) 12 representative *E. coli* ECOR strains were selected from those in our validation matrix using hierarchical clustering based on a concatenated marker gene tree, after filtering to bacterial strains infected by at least 2 phages. (B) 20 representative BASEL phages were selected from those in our validation matrix using hierarchical clustering based PEQ, after filtering to phages interacting with at least 2 *E. coli* strains. (C) The interaction matrix subset representing the 12 representative *E. coli* ECOR strains and 20 representative BASEL phages. Dark purple represents interactions resulting in infection and light purple indicates interaction that did not result in infection.

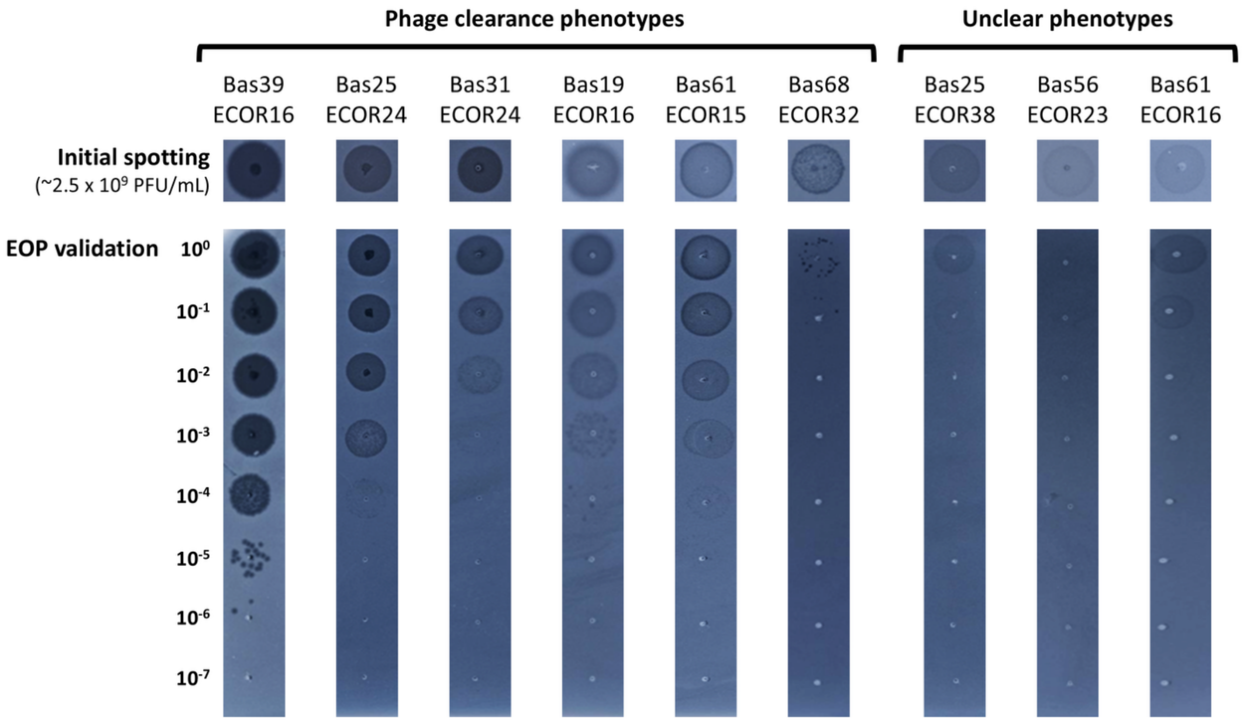

*Supplementary Figure 40. Representative clearance phenotypes in EOP experiments.*

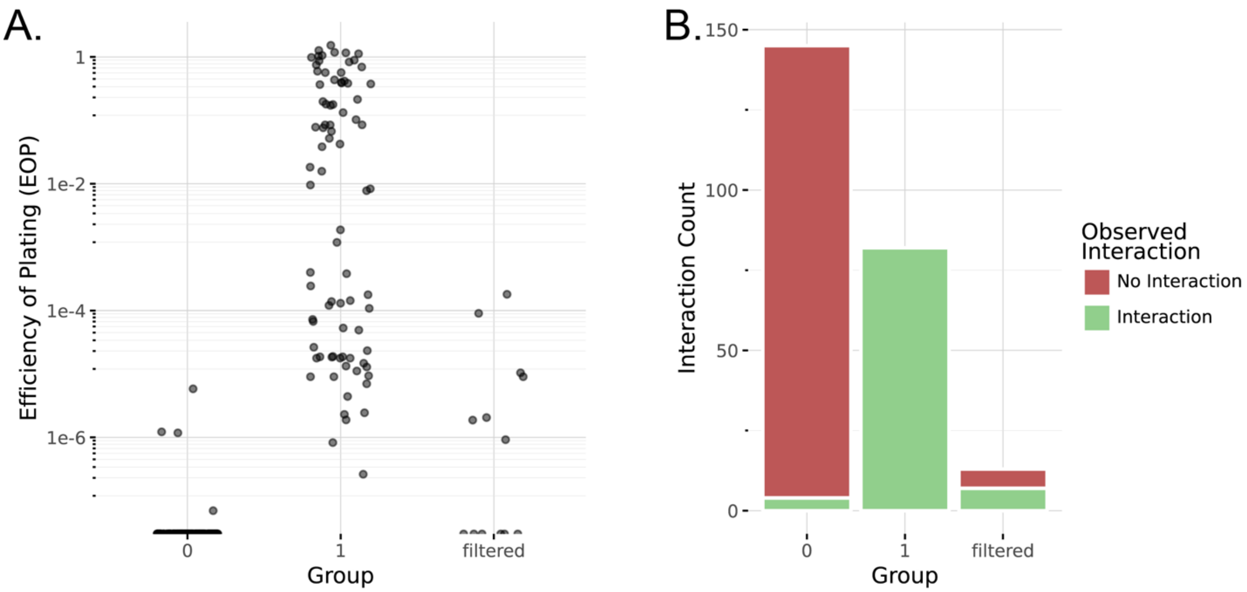

*Supplementary Figure 41. Efficiency of plating (EOP) experiments.* (A) Efficient of plating results for individual phage-host interactions classified as 0 (no interaction) in the original interaction matrix, classified as 1 (interaction), or the filtered based on an unclear phenotype. (B) The number of interactions classified into each group and the observed interaction based on EOP assays (color).

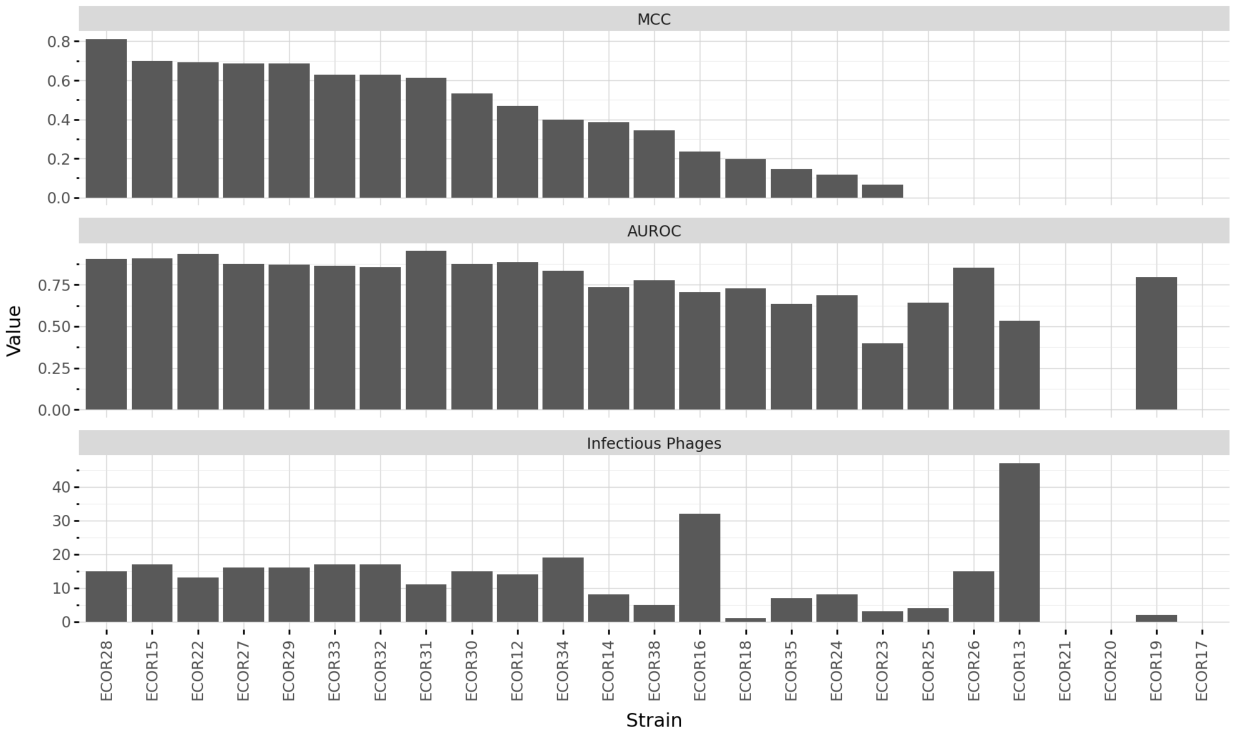

*Supplementary Figure 42. Strain-level predictive performance in validation matrix.* Bars show strain-level predictive performance based on MCC (top) and AUROC (middle), as well as strain susceptibility (total number of infectious phages).

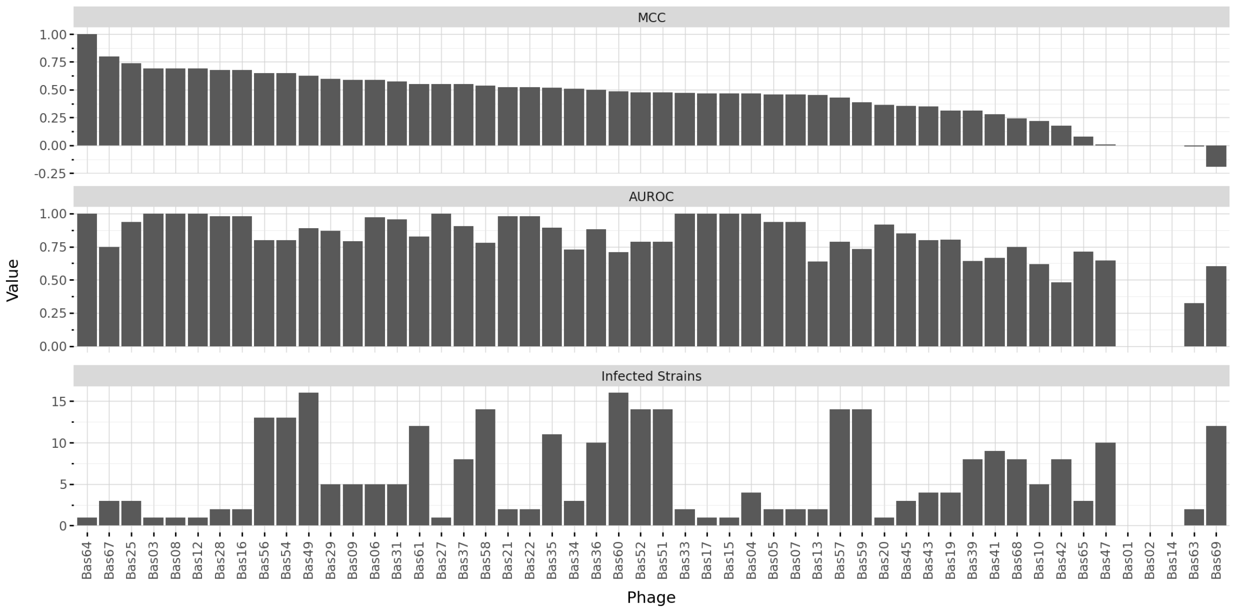

*Supplementary Figure 43. Phage-level predictive performance in validation matrix.* Bars show phage-level predictive performance based on MCC (top) and AUROC (middle), as well as phage infectivity (total number of infected strains).

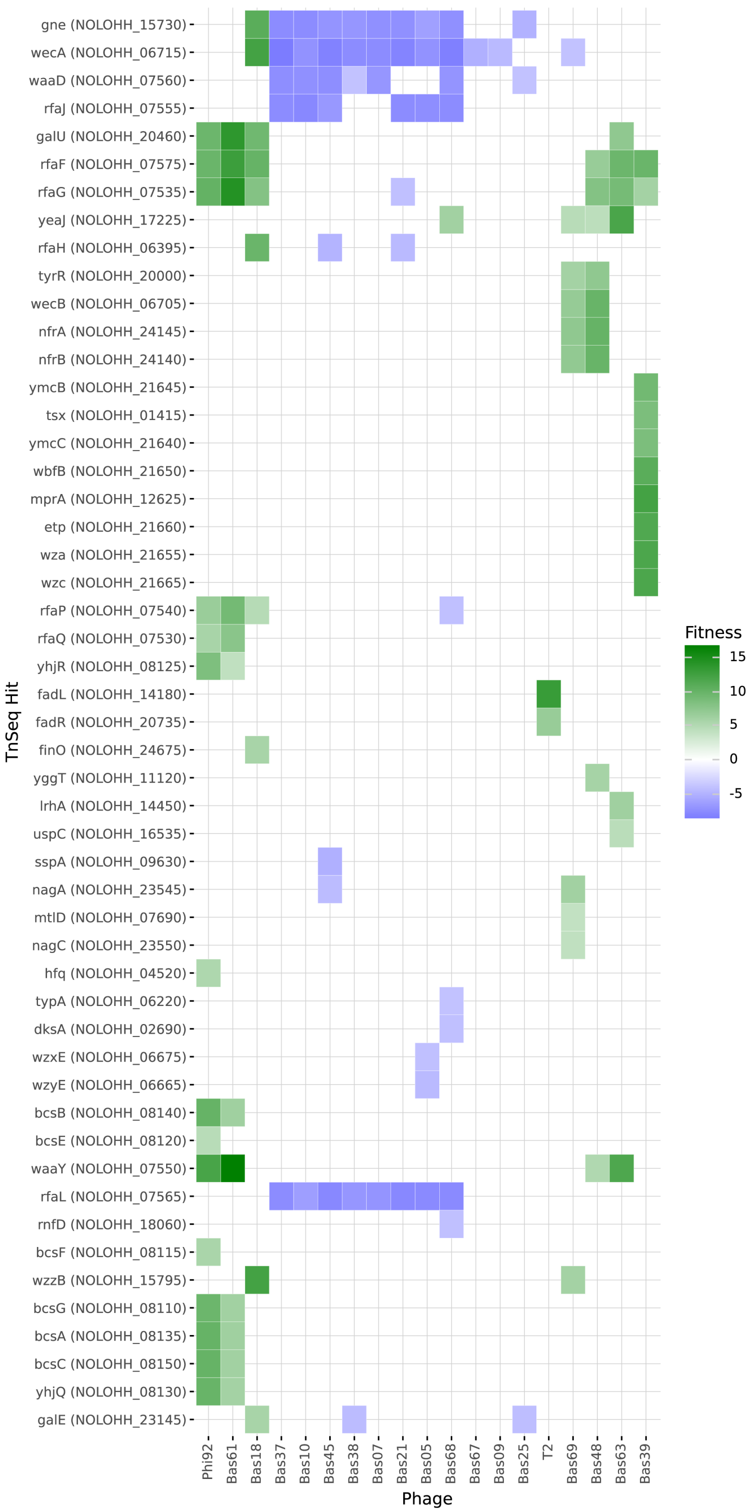

*Supplementary Figure 44. RB-TnSeq hits in E. coli ECOR27 identified across 19 phages.* Heatmap shows *E. coli* ECOR27 genes (X-axis) with strong fitness effects when submitted to one of 19 phages (Y-axis). Color represents fitness scores with positive in green and negative in blue.

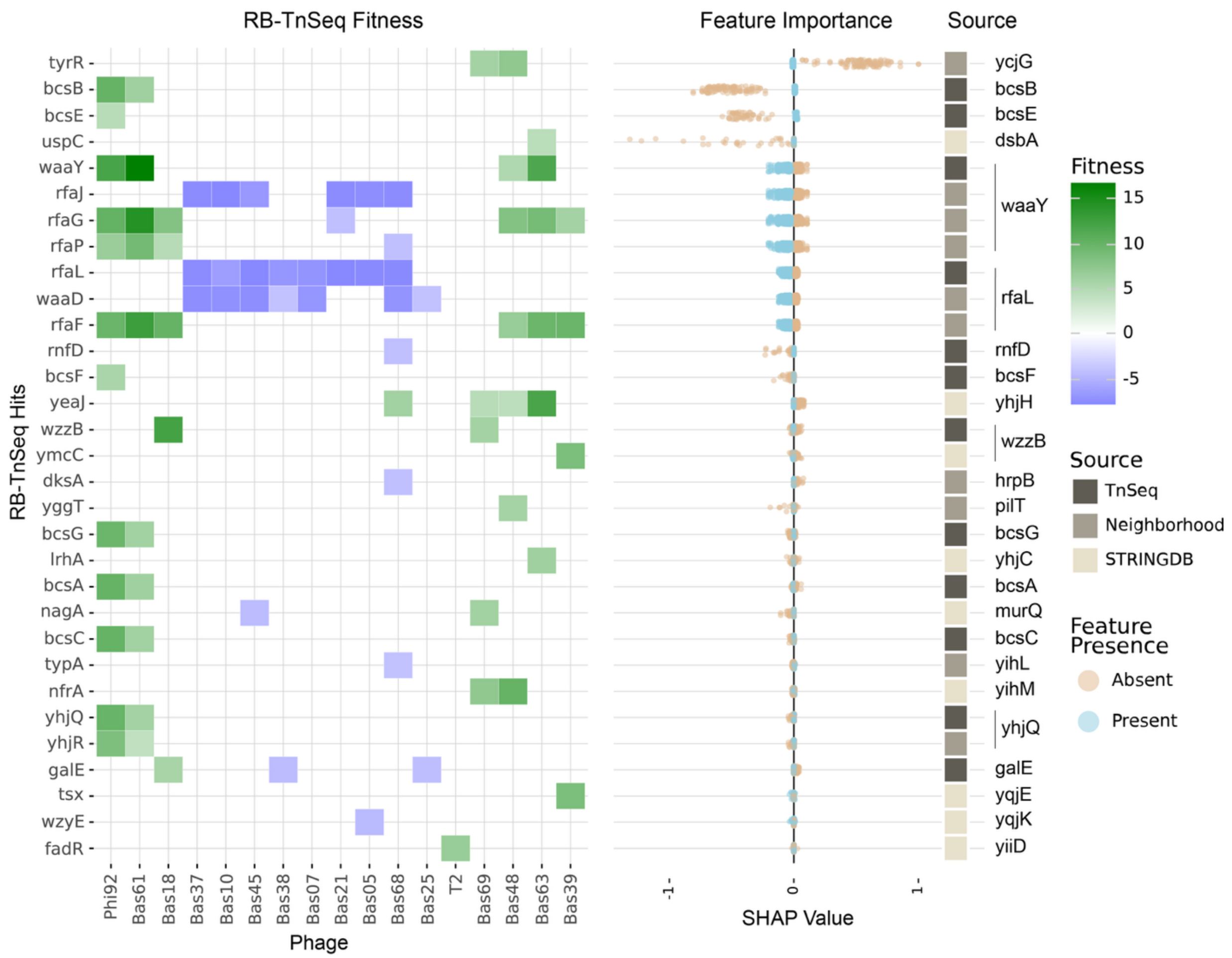

*Supplementary Figure 45. Mapping RB-TnSeq hits from E. coli ECOR27 to predictive features.* The heatmap shows fitness values associated with all RB-TnSeq hits with links predictive features in *E. coli* ECOR27. The X-axis shows tested phages and the Y-axis gene names of RB-TnSeq hits ordered by Shapley Additive exPlanations (SHAP) feature importances. SHAP values indicate whether presence of a given feature (blue) or absence (orange) is associated with increased likelihood of infection (SHAP value > 0) or decreased likelihood of infection (SHAP value < 0). The colored bar shows relationship between RB-TnSeq hits and predictive features, including whether the predictive feature in directly linked to the gene in question (dark brown), linked through a neighborhood analysis (medium brown), or linked through STRING-DB (light brown). Gene names on the far right show the annotation associated with the predictive feature.

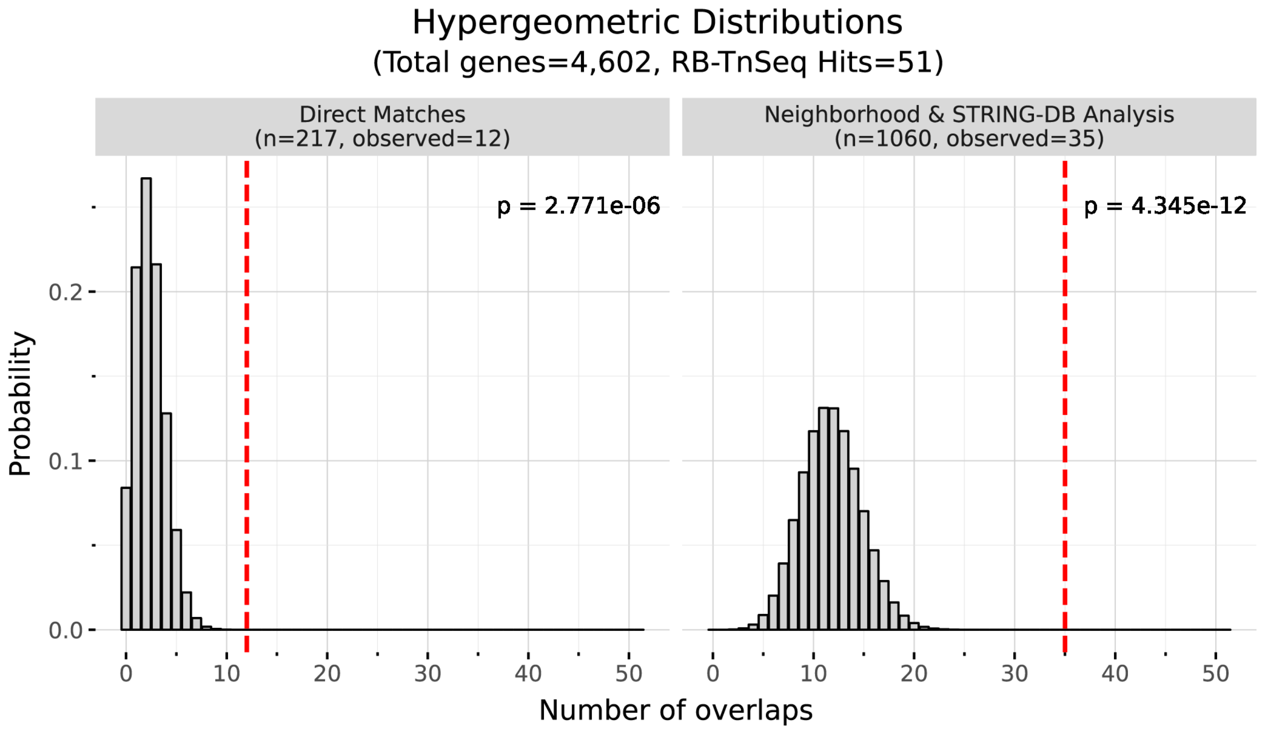

*Supplementary Figure 46. Enrichment of RB-TnSeq hits in predictive features.* Hypergeometric distributions show the expected overlap in predictive and RB-TnSeqs given random sampling. Dashed lines show observed overlap and associated p-values.

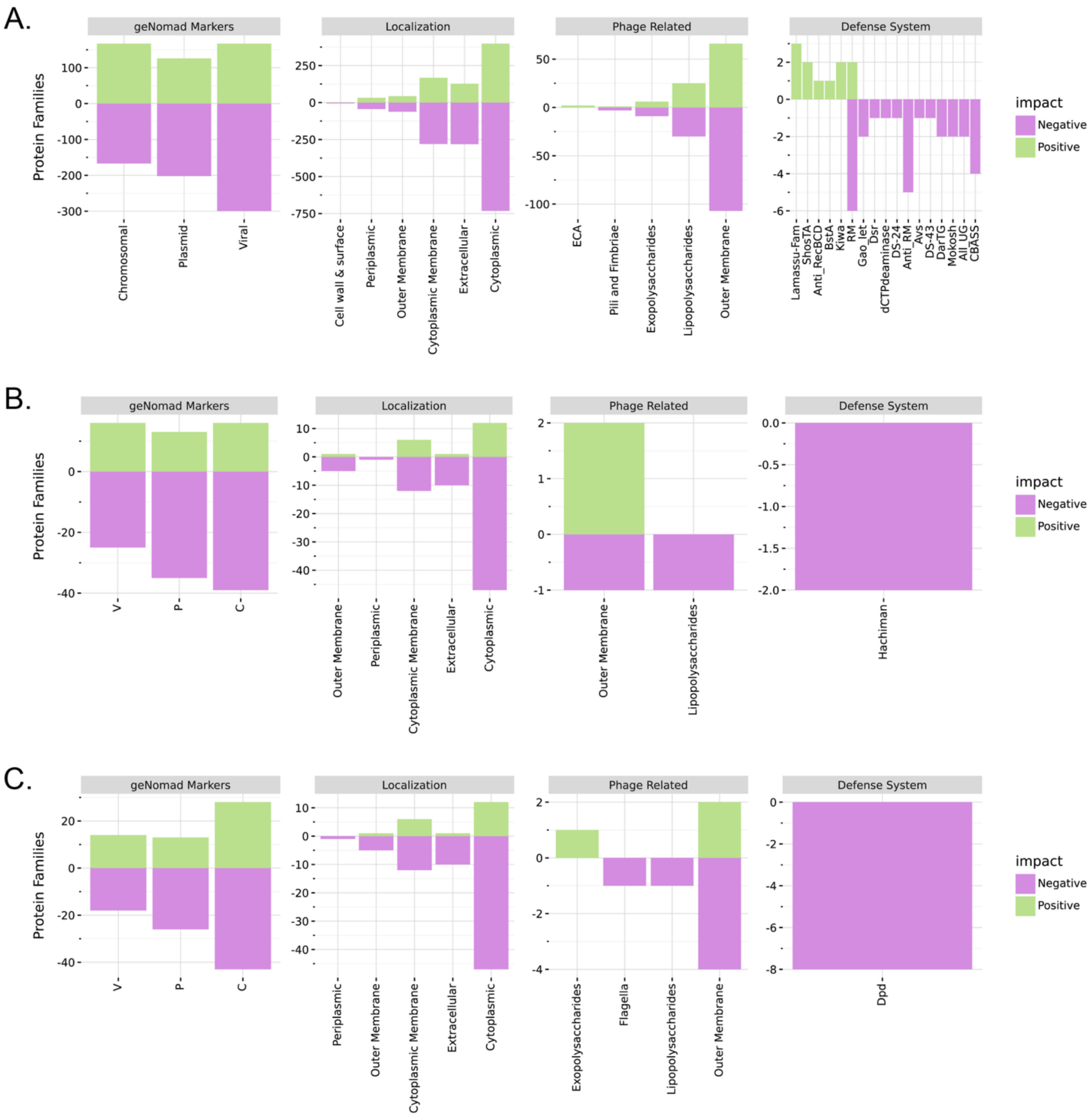

*Supplementary Figure 47. Classification of predictive features based on relevance to known mediators of phage-host interactions.* Proteins associated with predictive features were annotated and classified into having a positive (green) or negative (purple) impact of phage-host interactions across *E. coli* (A), *Klebsiella*-1 (B), and *Klebsiella*-2 (C) datasets.
